# Puf3 contributes to changes in mRNA solubility, translation elongation dynamics at rare arginine codons and loss of protein homeostasis in cells lacking Not4

**DOI:** 10.64898/2025.12.31.695943

**Authors:** Léna Audebert, George E. Allen, Siyu Chen, Olesya O. Panasenko, Susanne Huch, Christine Polte, Zoya Ignatova, Vicent Pelechano, Martine A. Collart

**Affiliations:** Department of Microbiology and Molecular Medicine, Faculty of Medicine and Institute of Genetics and Genomics Geneva, University of Geneva, Switzerland; Department of Infectious Diseases, The First Affiliated Hospital of Zhengzhou University, Zhengzhou, China; SciLifeLab, Department of Microbiology, Tumor and Cell Biology, Karolinska Institutet, Sweden; Biochemistry and Molecular Biology, University of Hamburg, Germany; The BioCode: RNA to Proteins Core Facility, Faculty of Medicine, University of Geneva, Switzerland

**Keywords:** Not4, Puf3, chaperones, Ribosome dwelling, translation elongation, rare arginine codons, tRNA, protein homeostasis, mRNA solubility

## Abstract

The Not proteins of the Ccr4-Not complex regulate translation elongation dynamics, essential for proper folding and assembly of new proteins. In yeast, ribosomes with non-optimal codons in the A-site are enriched within the pool of ribosomes bound by Not4 and Not5. Such ribosomes accumulate in cells lacking Not4 or Not5 that show defects in co-translational assembly and aggregation of new proteins. Recently we observed that depletion of Not1 and Not4 inversely regulate changes in mRNA solubility, correlating with inverse codon-specific changes in A-site ribosome dwelling occupancies (RDOs). Here we describe that mRNAs less soluble upon Not4 depletion are enriched for targets of the RNA-binding protein Puf3. We determine that Puf3 contributes to inverse changes of A-site RDOs upon Not1 and Not4 depletion, in particular at rare arginine codons, and it contributes to changes in mRNA solubility in *not4Δ*. Moreover, deletion of Puf3 suppresses temperature sensitivity and protein aggregation in the *not4Δ* strain, while overexpression of Puf3 is toxic. Puf3 post-translational modifications and the Puf3 interactome are altered in *not4Δ*. Taken together, our results associate alterations in Puf3 post-translational status and function, including contribution to translation elongation dynamics, with *not4Δ* mutant phenotypes.

**GRAPHICAL ABSTRACT:** 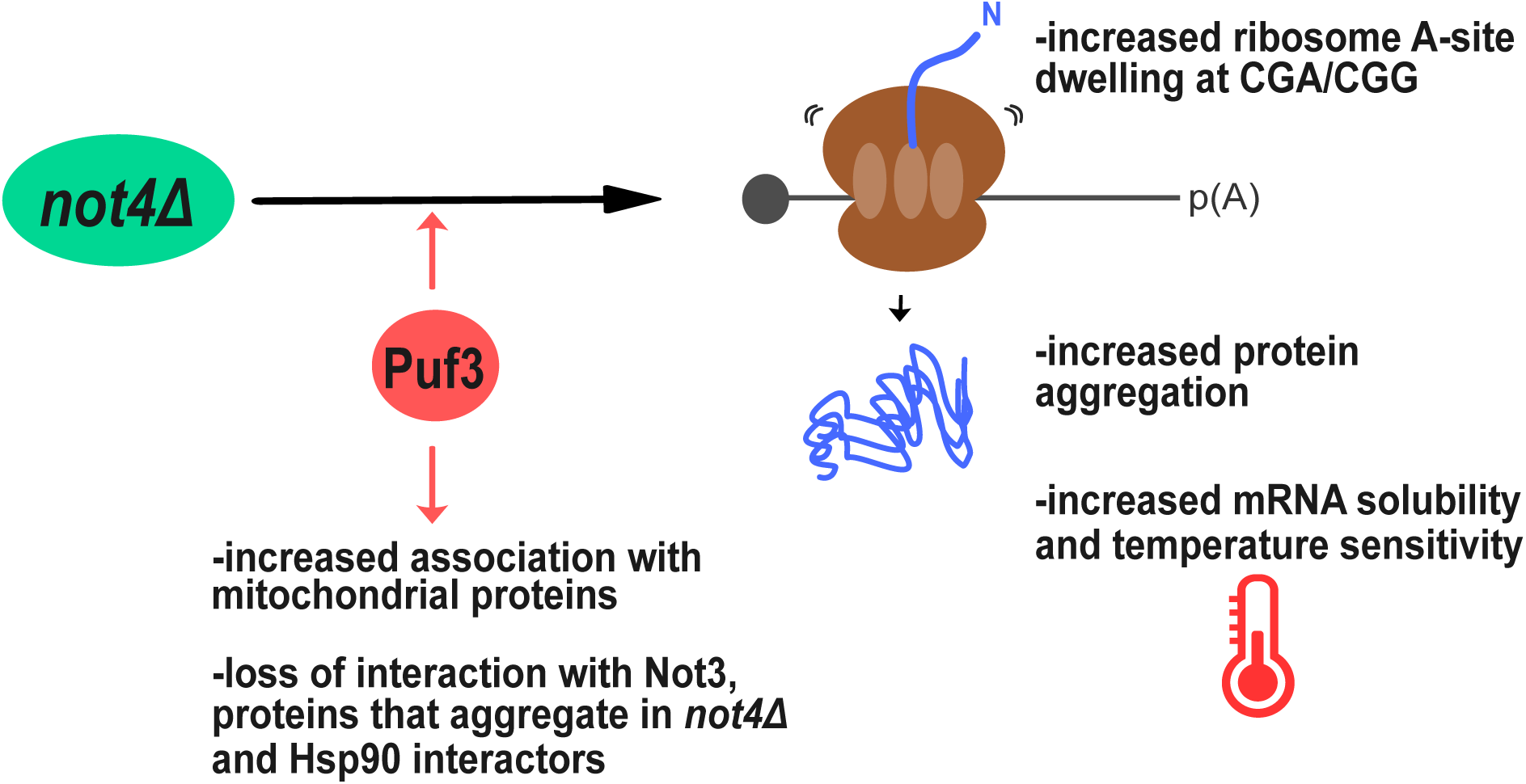

## INTRODUCTION

The Ccr4-Not complex is a global regulator of mRNA metabolism that is conserved in all eukaryotic cells (1). It is built upon a central scaffold protein Not1, onto which other subunits dock organized in functional modules (2–7). It is mostly described to repress gene expression when recruited to its target mRNAs, either by leading to mRNA deadenylation (8–11) or to interact with factors that block translation initiation (12–16). However, accumulating evidences in the last 15 years indicate that it also plays a critical role for translation elongation dynamics (17–24) and facilitates folding and co-translational assembly of proteins. The Ccr4-Not complex can be recruited to its target mRNAs by RNA-binding proteins (25–28), and also by interacting with translating ribosomes (29). Not5, in association with Not4, binds to the E site of post-translocation ribosomes when A sites are empty, a situation more likely to occur when a non-optimal codon is present in the A site. Thereby, it has been proposed that the Ccr4-Not complex monitors the mRNA turnover according to codon optimality (29). We have proposed instead a model whereby binding of the Not proteins to the translating ribosome modulates translation elongation dynamics, to facilitate co-translational events (19). The Not proteins form cytosolic condensates, named assemblysomes, and in human cells, CNOT1 condensates promote colocalization of mRNAs whose nascent peptides interact co-translationally (30). We thus proposed that condensation of Not-associated translating ribosomes regulates translation elongation dynamics (19). Following up on this model, we determined that mRNA solubility in budding yeast varies, with distinct translation elongation dynamics for soluble and insoluble mRNAs (17,19). Ribosomes dwelling at non-optimal A site codons reside more on insoluble mRNAs than on soluble ones (17). We also determined that depletion of 2 different subunits of the complex, the scaffold protein Not1 or the E3 ligase subunit Not4, inversely modified mRNA solubilities (17).

In this work, we zoomed into mRNAs less soluble upon Not4 depletion but more soluble upon Not1 depletion, and determined that they were enriched for targets of the Puf3 pumilio family RNA-binding protein (RBP) (31). Puf3 is a key regulator of mitochondrial mRNAs and acts at the post-transcriptional level through binding to a consensus sequence (UGUA_AUA)(32) in 3’UTRs (33). The mitochondrial targets of Puf3 include proteins involved in the oxidative phosphorylation pathway, mitochondrial protein import, transcription and translation (33–37). Many transcripts regulated by Puf3 are not linked to mitochondria (32,33) and Puf3 binds and regulates transcripts with non-canonical binding sites (35). Nevertheless, motif quality is the major determinant of Puf3-dependent mRNA stability for Puf3 targets *in vivo* (32). Puf3 promotes binding of the Ccr4-Not complex to mRNA isoforms with Puf3 binding sites (18) to accelerate their deadenylation (32,38), in particular during fermentation. Non-fermentable sugar conditions inhibit Puf3’s ability to accelerate mRNA deadenylation within minutes, via phosphorylation of Caf1 by Yak1 that abolishes the interaction between Puf3 and the Ccr4-Not complex (39). During respiratory conditions, Puf3 stabilizes mRNAs that encode mitochondrial proteins (39) and contributes to their translation to support the need for increased mitochondrial biogenesis. It has been proposed that differential phosphorylation of Puf3’s large N-terminal intrinsically disordered region, and Puf3 condensation, are components of a regulatory system that maintains Puf3 as a repressor under fermentation conditions but transforms it into an activator for pre-existing mRNAs upon switch to respiratory conditions (40,41). Puf3 has also been reported to interact with Rpb4, a subunit of RNA polymerase II (RNAPII), that contributes to its’ chromatin and mRNA association, to Puf3 stability and to cytoplasmic Puf3 functions (42). The Ccr4-Not complex’s subunit Not5 also associates with RNAPII via Rpb4 (43), and cytoplasmic localization of Rpb4 in response to stress requires Not5 (19). These previous observations suggest a possible link between Puf3 and the Ccr4-Not complex that connects transcription with the fate of mRNAs in the cytoplasm.

Our current results uncover a codon-specific role of Puf3 in translation elongation dynamics during fermentation and uncover a surprising dominant negative function of Puf3 in *not4Δ*, where it contributes to deregulate translation elongation dynamics and solubility of mRNAs, leading to temperature sensitivity and protein aggregation. They suggest that Not4’s role in protein homeostasis relates at least in part to its role in ensuring appropriate post-translational modifications and interactome of Puf3.

## METHODS

### Yeast strains and growth conditions

Strains used in this manuscript were from either from Euroscarf and derived from BY4739 (*MATα his3 leu2 ura3 lys2*), wild type (MY3672) and *not4Δ* (3417, isogenic except *not4::KANMX4*) used for ribosome profiling previously (19), *puf3Δ* (MY8058, isogenic except *puf3::KANMX4*) and *not4Δpuf3Δ* generated by PCR from MY8058 (leading to MY12455, isogenic to MY3672 except *puf3:KANMX4 not4::NATMX4*). The Not1 (MY13517) and Not4 (MY13771) degron strains have been described (17). *puf3Δ* was created by PCR in the Not1 and Not4 degron strains (leading to MY14441 and MY14401) and as well as in the isogenic wild type TIR-expressing strain (MY13472, leading to MY14416), by PCR. All strains were verified by PCR. The strain expressing Puf3-Taptag was from Euroscarf (SC1249, here called MY8057) and the *NOT4* gene was deleted in this strain by PCR (leading to MY12356). All strains were verified by PCR and are listed in **Supplementary Table S1**.

Not1 and Not4 depletions were obtained by addition of auxin (3-indoleacetic acid, stock solution at 250 mM in EtOH 100%) at 1 mM final for 15 min to exponentially growing cells diluted after an overnight culture in glucose rich medium (YPD), when they reached an OD_600_ of 0.8-1. Equivalent amounts of EtOH 100% were added for the control. All experiments were performed with cells growing in YPD or in selective SC medium for experiments with cells transformed with plasmids.

For the spot growth tests, cells were collected from plates and resuspended in appropriate media and then adjusted to an OD_600_/ml of 0.1-0.2 from which 4 or 5-fold serial dilutions were prepared (40 μl in 160 μl) and 4 or 5 μl of each dilution were spotted on appropriate plates.

Yeast growth media were standard.

### Plasmids and oligos

Plasmids and oligonucleotides used in this study are listed in **Supplementary Table S1**.

For the plasmid expressing the tRNA having the ACG anticodon that is converted to ICG in the mature tRNA, a PCR amplification using genomic DNA from a wild type strain (MY3672) was performed using primers 1419 and 1420. The generated fragment was digested with *Eco*RI and *Not*I together with the pRS426 plasmid. Both insert and vector were ligated using the T4 DNA ligase (NEB) and transformed in *E. coli* competent cells. Transformants were selected on LB ampicillin plates and clones were tested by digestion and confirmed by sequencing. This created plasmid pMAC1452. This plasmid was then digested with *Nco*I and *Not*I together with the pRS416 and further cloning steps were proceeded as described above to create pMAC1467.

For the plasmid expressing the tRNA having the anticodon UCG, the tRNA gene was amplified from DNA of a wild type strain (MY3672) using the oligo pairs 1419-1428 and 1427-1420 to mutate the CGT to CGA. The generated fragment was further amplified using 1419-1420 primers and the amplicon was digested with *EcoR*I and *Not*I. The pRS424 was digested using *EcoR*I and *Not*I. Both fragment and insert were ligated using the T4 DNA ligase and further cloning steps were proceeded as described above to create pMAC1457. This plasmid was then digested with *Nco*I and *Not*I together with the pRS416 and further cloning steps were proceeded as described above to create pMAC1466.

### Post alkaline lysis for analysis of total protein levels

Exponentially growing cells expressing Puf3-Tap-tag were pelleted and pelleted cells were resuspended in 0.1 M NaOH and incubated for 10 min at room temperature (RT). After a quick spin in a microfuge, the cell pellet was resuspended in 2 X sample buffer followed by standard SDS-PAGE and western blotting with antibodies to the protein A (Sigma, #P1291) and Gapdh (Thermo Fisher Scientific, #MA5-15738) for loading control.

### Test of protein integrity for affinity purification

Exponentially growing yeast cells expressing Puf3-Tap-tag were harvested at 1 OD_600_/ml (50 ml of cells) for 15 min at 3500 rpm at room temperature and kept at - 20°C. Before breaking the cells, pelleted cells were resuspended with 400 μL of lysis buffer (20 mM Hepes pH 7.4, 100 mM kOAc, 1 X VRC, protease inhibitor (Roche complete EDTA free)). Cells were broken using glass beads by vortexing for 10 min at 4°C. To pellet glass beads and cellular debris, samples were centrifugated for 20 minutes at 13000 rpm at 4°C then clear lysates were transferred into new tubes. Protein concentration was determined by a Bradford assay, and all samples were adjusted to the same concentration. 100 μL of adjusted samples were mixed with 50 μL 4 X sample buffer. Serial dilutions starting with 10 μL containing equal amounts of total protein were separated by SDS-PAGE and analyzed by western blot using antibodies to the protein A (Sigma, #P1291) and Gapdh (Thermo Fisher Scientific, #MA5-15738) for loading control.

### Preparation of total and soluble RNA

Total RNA was prepared either by the hot acid phenol method (44) or cells were prepared and lysed as for polysome fractionation (22) and total RNA was prepared from the lysate. Briefly, for total RNA, cell pellets from 30 ml of exponentially growing yeast were resuspended with 400 μl of TES buffer (10 mM Tris HCl pH 7.5, 10 mM EDTA and 0.5 % SDS) to which 400 μl of acid phenol were added. After vortexing and incubation for 10 min at 65°C, total RNA was extracted. For soluble RNA, cell pellets from 30 ml of exponentially growing yeast were resuspended in 400 μl of lysis buffer (20 mM Hepes, 20 mM KCl, 10 mM MgCl_2_ 1% triton, 1 mM PMSF, 1 mM DTT, protease inhibitor (Roche complete EDTA free) to which 200 μl of glass beads were added. Cells were vortexed at 4°C for 15 min and spun at 15000 g for 1 min. The supernatant was transferred to a new tube and spun 20 min at 4°C at 15000 g. The supernatant was combined with 400 μl of acid phenol for further RNA extraction as for the total RNA pool.

WT (MY3672) and *not4Δ* (MY3417) total and soluble RNAs were prepared and analyzed by RNA-Seq previously and soluble RNA from WT (MY3672) and *not4Δ* (MY3417) were also analyzed previously by Ribo-Seq (17,19). In this study, soluble RNAs from WT (MY3672), *puf3Δ* (MY8058) and *not4Δpuf3Δ* (MY12455) were prepared and analyzed similarly by RNA-Seq.

### Soluble and total RNA-Seq

The total and soluble RNA was prepared from 50 ml of exponentially growing cells in YPD. Libraries were generated as reported (17). In brief, 15 μg RNA, containing 5% total RNA from *Schizosaccharomyces pombe* as spike-in, was used. The samples were subject to random fragmentation prior to making the HT-5PSeq Libraries.

For HT-5PSeq libraries: 7.5 μg RNA was ligated overnight at 16°C to r5P_RNA_MPX oligo (CrArCrGrArCrGrCrUrCrUrUrCrCrGrArUrCrU rXrXrXrXrXrX rNrNrNrNrNrNrNrN) carrying a sample barcode (rX) and unique molecular identifiers (rN). Ligase was deactivated using 5mM EDTA and heated at 65°C for 10 minutes (up to X individual barcoded RNA ligations were pooled) and subsequently purified using 1.8x volumes of RNAClean XP beads (Beckman Coulter). Ligated RNA was then reverse transcribed using random hexamer (5Pseq-RT, GTGACTGGAGTTCAGACGTGTGCTCTTCCGATCTNNNNNN, 20 μM) and oligo-dT (5Pseq-dT, GTGACTGGAGTTCAGACGTGTGCTCTTCCGATCTTTTTTTTTTT at 0.05 μM) oligos to prime. After, the remaining RNA was degraded using NaOH. Ribosomal RNA was removed using previously described rRNA DNA oligo depletion mixes, following a duplex-specific nuclease (DSN, Evrogen) digestion. rRNA depleted cDNA was amplified by PCR (17 cycles) and the final product was enriched for fragments with the range of 300-500 nucleotides using Ampure XP. Size selected HT-5P libraries were quantified by fluorescence (Qubit, Thermo Fisher Scientific), the size was estimated using an Agilent Bioanalyzer and sequenced using a NextSeq500 Illumina sequencer (75 cycles High output kit).

Sequencing files were demultiplexed using bcl2fastq v2.20.0.422 (one mismatch, minimum length 35 nucleotides), and adapters were trimmed using cutadapt 2.3. at default settings, allowing one mismatch and a minimum read length of 35 nucleotides. In addition to standard illumine dual index (i5, i7), the inline sample and UMI barcode was analyzed using Umitools. Reads were mapped to the concatenated genome of *S. cerevisiae* (R64-1–1) and *S. pombe* (ASM294v2) using STAR.

Second read enables to split reads between oligo-dT or random primer. That information was not used in the current analysis. CDS positions were defined with the Ensembl gff version 94 for *S. cerevisiae* (R64-1–1). Counts in *S. cerevisiae* were calculated by aggregating RNA-Seq reads and 5′P-Seq 5′-ends, overlapping CDS positions.

### Solubility

We define this as the log fold change produced by DESeq2, dividing RNA-Seq counts for the soluble RNA fraction in a given sample by the corresponding counts for the total RNA fraction of the same sample. The solubility of deletion and depletion strains is then corrected to wild type basal solubility.

### Ribosome profiling (Ribo-Seq)

Soluble RNAs from WT (MY13472), *puf3Δ* (MY14373), Not1 degron (MY13517), Not4 degron (MY13771) Not1 degron with *puf3Δ* ( MY14441) and Not4 degron with *puf3Δ* (MY 14401) were analyzed by Ribo-Seq in biological triplicates with some modifications compared to the previous Ribo-Seq analysis (30), at the BioCode: RNA to Proteins Core Facility of the Faculty of Medicine (University of Geneva).

For cell preparation, exponentially growing cells were collected at an OD_600_/ml of 0.8-1 and before auxin treatment, an aliquot of cells was taken to verify the protein levels prior to depletion. Auxin was added at a final concentration of 1 mM for 15 min at 30°C under agitation, an aliquot of cells was taken to check for efficient protein depletion. Before cell collection by centrifugation, cycloheximide was added in the media at 0.1 mg/ml final concentration and cells were spun for 3 min at 4600 g at 4°C. Pelleted cells were resuspended in 30 ml of Polysome Buffer (PB: 20 mM Tris, pH 7.4, 150 mM NaCl, 5 mM MgCl_2_, 1 mM DTT, 0.1 mg/ml cycloheximide). Cells were spun for 3 min at 4600 g at 4°C. The cell pellet was resuspended in fresh PB and transferred to a new tube prior to spinning. The pellets were kept frozen at -80°C.

The degron protein levels were checked by post-alkaline lysis, standard SDS-PAGE and western blotting with antibodies to myc, from cells collected before and after auxin addition. Post-alkaline lysis involves incubating yeast cells in 0.1 M NaOH for 5 min at RT, pelleting, and suspending the pellet in 2 X sample buffer. After confirmation of protein depletion, the frozen cell pellet was processed further for Ribo-Seq and soluble RNA extraction from extracts prepared as described (30) before.

Soluble RNAs were isolated from the same extracts that were used to obtain RPFs. 3 volumes of QIAzol® (Qiagen, #79306) were added to 80 μl of extracts, mixed and proceed to RNA purification with Direct-Zol RNA Mini Prep Plus kit (Zymo Research, #R2070).

RNAs were sent to the iGE3 Genomic Platform (University of Geneva) for stranded mRNA libraries preparation and sequencing. Quality control of RNAs was check by Bioanalyzer. RNA with quality higher than RIN: 9.80 were used for libraries preparation.

Libraries were sequenced on an Illumina NovaSeq 6000, single-reads, 1x 100 bp.

To obtain ribosome footprints 100 µl of total extracts, containing 500 µg of total RNA, were treated with RNAse I (Ambion, 100 units/µl #AM2294) in concentration 1600 U of RNAse I per 1 mg of total RNA, for 1 h at 20°C with slow agitation. 10 ml SUPERaseIn RNase inhibitor (Ambion, #AM2694) was added to stop nuclease digestion. Monosomes were isolated using by size-exclusion chromatography using MicroSpin S-400 HR columns (Cytiva, #27514001). 3 volumes of QIAzol® (Qiagen, #79306) were added to the S-400 eluate, mixed and proceed to RNA purification with Direct-Zol RNA Mini Prep Plus kit (Zymo Research, #R2070).

For Ribo-Seq libraries preparation, RPFs (25-34 nt) were size-selected by electrophoresis using a 15% TBE-Urea polyacrylamide gel electrophoresis (PAGE) and two RNA markers, 25-mer (5’ AUGUACACGGAGUCGAGCACCCGCA 3’) and 34-mer (5’AUGUACACGGAGUCGAGCACCCG CAACGCGAAUG 3’). After dephosphorylation with T4 Polynucleotide Kinase (NEB, #M0201S) the adapter Linker-1 (5’ rAppCTGTAGGCACCATCAAT/3ddC/ 3’) was ligated to the 3′ end of the RPF using T4 RNA Ligase 2 (NEB, #M0242S). The ligated products were purified using 10% TBE-Urea PAGE. Ribosomal RNAs were subtracted using RiboCop rRNA Depletion Kit V2 Yeast (Lexogen, #190). The adapter Linker-1 was used for priming reverse transcription with the primer Ni-Ni-9 (5’AGATCGGAAGAGCGTCGTGTAGGGAAAGAGTGTAGATCTCGGTGGTCGC_5_CA CTCA_5_TTC AGACGTGTGCTCTTCCGATCTATTGATGGTGCCTACAG 3’) containing 2 internal 18-atom hexa-ethylene glycol (HEG) chains, using ProtoScript® II Reverse Transcriptase (NEB, #M0368S). RT products were purified using 10% TBE-Urea PAGE. The cDNA was circularized with CircLigase™ II ssDNA Ligase (Epicentre, #CL9021K). The final libraries were generated by PCR using forward index primers.

Amplified libraries were purified using 8% TBE-PAGE and analyzed with TapeStation High Sensitivity. Libraries were sequenced on an Illumina NovaSeq 6000, single-reads, 1×100 bp.

For *not4Δ* and associated WT Ribo-Seq samples, data was analyzed as previously described (19).

Published Ribo-Seq data for *puf3Δ* was downloaded to be re-analyzed from (45) and new Ribo-Seq data generated for WT, *Not1d*, *Not4d*, *puf3Δ*, *Not1d puf3Δ* and *Not4d puf3Δ*, fastq files were adaptor stripped using cutadapt (46). Only trimmed reads were retained, with a minimum length of 15 and a quality cutoff of 2 (parameters: -a CTGTAGGCACCAT –trimmed-only –minimum-length = 15 –quality-cutoff = 2). Histograms were produced of ribosome footprint (RPF) lengths that were very homogeneous with highest reads between 26 and 29 that were kept for the analysis. Reads were mapped, using default parameters, with HISAT2 (47) to R64-1-1, using Ensembl release 84 gtf for transcript definitions. UTR definitions were taken from the *Saccharomyces* Genome Database and a standard region of 100 bp was used where a gene’s UTR was not defined. A full set of 6692 CDSs were established for R64-1-1 Ensembl release 84 and extended by the same UTR sequences defined above. Definitions of rRNA loci were taken from UCSC Repeat Masker tracks for R64-1-1 and all reads mapping to these regions were removed. The filtered reads were then mapped to this transcriptome with Bowtie2 (48) using default parameters. Only unique alignments to transcripts were retained. For all downstream analyses, dubious ORFs were filtered to leave 5929 transcripts. The A/P site position of each read was predicted by riboWaltz (49) and aggregated over all transcripts. Differential expression was performed using DESeq2 (50).

Where RDOs are reported for E, P, A sites, a count on a given codon is taken to be the number of E, P, A sites covering it, respectfully. These values are divided by the mean codon coverage for every respective CDS included in the analysis and averaged for each codon type. Transcripts used in the analysis of RDO were restricted to those with RPKMs greater than 1 across all samples, leaving 4060 transcripts. The log_2_ ratio of deletions or depletions for RDOs were taken relative to wild type for each codon. When reporting a pairwise correlation, RDO values for each codon were calculated for each sample and correlated across all 61 codons. Optimal codons were defined as the 15 codons with the highest tRNA adaptation index (tAI) and non-optimal codons as the 15 with the lowest tAI (51).

### Quantification and statistical analysis

Reported correlations were the Pearson’s product-moment correlation coefficient and were used as test statistics to generate the associated p-value by t-test. All correlations and correlation tests were performed on groups of at least 30 in size. Enrichment of gene sets is defined via FDR after Benjamini-Hochberg adjustment from p-values generated using a hypergeometric test. All t-tests were performed on sample sizes of 30 or higher, where the Central LimitTtheorem applies regarding the normality assumption. We use Welch’s t test in all cases, rather than the Student’s t-test, resulting in a more conservative p-value, which is more reliable where variances and sample sizes are unequal.

### tRNA microarrays

To determine the fraction of charged tRNAs, we followed the procedure described in (52). Briefly, total RNA was isolated in mild acidic conditions (pH 4.5) which preserves the aminoacyl-group. Each sample was split into two aliquots and one was oxidized with periodate to which the charged tRNAs remain intact. Following subsequent deacylation (100 mM Tris (pH 9.0) at 37°C for 45 min) it was hybridized to a Cy3-labeled RNA/DNA stem-loop oligonucleotide. The second aliquot was deacylated to obtain the total tRNA and hybridized to Atto647-labeled RNA/DNA stem-loop oligonucleotide. Both labeled aliquots were mixed in equimolar amounts, spiked in with standards and analyzed on the same tRNA microarrays. The slides for the microarrays contain tRNA probes covering the full-length sequence of cytoplasmic tRNA species as described previously (19).

For tRNA abundance, the total RNA was isolated at alkaline pH to simultaneously deacylate all tRNAs. tRNAs labeled with Cy3-labeled RNA/DNA stem-loop oligonucleotide were hybridized on the same microarray with tRNAs isolated from wild-type strain labeled with Att647-labeled RNA/DNA stem-loop oligonucleotide. The arrays were normalized to spike-in standards, processed and quantified with in-house python scripts as described (52).

### Preparation of soluble extracts, insoluble fractions and aggregated proteins

Protein aggregates were isolated as described previously (22,53). 50 OD_600_ of cells growing exponentially were collected by centrifugation at room temperature (4000 rpm, 5 min). Cell pellets were washed with H_2_O and transferred to Eppendorf tubes. The pellets were then resuspended in 0.5 ml lysis buffer (20 mM NaPi pH 6.8, 1 mM EDTA, 0.1% Tween 20, 10 mM DTT, 1 mM PMSF, protease inhibitor (Roche complete EDTA free), 3 mg/ml Zymolase T20) at 30°C for 20 min. Samples were put on ice and sonicated (10 counts, 3 times, 40% intensity). Samples were spun for 20 min at 200 g at 4°C. The supernatant was kept, and a Bradford assay was done to determine protein concentration. All samples were adjusted to the same protein concentration. An aliquot of the total extract was kept for total extract analysis (mixed with 2 X sample buffer) and the remaining of the samples was spun for 20 min at 16000 g at 4°C. Pellets were washed in 0.5 ml of WB-NP (20mM NaPi pH 6.8, 1mM PMSF, protease inhibitor (Roche complete EDTA free) and 2% of NP-40) and sonicated (10 counts 1 time 40% of intensity). This was repeated once and pellets were resuspended in 0.5 ml of WB (20 mM NaPi pH 6.8, 1mM PMSF, protease inhibitor (Roche complete EDTA free)) and sonicated (10 counts, 1 time, 40% intensity). Samples were spun for 20 min at 16000 g at 4°C and pellets were resuspended in 2 X sample buffer). Samples were boiled at 100°C for 5 min and loaded on gradient SDS gels (4-12% NuPAGE™ bis-acrylamide, Thermo Fisher Scientific) for silver nitrate analysis.

### Mass spectrometry characterization of aggregated proteins

The quantitative proteomic analysis was performed by the LC-MS at the Proteomics Core Facility of the Faculty of Medicine (University of Geneva).

Samples of the protein aggregates were prepared as described above, except that an extra washing in WB was done to eliminate traces of detergent, and the aggregate samples were not resuspended in 2X sample buffer but kept as dried pellets at -20°C. Cell pellets were resuspended in 50 μL of 0.1 % RapiGest Surfactant (Waters) in 50 mM Ammonium Bicarbonate (AB). Samples were heated for 5 min at 100°C and sonicated (6 x 30 sec) at 70% amplitude and 0.5 pulse. Samples were kept 30 sec on ice between each cycle of sonication. Samples were centrifuged for 10 min at 14000 x g. Sample volume was adjusted to 100 μL with 0.1 % RapiGest in 50 mM AB. 2 μL of dithioerythritol (DTE) 50 mM were added and the reduction was carried out at 37°C for 1h. Alkylation was performed by adding 2 μL of iodoacetamide 400 mM during 1 h at room temperature in the dark. Overnight digestion was performed at 37 °C with 8 μL of freshly prepared trypsin (Promega: 0.1 μg/μL in 50 mM AB). To remove RapiGest, samples were acidified with TFA, heated at 37°C for 45 min and centrifuged 10 min at 17000 x g. Supernatants were then desalted with a C18 microspin column (Harvard Apparatus, Holliston, MA, USA) according to manufacturer’s instructions, completely dried under speed-vacuum and stored at -20°C.

### Puf3 purification and mass spectrometry

Exponentially growing yeast cells were harvested at 1 OD_600_/ml (1.8L of cells) for 15 min at 3500 rpm at room temperature and kept at -20°C. Before breaking the cells, pelleted cells were appropriately mixed together with 5 ml of lysis buffer (20 mM Hepes pH 7.4, 100 mM kOAc, 1 X VRC, protease inhibitor (Roche complete EDTA free). Cells were broken using glass beads by vortexing for 10 min at 4°C. To pellet glass beads and cellular debris, samples were centrifugated for 20 minutes at 13000 rpm, 4°C and clear lysates were transferred into new tubes.

IgG coupled Superparamagnetic Beads were prepared accordingly to (54). 20 μL of IgG coupled beads per sample were washed 4 times in the extraction buffer (20mM Hepes pH 7.4, 100 mM kOAc,0,5% triton X-100). Beads were resuspended in 100 μL of extraction buffer (per sample) and added to the clear lysates. Samples were put on rotation at 4°C for 2h. After binding, beads were washed 5 times in the extraction buffer. 1 extra washing was performed in the extraction buffer without triton X-100. Elution was performed by adding 50 μL of SDS elution buffer (10 mM Tris-HCl, 1 mM EDTA, 2% SDS) for 10 min at 65°C.

5 μL of the elution was taken to perform a western blot with an antibody against the protein A (Sigma, #P1291). And the remaining of the sample was precipitated with methanol-chloroform accordingly to (54). Dry pellets were conserved at -20°C before proceeding to mass spectrometry.

Sample preparation: Samples were prepared using iST kits (Preomics) according to the manufacturer’s instruction. Briefly, pellets were re-suspended in 50 μl of provided lysis buffer and proteins were denatured, reduced and alkylated during 10 min at 60°C. The resulting slurries (beads and lysis buffer) were transferred to dedicated cartridges and proteins were digested with a Trypsin/LysC mix for 2 hours at 37°C. After two cartridge washes, peptides were eluted with 2 x 100 μL of provided elution buffer. Samples were finally completely dried under speed vacuum and stored at -20°C.

LC-ESI-MSMS: Samples were dissolved in 20 μl of loading buffer (5% CH3CN, 0.1% FA) and 4 μl were injected on a column. LC-ESI-MSMS was performed on a Q-Exactive HF Hybrid Quadrupole-Orbitrap Mass Spectrometer (Thermo Fisher Scientific) equipped with an Easy nLC 1000 liquid chromatography system (Thermo Fisher Scientific). Peptides were trapped on a Acclaim pepmap100, C18, 3μm, 75μm x 20mm nano trap-column (Thermo Fisher Scientific) and separated on a 75 μm x 250 mm, C18, 2μm, 100 Å Easy-Spray column (Thermo Fisher Scientific). The analytical separation was run for 90 min using a gradient of H_2_O/FA 99.9%/0.1% (solvent A) and CH_3_CN/FA 99.9%/0.1% (solvent B). The gradient was run as follows: 0-5 min 95 % A and 5 % B, then to 65 % A and 35 % B for 60 min, and 5 % A and 95 % B for 20 min at a flow rate of 250 nL/min. Full scan resolution was set to 60000 at m/z 200 with an AGC target of 3 x 106 and a maximum injection time of 60 ms. Mass range was set to 400-2000 m/z. For data dependent analysis, up to twenty precursor ions were isolated and fragmented by higher-energy collisional dissociation HCD at 27% NCE. Resolution for MS2 scans was set to 15000 at m/z 200 with an AGC target of 1 x 10^5^ and a maximum injection time of 60 ms. Isolation width was set at 1.6 m/z. Full MS scans were acquired in profile mode whereas MS2 scans were acquired in centroid mode. Dynamic exclusion was set to 20s.

Database search peak lists (MGF file format) were generated from raw data using the MS Convert conversion tool from ProteoWizard. The peaklist files were searched against the *Saccharomyces cerevisiae* reference proteome database (Uniprot,2025-03, 6066 entries), the Puf3-TAP bait protein sequence (*Saccharomyces* Genome Database (SGD), https://www.yeastgenome.org/) and an in-house database of common contaminant using Mascot (Matrix Science, London, UK; version 2.6.2). Trypsin was selected as the enzyme, with one potential missed cleavage. Precursor ion tolerance was set to 10 ppm and fragment ion tolerance to 0.02 Da. Variable amino acid modifications were oxidized methionine, deaminated (NQ) phosphorylated (STY) and GG (K). Fixed amino acid modification was carbamidomethyl cysteine. The Mascot search was validated using Scaffold 5.3.3 (Proteome Software). Peptide identifications were accepted if they could be established at greater than 33,0% probability to achieve an FDR less than 0,1% by the Percolator posterior error probability calculation (55). Protein identifications were accepted if they could be established at greater than 99,0% probability to achieve an FDR less than 1,0% and contained at least 2 identified peptides. Protein probabilities were assigned by the Protein Prophet algorithm (56). Proteins that contained similar peptides and could not be differentiated based on MS/MS analysis alone were grouped to satisfy the principles of parsimony.

For the PCA plots and differential expression analysis, normalised abundance values of mass spectrometry data from Puf3–TAP affinity purifications were log₂-transformed after addition of a pseudocount of one to stabilize variance and normalize signal distributions across samples. PCA analysis was then performed using the prcomp function in R and the first two principal components were plotted including all samples.

To carry out differential expression analysis, samples were categorized into three experimental groups: No-Tag controls (n = 3), Puf3–TAP in WT (n = 2), and Puf3–TAP in *not4Δ* (n = 3). A linear model was fitted using limma (57) (with a design matrix specifying these groups). Contrasts were defined to assess differential protein abundance between Puf3–TAP and NoTag, and between Puf3–TAP *not4Δ* and Puf3–TAP. Moderated t-statistics were computed using the empirical Bayes framework implemented in limma, providing robust variance estimation across proteins.

P-values were derived from these t-statistics and proteins were selected as upregulated if they showed a fold change > 1 and a p-value <0.05 in both comparisons, *not4Δ*/WT and *not4Δ*/ NoTag. Proteins were selected as downregulated if they showed a fold change > 1 and a p-value < 0.05 in both comparisons, WT/ *not4Δ* and WT/ NoTag.

The upregulated set and the downregulated set were then subjected to an enrichment test against gene ontology gene sets, with a handful of custom gene sets derived from our own data. This was done via a hypergeometric test, using all annotated proteins.

### ESI-LC-MSMS

Samples were dissolved in 19 μl of loading buffer (5% CH3CN, 0.1% FA). Biognosys iRT peptides were added to each sample and 2 μl of peptides were injected on the column. LC-ESI-MS/MS was performed on an Orbitrap Fusion Lumos Tribrid mass spectrometer (Thermo Fisher Scientific) equipped with an Easy nLC1200 liquid chromatography system (Thermo Fisher Scientific). Peptides were trapped on a Acclaim pepmap100, C18, 3μm, 75μm x 20mm nano trap-column (Thermo Fisher Scientific) and separated on a 75 μm x 500 mm, C18 ReproSil-Pur (Dr. Maisch GmBH), 1.9 μm, 100 Å, home-made column. The analytical separation was run for 135 min using a gradient of H_2_O/FA 99.9%/0.1% (solvent A) and CH_3_CN/H_2_O/FA 80.0%/19.9%/0.1% (solvent B). The gradient was run from 8 % B to 28 % B in 110 min, then to 42% B in 25 min, then to 95%B in 5 min with a final stay of 20 min at 95 % B. Flow rate was of 250 nL/min an total run time was of 160 min. Data-Independant Acquisition (DIA) was performed with MS1 full scan at a resolution of 60,000 (FWHM) followed by 30 DIA MS2 scan with sequential fix isolation windows of 28 m/z. MS1 was performed in the Orbitrap with an AGC target of 1 x 106, a maximum injection time of 50 ms and a scan range from 400 to 1240 m/z. DIA MS2 was performed in the Orbitrap using higher-energy collisional dissociation (HCD) at 30%, an AGC target of 1 x 10^6^ and a maximum injection time of 54 ms.

### Data analysis

DIA raw files were loaded into Spectronaut v.18 (Biognosys) and analyzed by directDIA using default settings. Briefly, data were searched against the *Saccharomyces cerevisiae* database (Uniprot release 2023-08, 6060 entries). Trypsin was selected as the enzyme, with one potential missed cleavage. Variable amino acid modifications were oxidized methionine. Fixed amino acid modification was carbamidomethyl cysteine. Both peptide precursor and protein FDR were controlled at 1% (*Q value* < 0.01). Single Hit Proteins were excluded. For quantitation, Protein LFQ method was set to “QUANT 2.0”, “only protein group specific” was selected as proteotypicity filter and normalization was set to “none”. The quantitative analysis was performed with Spectronaut, proteins were considered to have significantly changed in abundance with a Qvalue ≤ 0.05 and an absolute fold change FC ≥ |1.5| (log_2_FC ≥ |0.58|).

To assess the relative contributions of length and codon content on aggregation, a linear model was formulated. For each codon ’c’, its frequency in each transcript with a protein product present in aggregation data for *not4Δ* and *not4Δpuf3Δ* was calculated and then normalised to the transcript length in amino acids, giving the proportion of codon ’c’. Separately for each codon ’c’, a linear model was then constructed across transcripts with the relative aggregation rate log_2_ *not4Δ* / *not4Δpuf3Δ* of their protein products as the dependent variable and the proportion of codon ’c’ in each transcript and the natural logarithm of the length of each transcript as dependent variables. The contribution of the proportion of each codon ’c’ across transcripts to the change in aggregation rate between *not4Δ* and *not4Δpuf3Δ* in their protein products, while controlling for transcript length, was then calculated as the coefficient of the codon proportion variable in the linear model for codon ’c’. Assessing this value across all codons quantifies their relative contribution to change in aggregation between strains, while controlling for transcript length.

## RESULTS

### Puf3 contributes to the solubility of mRNAs in cells lacking Not4, in particular of mRNAs whose solubility is inversely regulated by Not1 and Not4

When analyzing our previously published data (17) for Puf3 targets, we found them enriched within the pool of mRNAs whose solubility was increased upon Not1 depletion but decreased upon Not4 depletion under fermentation conditions (**Supplementary Figure S1A**) (17). Hence, we compared the solubility of Puf3 target mRNAs in wild-type cells or in cells lacking Puf3. We performed RNA-Seq of the total and soluble mRNA pools and computed their log_2_ fold changes in cells lacking Puf3 versus WT (**Figure 1A**, x-values for total, y-values for soluble). We also evaluated solubility, which is defined as the log_2_ fold change of expression in the soluble RNA-Seq data set relative to expression in total RNA-Seq. To assess the effect of Puf3 loss on solubility, we subtracted WT solubility from solubility in cells without Puf3 (this value determines the color of the dots in **Figure 1A**). These values are plotted for Puf3 target mRNAs, classified according to the conservation of the consensus motif (32). As previously reported (32), Puf3 target mRNAs were more likely to be upregulated in the absence of Puf3 if the mRNAs carry a conserved consensus motif (compare category 1 to non-mitochondrial category 4 target mRNAs in **Figure 1A**). We observed no overall shift in mRNA solubility in cells lacking Puf3, regardless of the quality of the Puf3 consensus motif. Overall, the correlation of expression changes in soluble versus total slightly decreased if transcripts were more expressed in both soluble and total in the mutant relative to the wild type. This suggests some disruption in the solubility balance, but the divergence appears to have no specific direction. We noted that changes in expression of the Puf3 target mRNAs upon Not4 depletion followed similar trends as in cells lacking Puf3 (R= 0.39, **Supplementary Figure S1B,** left panel), consistent with the role of the Ccr4-Not complex for accelerated deadenylation of mRNA targets by Puf3 (32). However, we also noted that the expression of Puf3 target mRNAs increased in the soluble pool in *puf3Δ*, but much less so upon Not4 depletion, suggesting that different mechanisms are involved (**Supplementary Figure S1B**, middle panel). This, in turn, explains the lack of correlation of changes in mRNA solubility (**Supplementary Figure S1B**, right panel).

**Figure 1.**
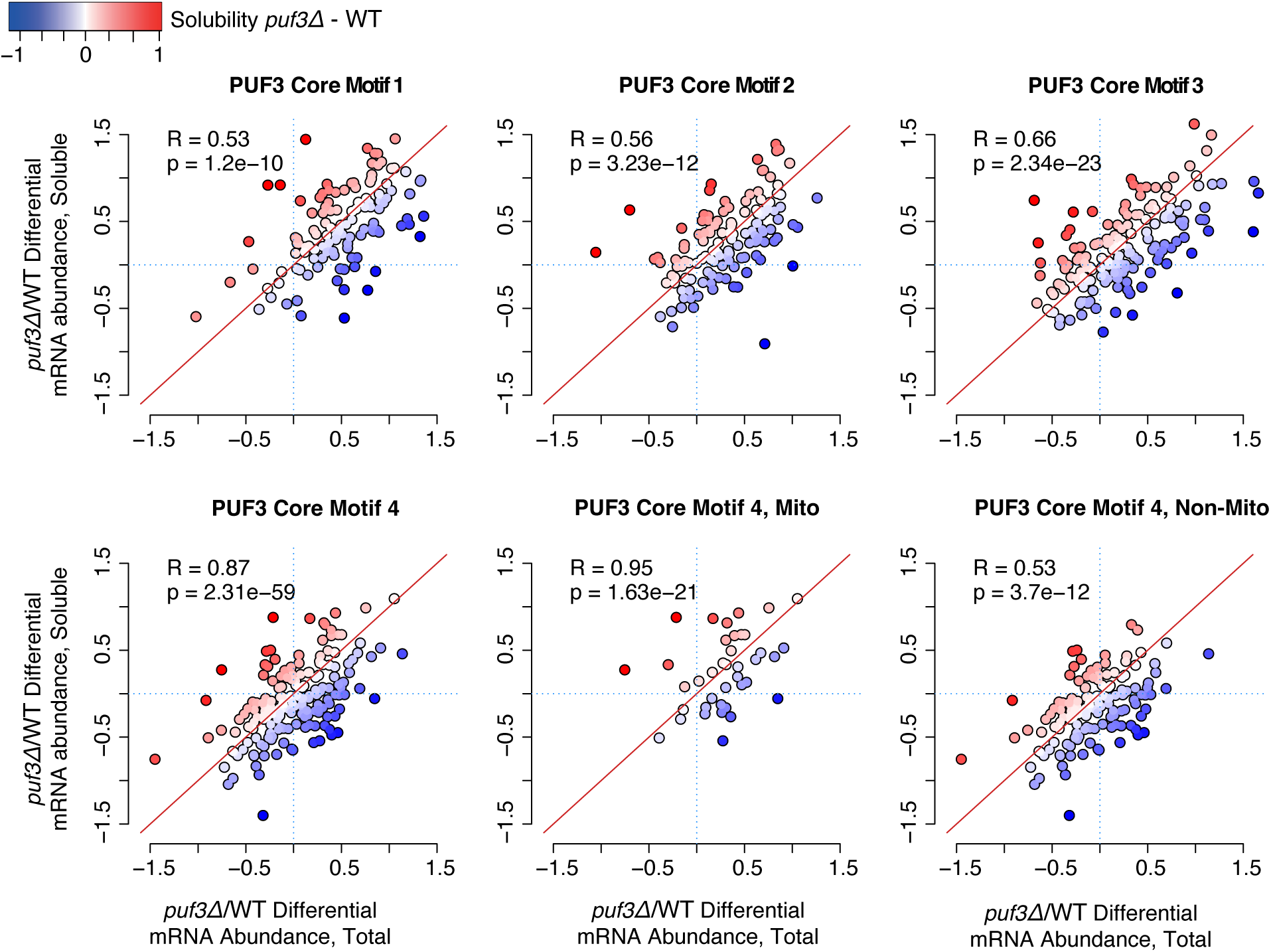
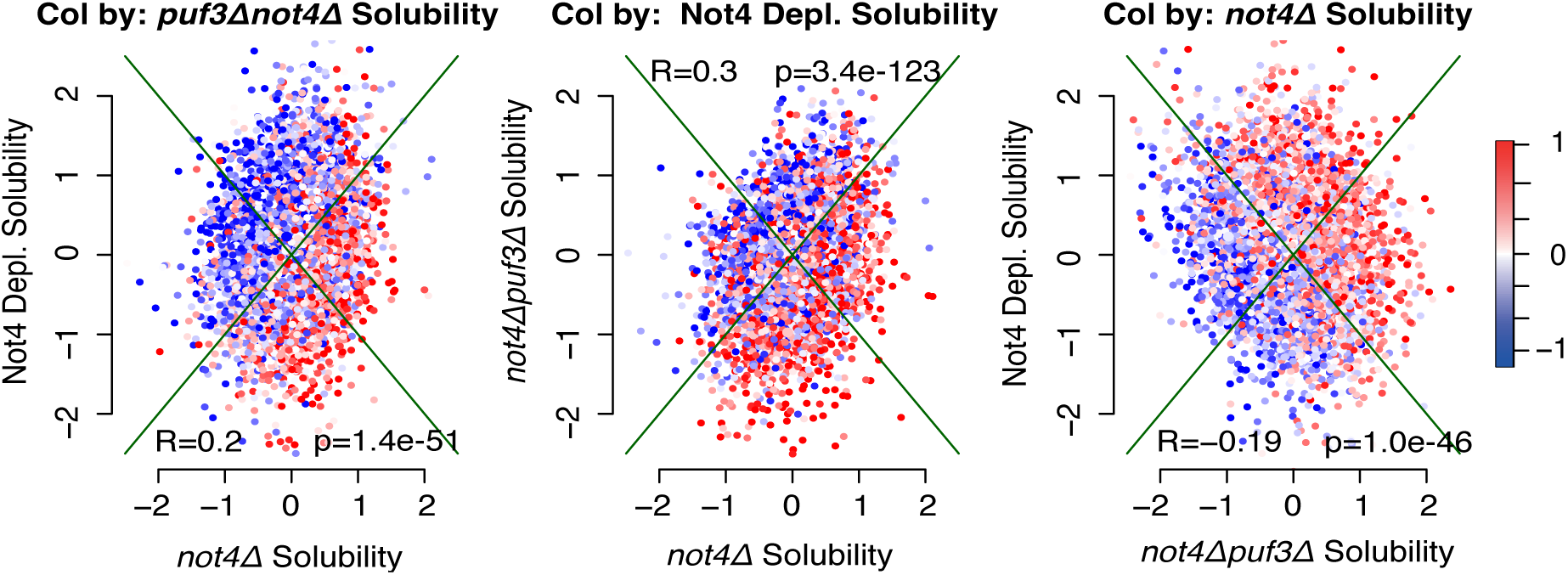
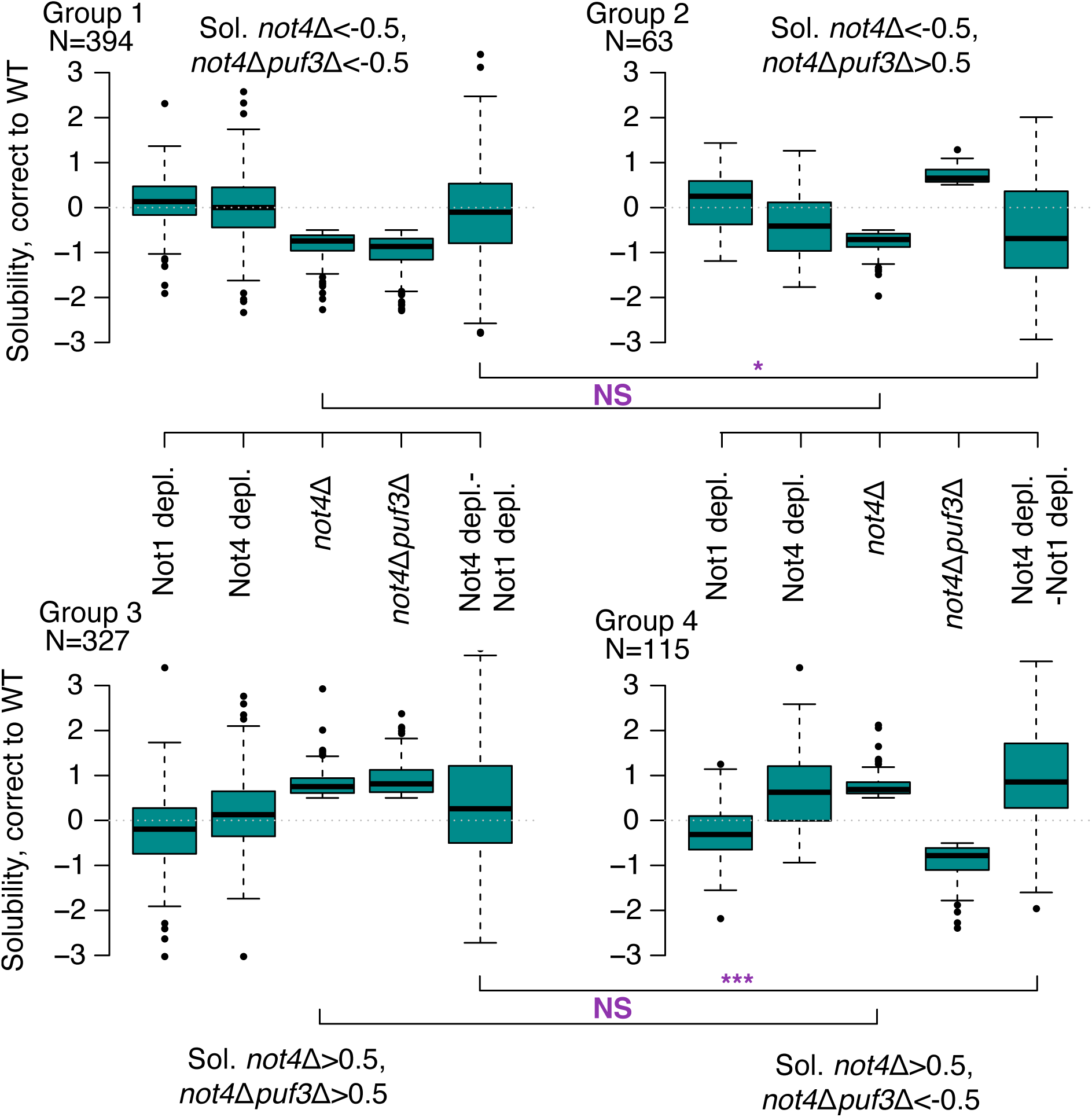
Puf3 counteracts changes in mRNA solubility in *not4Δ* specifically for mRNAs inversely regulated upon Not1 and Not4 depletion. **A.** Total RNA and RNA from cell lysates (soluble RNA) was extracted from wild type and *puf3Δ* and analyzed by RNA-Seq. Log_2_ fold changes in expression of total versus soluble RNA of Puf3 mRNA targets classified according to the quality of the Puf3 binding motif (32) in *puf3Δ* versus wild type cells are plotted against each other. Solubility (log_2_ ratio of RNA level in the lysate versus in the total RNA pool) differences in *puf3Δ* versus wild type cells defines the color of the dots. **B**. Pairwise scatterplots of log_2_ fold change in solubility change relative to WT of *not4Δ* versus Not4 depletion (left panel, dots colored by *not4Δpuf3Δ* solubility), *not4Δ* versus *not4Δpuf3Δ* (middle panel, dots colored by Not4 depletion solubility), *not4Δpuf3Δ* versus Not4 depletion (right panel, dots colored by *not4Δ* solubility) **C**. mRNAs that show a log_2_ fold change in solubility in *not4Δ* or *not4Δpuf3Δ* compared to wild type smaller than -0.5 are represented in the upper left panel; those that show a log_2_ fold change in solubility in *not4Δ* compared to wild type smaller than -0.5 but in *not4Δpuf3Δ* greater than 0.5 in the upper right panel; those that show a log_2_ fold change in solubility in *not4Δ* compared to wild type or *not4Δpuf3Δ* compared to wild type a log_2_ fold change in solubility greater than 0.5 in the lower left panel, and finally those that show a log_2_ fold change in solubility in *not4Δ* compared to wild type higher than 0.5, but in the *not4Δpuf3Δ* lower than -0.5 in the lower right panel. In each panel the change in solubility of the relevant pool of mRNAs of the panel upon Not1 and Not4 depletion is indicated (left 2 columns of each panel). Finally, the difference in change of solubility of each pool of mRNAs upon Not1 and Not4 depletion is shown in the right column of each panel.

Since Puf3-target mRNAs are enriched within the pool of mRNAs less soluble upon immediate Not4 depletion, we looked at the impact of Puf3 on solubility of mRNAs in cells lacking Not4 altogether (see strain list in **Supplementary Table S1**). From previous studies (17) we know that the complete absence of the Not4 E3 ligase has phenotypes distinguishable from those measured immediately upon Not4 depletion. With regard to mRNA solubility, we noted that *not4Δ* weakly correlates with Not4 depletion, it also weakly correlates with *not4Δpuf3Δ,* while the depletion of Not4 weakly anti-correlates with *not4Δpuf3Δ* (**Figure 1B**). The fact that *not4Δpuf3Δ* and Not4 depletion solubilities anticorrelate, despite both having a weak positive correlation with *not4Δ* solubility, suggests that they both often diverge from *not4Δ* and frequently in opposite directions. Indeed, we see that solubility changes in Not4 depletion are decreased in transcripts increasing solubility in *not4Δpuf3Δ* relative to *not4Δ* (**Figure 1B**, middle panel). To investigate this further we created boxplots of solubilities comparing transcripts where *not4Δ* and *not4Δpuf3Δ* have opposite effects, to those where their effects are similar. We observed that, among mRNAs which lose solubility in *not4Δ* (**Figure 1C**, upper panels), some reverse this change in *not4Δpuf3Δ* (**Figure 1C**, upper right panel) and some do not (**Figure 1C**, upper left panel). Symmetrically, among mRNAs which gain solubility in *not4Δ* (**Figure 1C**, lower panels), some reverse this change in *not4Δpuf3Δ* (**Figure 1C**, lower right panel) and some don’t (**Figure 1C**, lower left panel). Those transcripts for which solubility change in *not4Δ* is reversed in *not4Δpuf3Δ* show similar changes in solubility in *not4Δ* and upon Not4 depletion (**Figure 1C**, right panels). This was not the case for the transcripts for which solubility changes in *not4Δ* is not reversed in *not4Δpuf3Δ* (**Figure 1C**, left panels). Moreover, the differences in solubilities between Not4 and Not1 depletions is significantly higher for mRNAs whose changes in solubility in *not4Δ* is reversed in *not4Δpuf3Δ* compared to those that are not (**Figure 1C**, compare last columns of right and left panels).

Taken together, these results indicate that Puf3 does not specifically contribute to the solubility of its own target mRNAs, but it can reverse changes of solubility due to the absence of Not4, in particular for mRNAs inversely regulated by Not1 and Not4, and whose solubility changes similarly upon Not4 depletion and in the complete absence of Not4.

### Puf3 contributes to protein aggregation and temperature sensitivity in cells lacking Not4

Previous studies have indicated that Not4 is an important player in protein homeostasis and that new proteins aggregate in the absence of Not4 (21,22,58). We proposed that condensation of mRNAs by the Not proteins, hence potentially regulation of mRNA solubility, could be an active mechanism during translation elongation (19). We thus questioned whether the deletion of Puf3 that alters mRNA solubility in *not4Δ* (see above **Figure 1C**), also had an impact on protein aggregation in this mutant. We prepared cell lysates from which we isolated aggregated proteins, for wild type cells, single mutants and the double *not4Δpuf3Δ* mutant. We analyzed the profile of proteins in the total lysate and in the aggregates by SDS-PAGE and silver staining. The profile of total proteins in cell lysates was similar in all 4 strains (**Figure 2A**, lanes 1-4). However, comparing the same amount of total protein lysate, there were few proteins in aggregates from wild type cells (**Figure 2A**, lane 5) compared to much higher protein levels in aggregates from cells lacking Not4 (**Figure 2A**, lane 6). In cells lacking Puf3, the level of protein aggregation was comparable to that in wild type cells (**Figure 2A**, compare lane 7 to lane 5). The deletion of Puf3 in *not4Δ* reduced protein aggregation, however to levels that remained higher than in wild type cells (**Figure 2A**, compare lane 8 to lanes 6 and 5).

**Figure 2.**
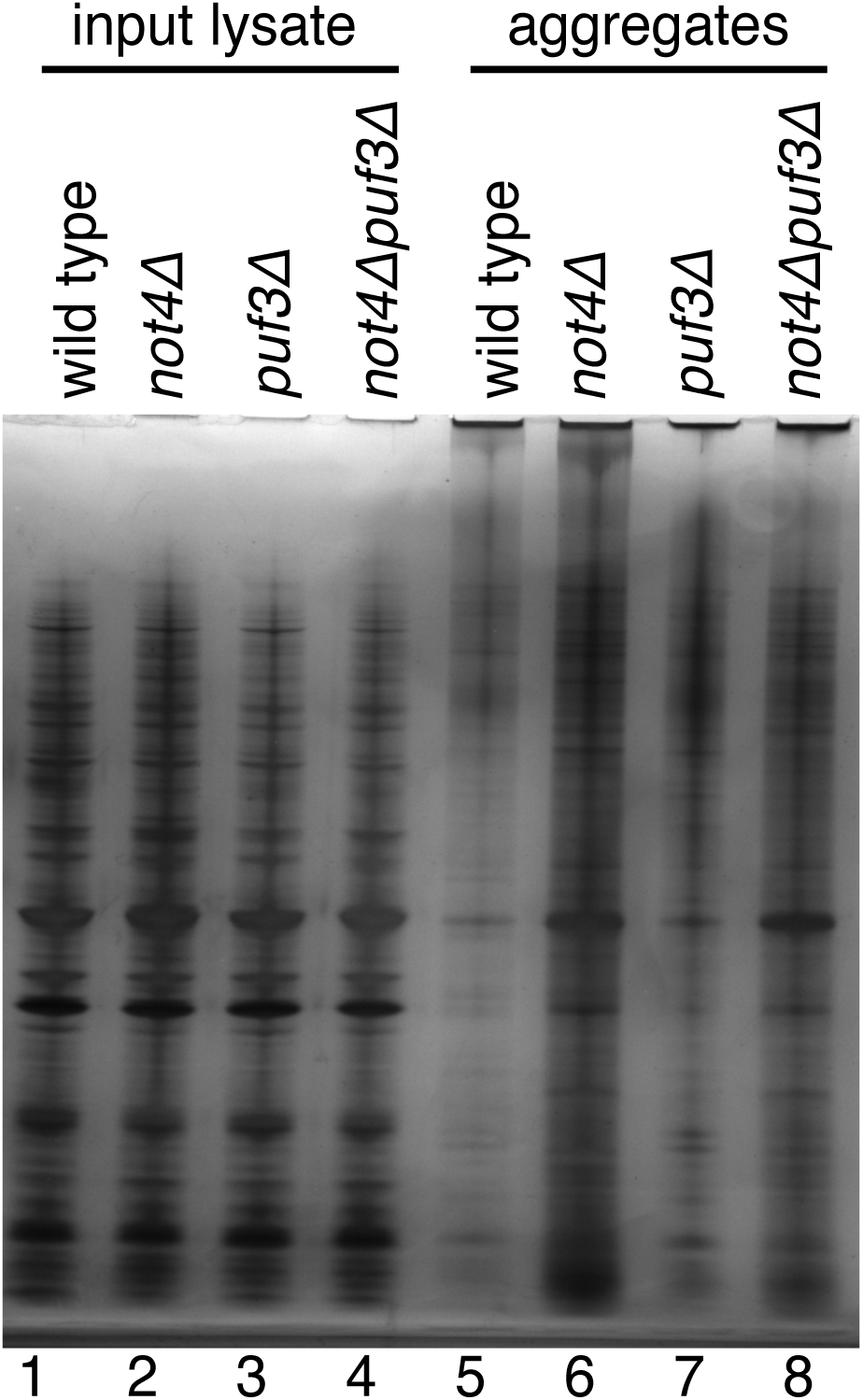
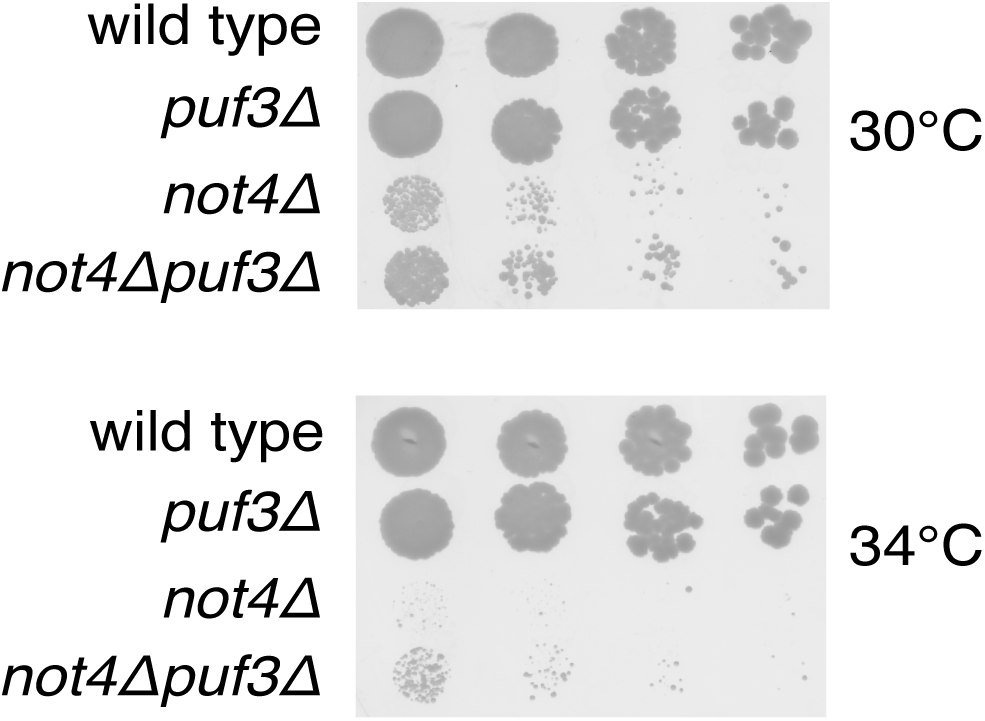
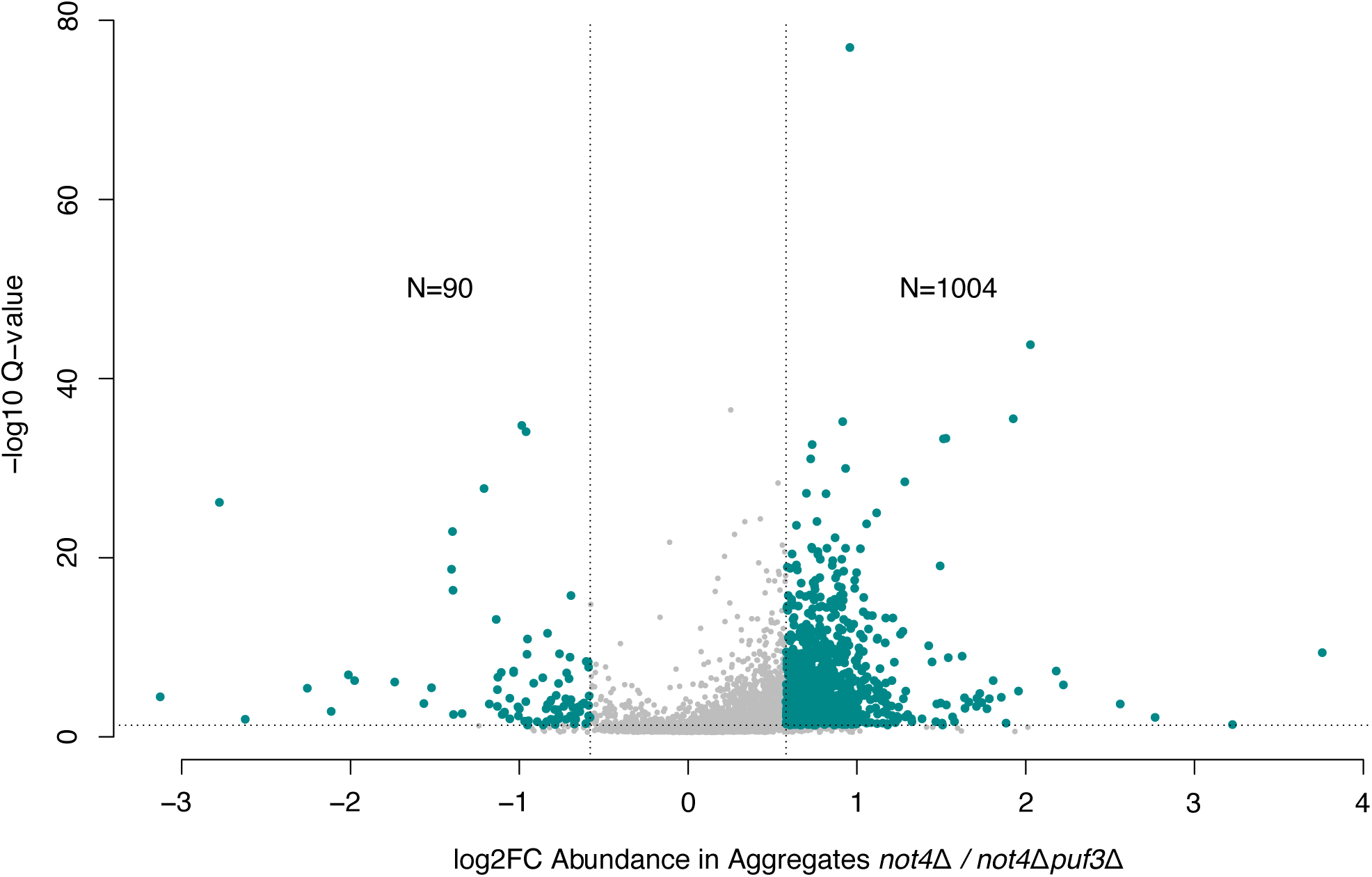
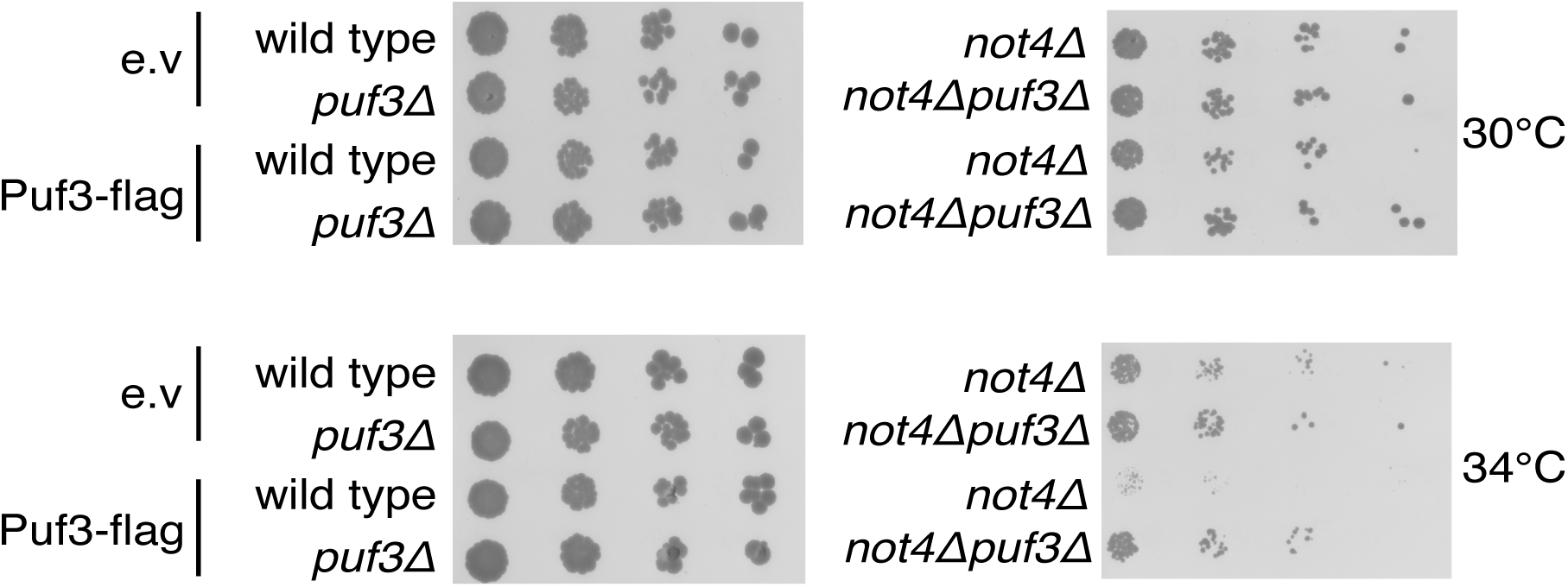
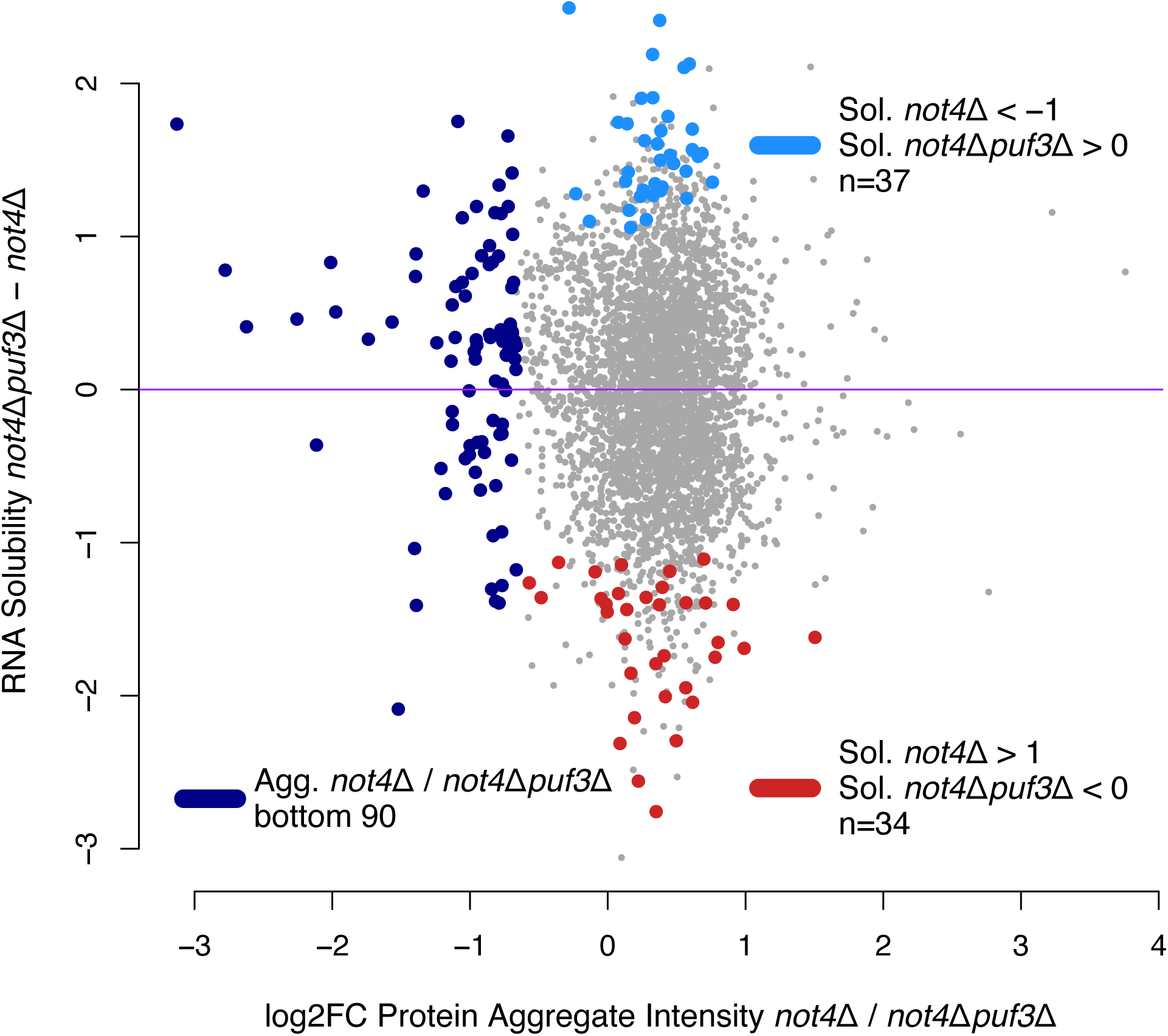
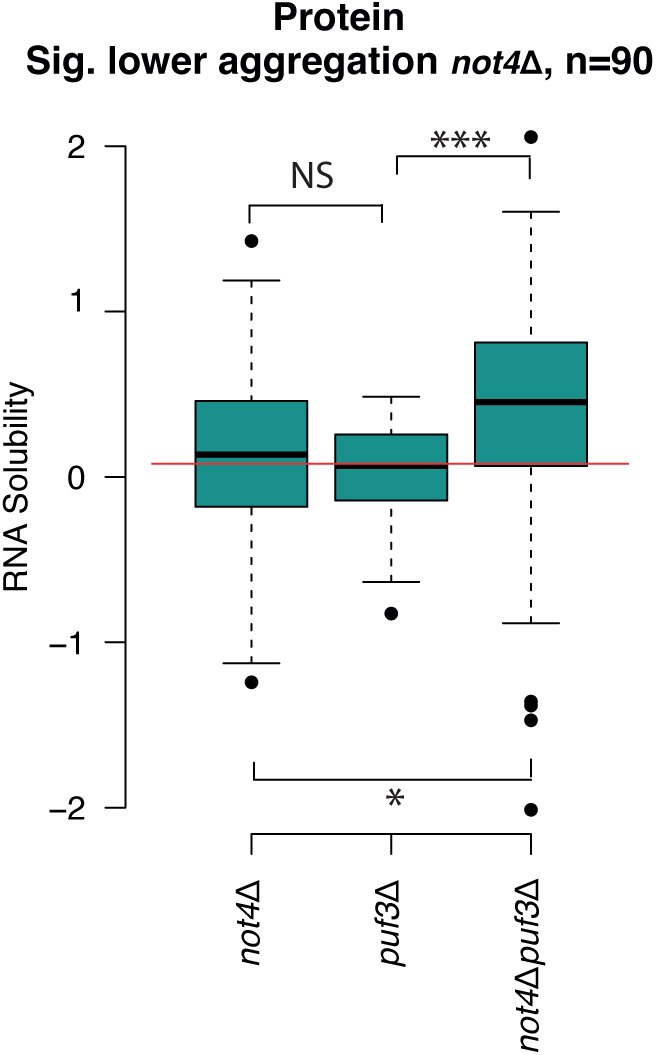
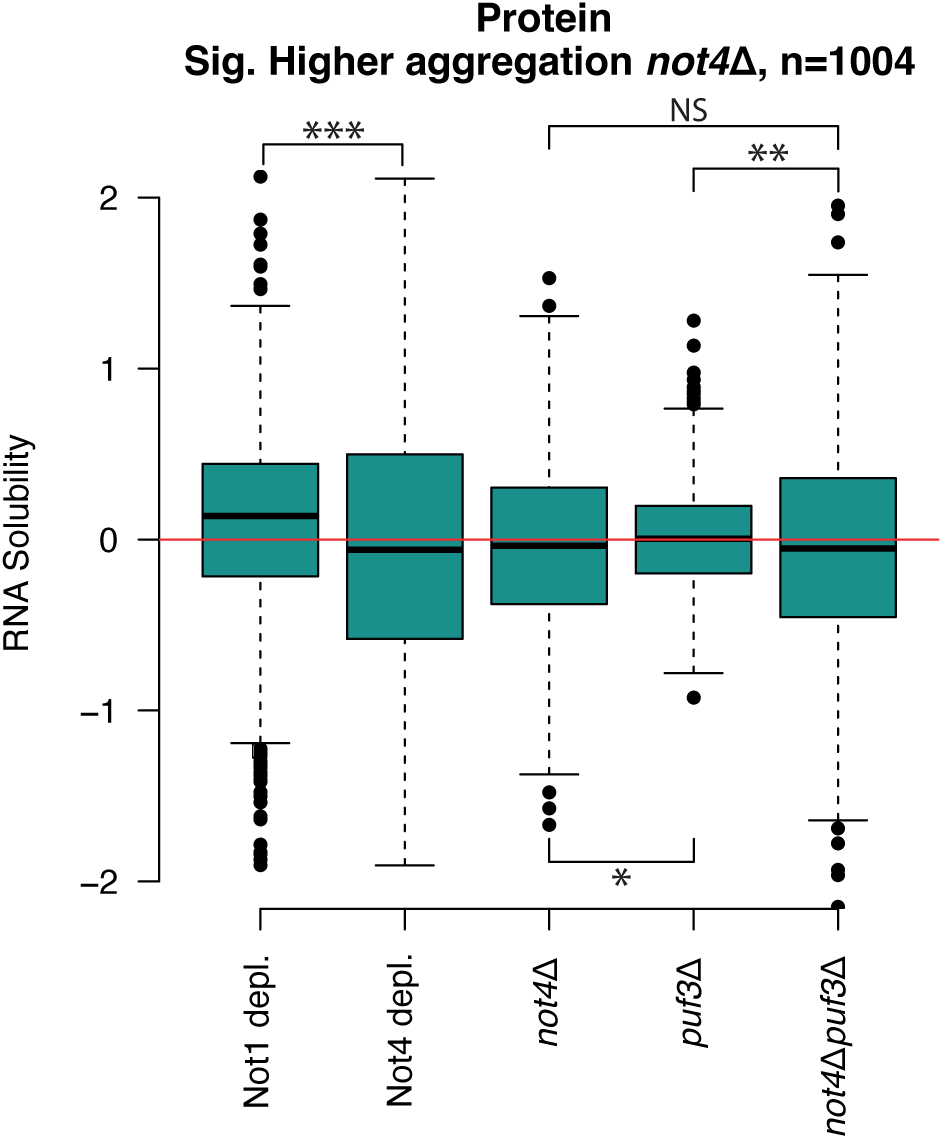
Puf3 contributes to protein aggregation and temperature sensitivity of *not4Δ* cells. **A**. Aggregates (aggregates, lanes 5-8) were isolated from total cellular lysates (input lysate, lanes 1-4) prepared from wild type, *not4Δ*, *puf3Δ* and *not4Δpuf3Δ* cells and analyzed by SDS-PAGE and silver staining. **B**. Spot assay comparing the growth of wild type, *puf3Δ*, *not4Δ*, and *not4Δpuf3Δ* cells. Cells were serially diluted and spotted on YPD plates that were placed at 30°C or 34°C and left to grow for 3 days. **C.** Aggregated proteins from the same amount of protein lysates from *not4Δ* and *not4Δpuf3Δ* in triplicate, were collected for quantitative mass spectrometry. The quantitative differential presence of identified proteins in the 2 strains is represented by a volcano plot (log2FC is log_2_ fold change). **D**. Spot assay comparing the growth of wild type, *not4Δ*, *puf3Δ* and *not4Δpuf3Δ* cells transformed with a plasmid expressing Puf3-flag or an empty vector (*e.v*, pRS416) as a control. Cells were serially diluted and spotted on -Ura plates that were placed at 30°C or 34°C and left to grow for 3 days. **E**. Scatterplot comparing log_2_ fold changes (log2FC) in solubility of mRNAs in *not4Δpuf3Δ* relative to *not4Δ* on the y axis and log_2_ fold differences in protein presence in aggregates from *not4Δ* relative to *not4Δpuf3Δ* on the x axis. mRNAs encoding proteins showing the 90 most increased protein aggregation in the double mutant are indicated in dark blue, those with greatest increase in solubility in the double mutant compared to *not4Δ* in light blue (log_2_ fold change in *not4Δ* below -1 and in *not4Δpuf3Δ* above 0) and those with the greatest decrease in solubility in the double mutant compared to *not4Δ* in red. **F**. Box plot comparing changes in solubility in single mutants and in the double mutant compared to the wild type for the 90 mRNAs whose encoded proteins aggregate significantly more in the double *not4Δpuf3Δ* mutant compared to *not4Δ*. **G**. Box plot as in **F** but for the 1004 mRNAs whose encoded proteins aggregate significantly less in the double *not4Δpuf3Δ* mutant compared to *not4Δ* and also including changes in solubility upon Not1 and Not4 depletion.

To determine whether the differences in protein aggregation levels was functionally relevant, we compared the growth of the different strains at a higher temperature when the pressure for protein folding is increased. Mirroring the differences in protein aggregation, we noted that *not4Δ* was temperature sensitive, corroborating earlier observations (21), while this sensitivity was partially suppressed at 34°C by the additional deletion of Puf3 (**Figure 2B**).

We next identified and compared the proteins in aggregates in *not4Δ* and in *not4Δpuf3Δ* by quantitative mass spectrometry (**Supplementary Figure S2A** and **Supplementary Table S2**). Only few proteins aggregated more in the double mutant compared to the single mutant (**Figure 2C**, 90 proteins with a Q-value < 0.05 and a log_2_ fold change < -0.5 on the left of the volcano plot), whereas instead a great number of proteins aggregated less in the double mutant compared to the single mutant (**Figure 2C**, 1004 proteins with a Q-value < 0.05 and a log_2_ fold change > 0.5 on the right of the volcano plot). Hence, these results confirm that Puf3 contributes to protein aggregation in cells lacking Not4.

Since the deletion of Puf3 has a suppression effect in *not4Δ* we tested the consequence of overexpressing Puf3 in *not4Δ* at 30°C and 34°C (**Figure 2D**). Overexpression of Puf3 on a centromeric plasmid with its own promoter and terminator had no detectable effect on the cell growth of wild type cells or of cells lacking Puf3, but it was toxic in *not4Δ* at 34°C. The toxicity of Puf3 overexpression was also visible in *not4Δpuf3Δ* cells, though to a lesser degree. Thus, Puf3 has a dominant negative effect in *not4Δ*.

### Changes in protein aggregation and mRNA solubility do not generally correlate when Puf3 is deleted in *not4Δ*

We next addressed whether differences in mRNA solubility between the *not4Δ* single mutant and the *not4Δpuf3Δ* double mutant correlated with changes in protein aggregation. We noted that proteins that aggregated less in the double mutant compared to the single mutant were expressed from mRNAs that were either more or less soluble (**Figure 2E**, dark blue mRNAs) and that proteins encoded by mRNAs with the most extreme changes in solubility did not show any particular trend with regard to changes in protein aggregation (**Figure 2E**, light blue and red mRNAs).

We next focused on the 90 proteins that aggregated more in the double mutant compared to the single mutant (**Supplementary Table S2**). Overall their mRNAs were more soluble in the double mutant cells (**Figure 2F**) and they were highly enriched for mRNAs that encode interactors of the Hsp90 chaperone (59) and targets of Puf3 (32) (**Supplementary Figure S2B**, **Supplementary Table S3**). This observation suggests that Not4 and Puf3 collectively reduce mRNA solubility of specific mRNAs to safeguard that the encoded proteins remain soluble.

We then looked at the mRNA set encoding proteins that aggregated less in the double mutant cells (**Supplementary Table S2**). They did not show a particular common trend regarding changes in mRNA solubility. Interestingly, however, these were mRNAs showing opposite changes in solubility upon Not1 and Not4 depletion (**Figure 2G**) and they were most highly enriched for GO-terms related to ribosomal biogenesis and cytoskeleton organization (**Supplementary Figure 2C**, **Supplementary Table S4**).

### Puf3 deletion reduces accumulation of ribosomes at the stop codon in *not4Δ*

Since the mRNAs encoding proteins that aggregated less in double mutants compared to *not4Δ* did not show a particular trend for changes in mRNA solubility, we focused on translation elongation dynamics. We performed Ribo-Seq analysis comparing wild type cells upon Not1 or Not4 depletion, in presence and in absence of Puf3.

Metagene analyses for the different strains revealed that ribosomes accumulated at the stop codon specifically upon depletion of Not4, as observed previously when sequencing decay intermediates (17). Interestingly this increase was partially suppressed upon deletion of Puf3 (**Figure 3A**). By assessing the differences between stop codon coverage across all transcripts that had sufficient depth and showed a significant increase upon depletion of Not4 relative to WT and upon depletion of Not4 in cells lacking Puf3, we could determine that the accumulation of ribosomes upon Not4 depletion at the stop codon was significantly less when Puf3 was absent (**Supplementary Figure S3A)**. At the start codon, ribosomes accumulated less in all of the mutant strains compared to the wild type (**Figure 3B**).

**Figure 3.**
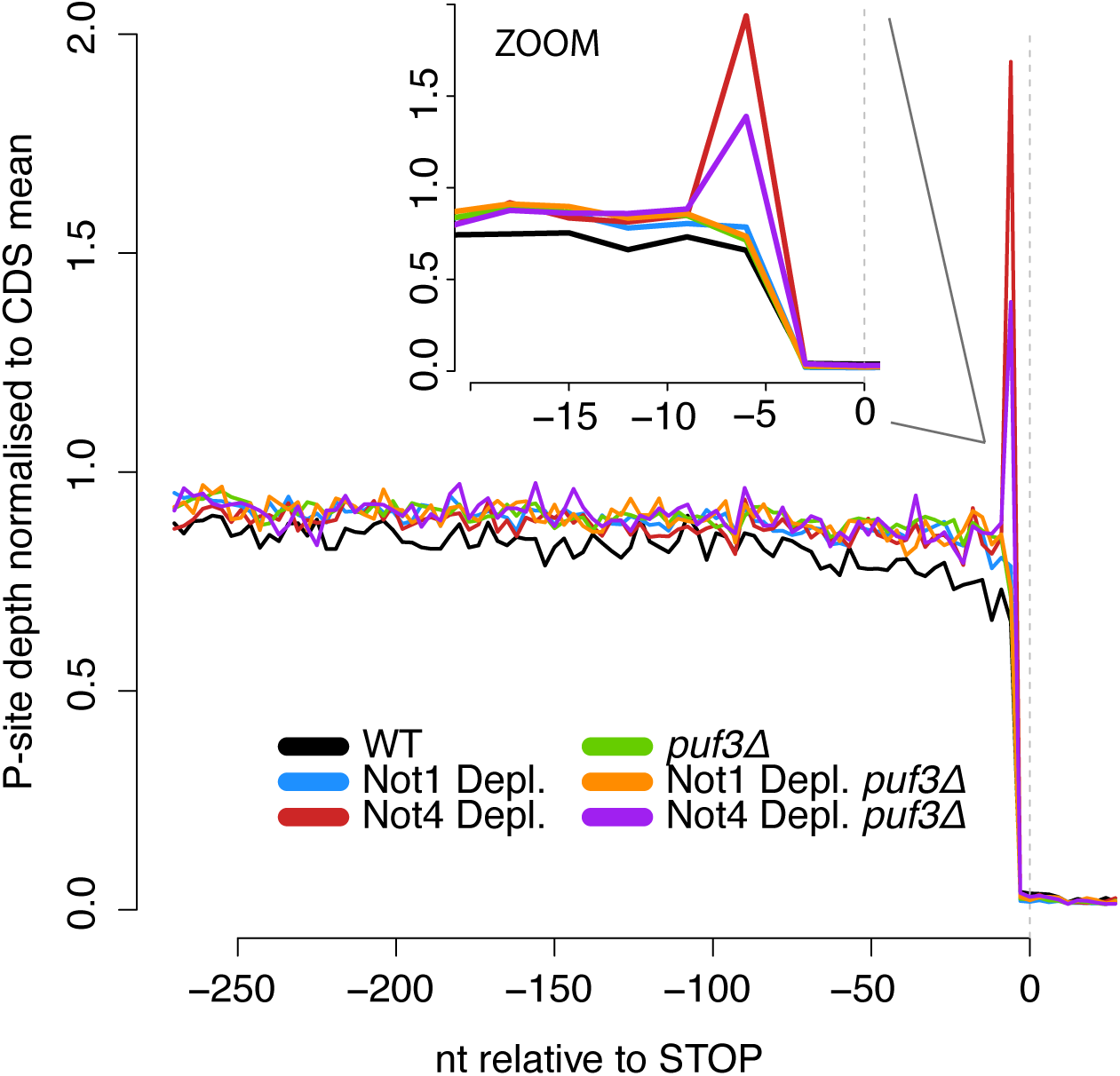
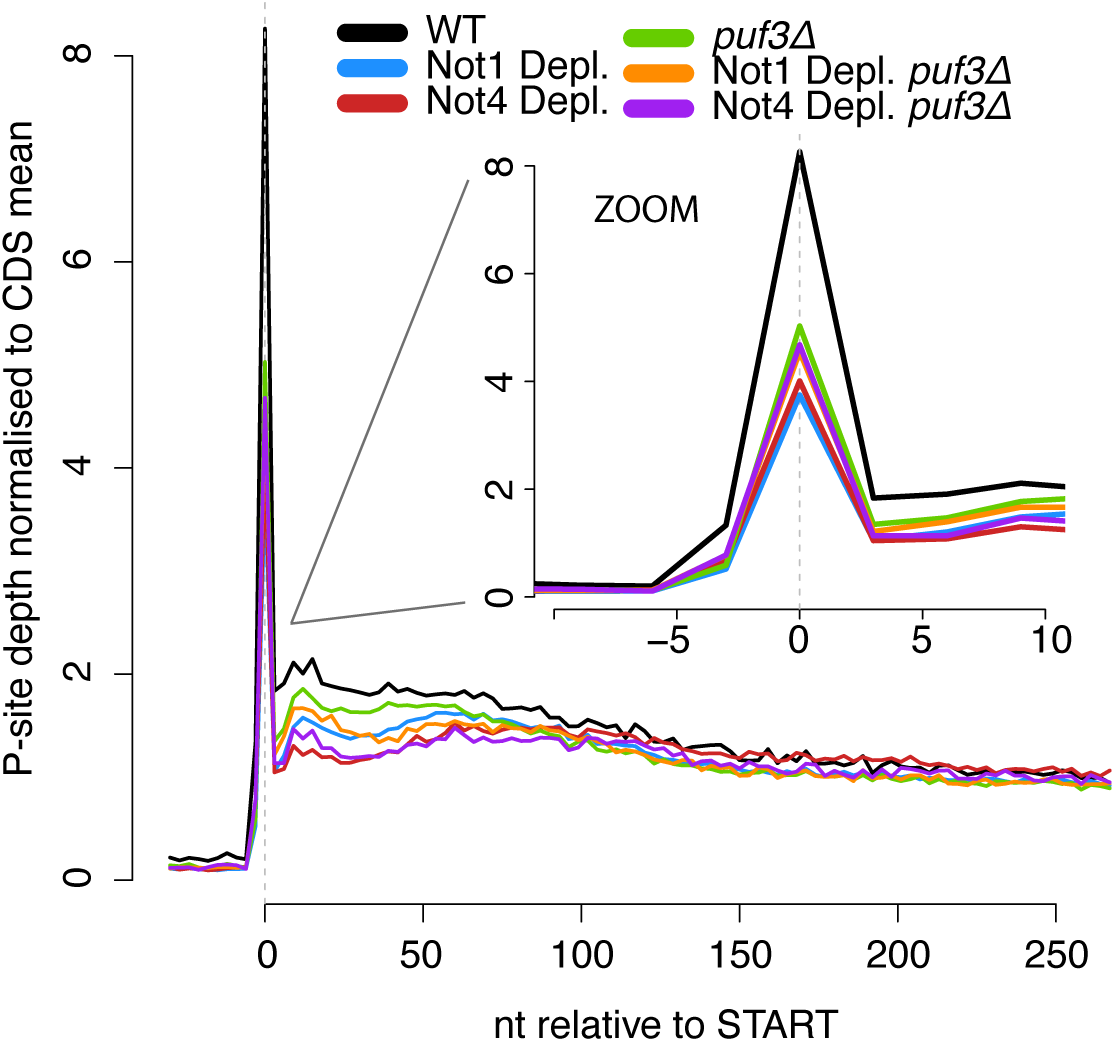
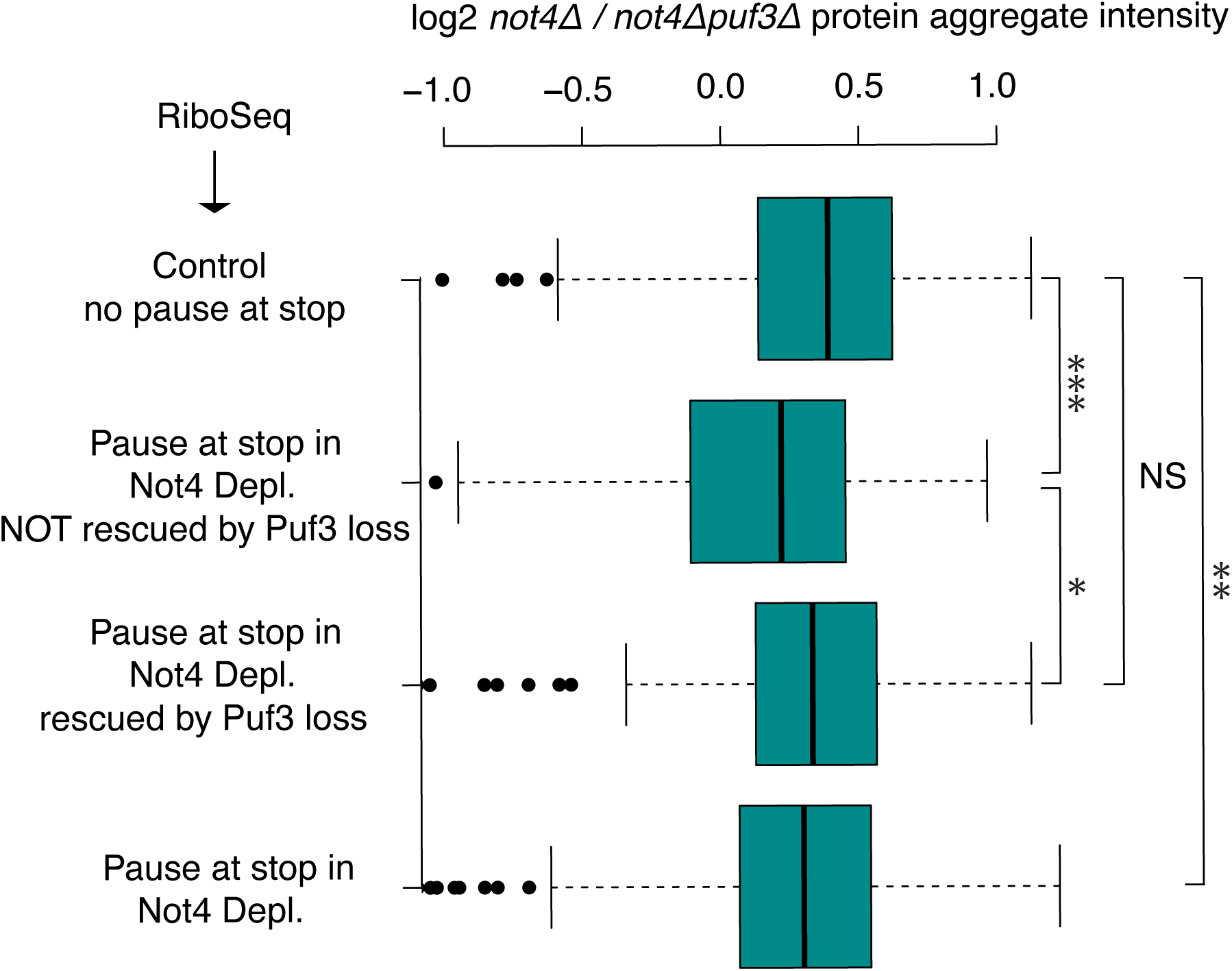
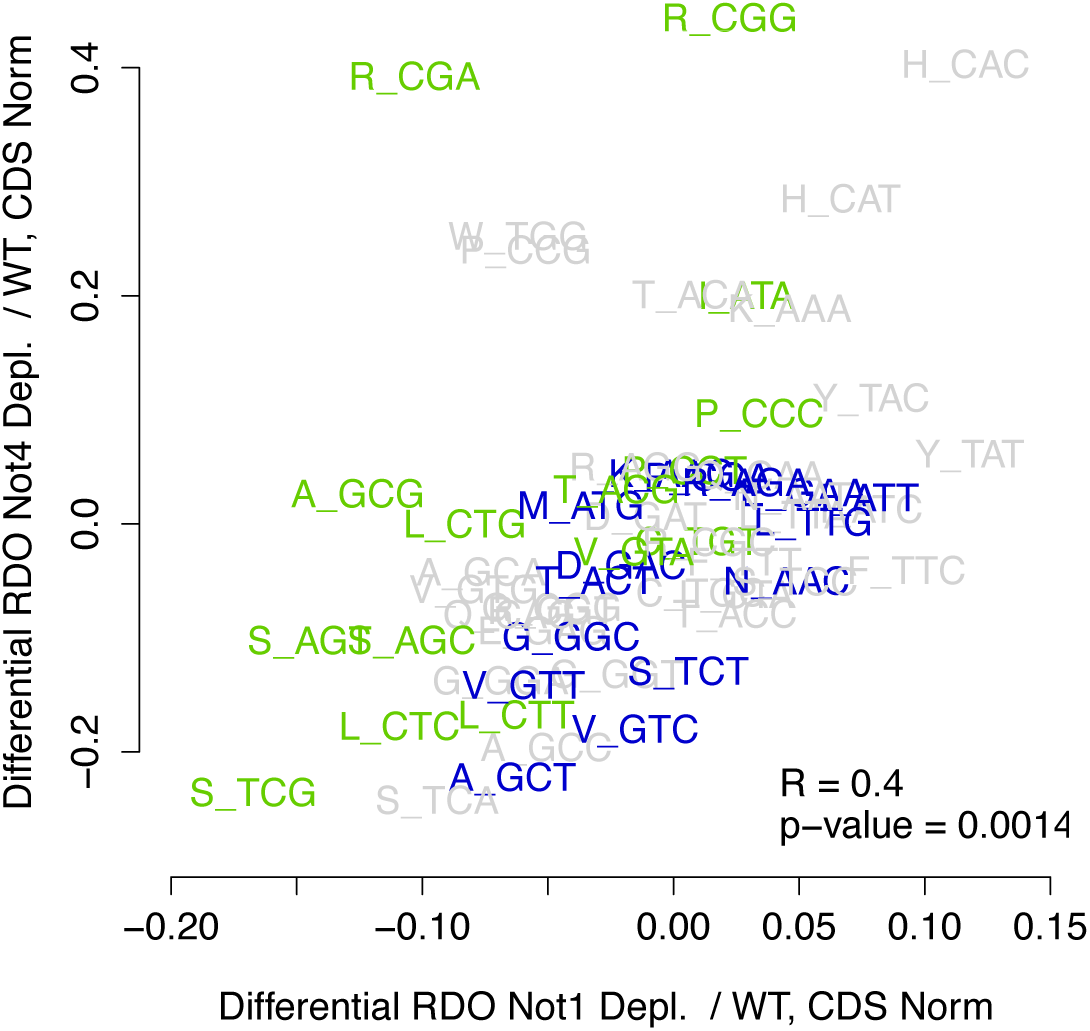
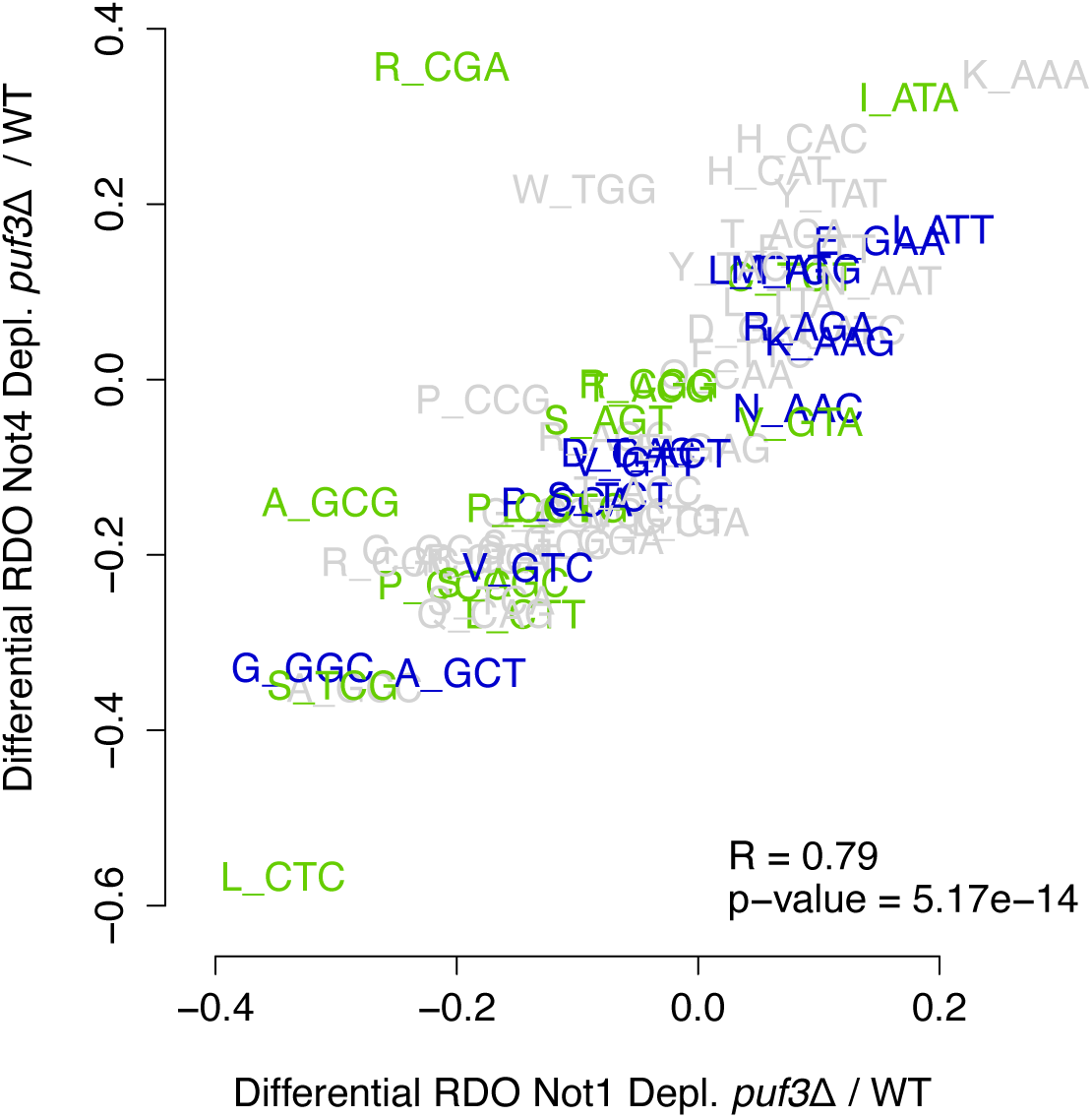
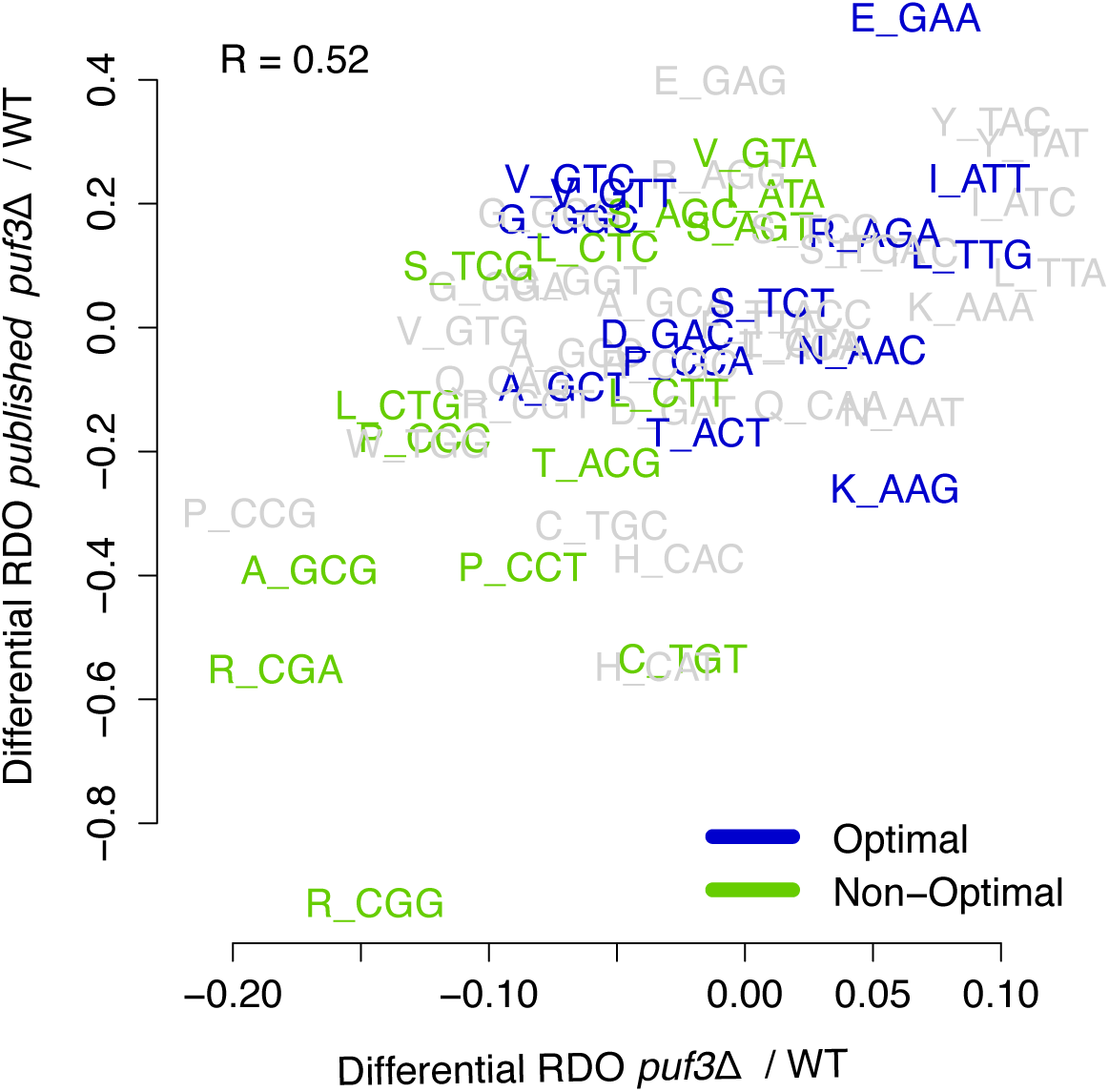
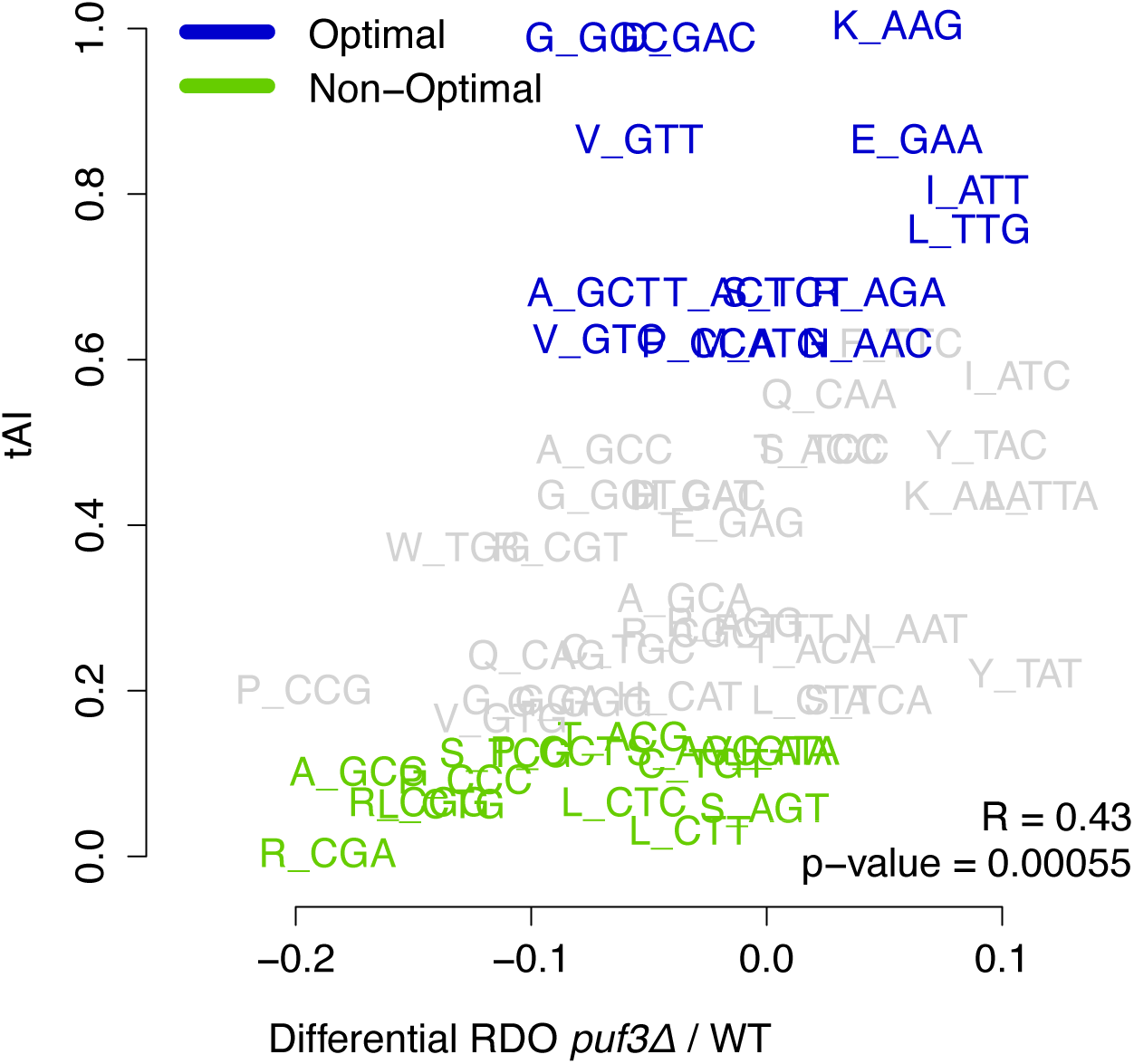
Difference in changes of codon-specific A-site ribosome dwelling occupancies upon Not1 and Not4 depletion are in large part due to Puf3. Metagene analysis comparing ribosome dwelling at the stop (**A**) and start (**B**) codons, in wild type and upon Not1 or Not4 depletion with or without Puf3, extracted from Ribo-Seq. **C.** Box plot of log_2_ fold changes in protein aggregation in *not4Δ* versus *not4Δpuf3Δ* for mRNAs that show increased ribosome pausing at stop or not upon Not4 depletion and for the former that is suppressed or not in the double mutant. (**D-G**). Scatterplot analyses comparing codon-specific changes in A-site ribosome dwelling occupancies, upon Not1 and Not4 depletion **(D**); upon Not1 and Not4 depletion in cells lacking Puf3 **(E)**; for *puf3Δ* cells growing in glucose (x-axis) compared to *puf3Δ* cells transferred to respiratory growth (45) (y-axis) (**F**) and finally for *puf3Δ* cells growing in glucose (x-axis) compared to the tRNA adaptation index (51) (y-axis) (**G**).

We compared the mRNAs according to the change in accumulation of ribosomes at the stop codon upon Not4 depletion in presence and absence of Puf3, regarding differences in protein aggregation of encoded proteins and mRNA solubility in *not4Δ* versus *not4Δpuf3Δ* (**Supplementary Figure S3B**). There was no correlation between changes in mRNA solubility and changes in ribosome occupancy at the stop codon overall. However, we noted that mRNAs without ribosome accumulation at the stop codon upon Not4 depletion compared to wild type exhibited higher aggregation of encoded proteins in *not4Δ* relative to the double mutant than those with higher ribosome occupancy at the stop (**Figure 3C**, compare “Control no pause at stop” to “Pause at stop upon Not4 depletion”). Moreover, mRNAs for which the occupancy at stop was not reduced in the double mutant showed less aggregation of encoded proteins in the single versus double mutant (**Figure 3C**, “Pause at stop upon Not4 depletion NOT rescued by Puf3 loss”). Instead, those with occupancy reduction showed the same aggregation of encoded proteins in the single versus double mutant as mRNAs without ribosome accumulation (**Figure 3C**, compare “Control no pause at stop” to “Pause at stop upon Not4 depletion rescued by Puf3 loss”). Consistently, GO-term analysis of the mRNAs with increase in ribosome occupancy at the stop codon upon depletion of Not4 were enriched in mRNAs encoding proteins prone to aggregate in *not4Δ*. Additionally, they were enriched for mRNAs whose solubility was inversely regulated upon Not1 and Not4 depletion, mRNAs encoding mitochondrial ribosomal proteins and Hsp90 interactors, interactors of the Zuo1 co-chaperone and Puf3 core targets, as well as mRNAs more soluble in *not4Δpuf3Δ* than in *not4Δ*, and mRNAs with ribosome accumulation at the start in the absence of Not4 or Not5 (**Supplementary Table S5,** “Not4 and Not5_PeakStart” category) (19). These observations link aggregation of proteins in *not4Δ* to increased ribosome occupancy at start or stop codons, mRNA solubility and Puf3 function (**Supplementary Table S5**).

### Puf3 affects translation elongation dynamics related to codon optimality

We next evaluated codon-specific changes in A-site ribosome dwelling occupancies (RDOs) in the different strains. In presence of Puf3, changes in A-site RDOs showed only minimal correlation upon Not1 and Not4 depletion, with notably significant differences in dwelling at the two yeast rare arginine codons, CGA and CGG (**Figure 3D**, R=0.4). Indeed, we detected higher RDOs at CGG or CGA codons in the A site upon Not4 depletion, which remained either unaffected or decreased upon Not1 depletion. In Puf3 absence, the overall correlation between changes in codon-specific A-site RDOs upon Not1 and Not4 depletion markedly increased (**Figure 3E**, R=0.79). These findings indicate that Puf3 affects differences in A-site RDOs upon Not1 and Not4 depletion. Importantly Puf3 contributed significantly to differences in changes of A-site RDOs upon Not1 and Not4 depletion for mRNAs whether they were or not targets of Puf3 (**Supplementary Figure S3C**). Of note, the A-site RDOs at rare arginine codons upon Not4 depletion was more markedly increased for mRNAs that are not targets of Puf3 (**Supplementary Figure S3C**, upper panels) than for Puf3 targets (**Supplementary Figure S3C**, lower panels), whereas in Puf3 targets the A-site RDOs decreased more markedly at rare arginine codons upon Not1 depletion (**Supplementary Figure S3C**, lower panels) compared to non-Puf3 targets (**Supplementary Figure S3C**, upper panels).

These results indicate that in the context of a mutant Not function, Puf3 impacts translation elongation dynamics in yeast cells growing fermentatively, conditions where it has been thought so far to inhibit its target mRNAs, either via degradation or translation inhibition. To determine whether this was also true in the context of unaffected Not protein function, we analyzed codon-specific A-site RDO changes between wild type and *puf3Δ* cells growing in glucose. The deletion of Puf3 had a codon-specific effect on A-site RDO changes in cells growing fermentatively. They correlated (R=0.52) with codon-specific A-site RDO changes between wild type and *puf3Δ* cells grown under respiratory conditions, as calculated from published ribosome profiling data (45) (**Figure 3F**). Interestingly, reduced A-site RDOs at rare arginine codons was the most significant change observed in cells lacking Puf3. In addition, there was a weak but noticeable correlation (R=0.43) between changes in A-site RDOs and codon optimality (**Figure 3G**) and to changes observed upon depletion of a translation initiation and elongation factor, eIF5A, as calculated from published ribosome profiling data (60) (**Supplementary Figure S3D**, R= 0.30). This is interesting considering our previous observation that changes in A-site RDOs upon eIF5A depletion inversely correlate to changes observed in absence of Not5 (19).

### Dwelling at rare arginine codons decreased in *not4Δpuf3Δ* relative to *not4Δ* but only for mRNAs encoding proteins that aggregate less in *not4Δpuf3Δ* relative to *not4Δ*

The findings above show that changes in A-site RDOs at rare arginine codons is the most noticeable change upon Not4 depletion (increased) and in the absence of Puf3 (decreased). To determine how codon-specific changes in A-site RDOs are related to changes in protein aggregation between *not4Δ* and *not4Δpuf3Δ*, we compared two equal sets of 300 mRNAs. The first group are the 300 mRNAs encoding proteins that aggregate in *not4Δ* but show the least change in aggregation of encoded proteins in the double mutant. The second group are the 300 mRNAs encoding proteins that aggregate in *not4Δ* but exhibit the greatest reduction in aggregation of encoded proteins in the double mutant. For these two categories, we computed the A-site RDO changes in the double mutant (depletion of Not4 and *puf3Δ*) relative to the single mutant (depletion of Not4). For the mRNAs whose encoded proteins show the greatest reduction in protein aggregation in *not4Δpuf3Δ* relative to *not4Δ*, the A-site RDOs at cysteine TGC codons were significantly higher in the double mutant than in the single mutant (p=0.0153), whereas the A-site RDOs at the rare CGA and CGG arginine codons showed the greatest decrease (**Supplementary Figure S4A**).

These results suggest that RDOs at rare arginine codons fine-tune production of soluble proteins and is significantly impaired upon Not4 depletion but somewhat restored when Puf3 is additionally deleted. We thus classified mRNAs encoding proteins that aggregate in *not4Δ* according to their rare arginine codon content and evaluated the relative aggregation of the encoded proteins in *not4Δ* and in *not4Δpuf3Δ* mutants (**Figure 4A**). Compared to the single mutant, we observed less aggregation in the double mutant proportional to the amount of rare arginine codons. Since one might expect that longer mRNAs may have chances to bear more rare arginine codons and that mRNAs with rare arginine codons are more likely to be less expressed, we evaluated changes in A-site RDOs at rare arginine codons across broad expression level and mRNA lengths upon Not4 depletion (**Supplementary Figure S4B**) and upon Not4 depletion in the additional absence of Puf3 (**Supplementary Figure S4C**) and compared them to wild type. Upon Not4 depletion, A-site RDOs increased at rare arginine codons, and relatively decreased in the double mutant, for mRNAs at all expression levels and lengths. We then addressed whether the number of arginine codons had an impact on the relative solubility of mRNAs in the single *not4Δ* versus the double mutant *not4Δpuf3Δ* (**Figure 4B**). Indeed, mRNAs with more arginine codons tended to be less soluble in the double mutant compared to the single mutant compared to mRNAs with less arginine codons.

**Figure 4.**
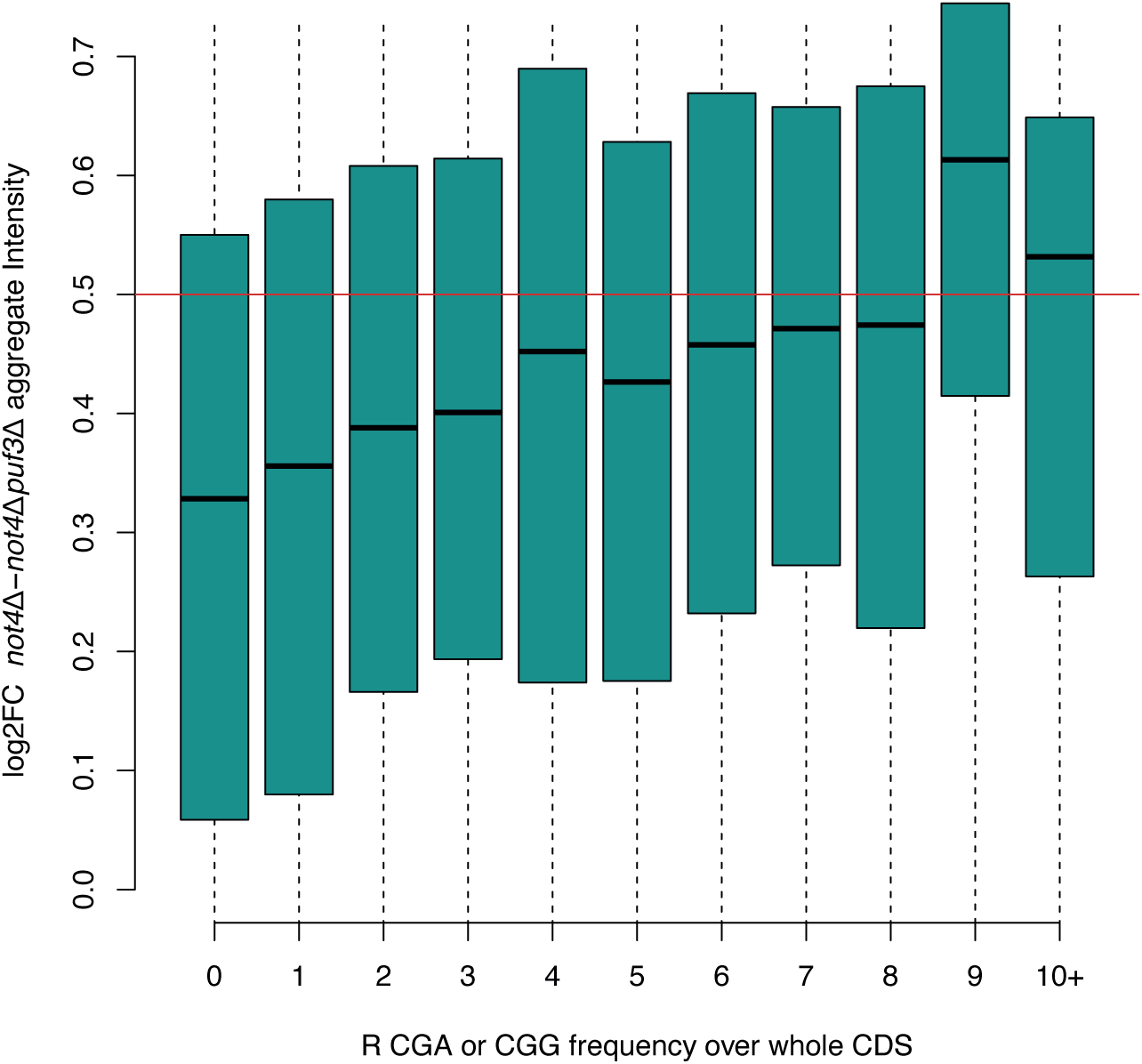
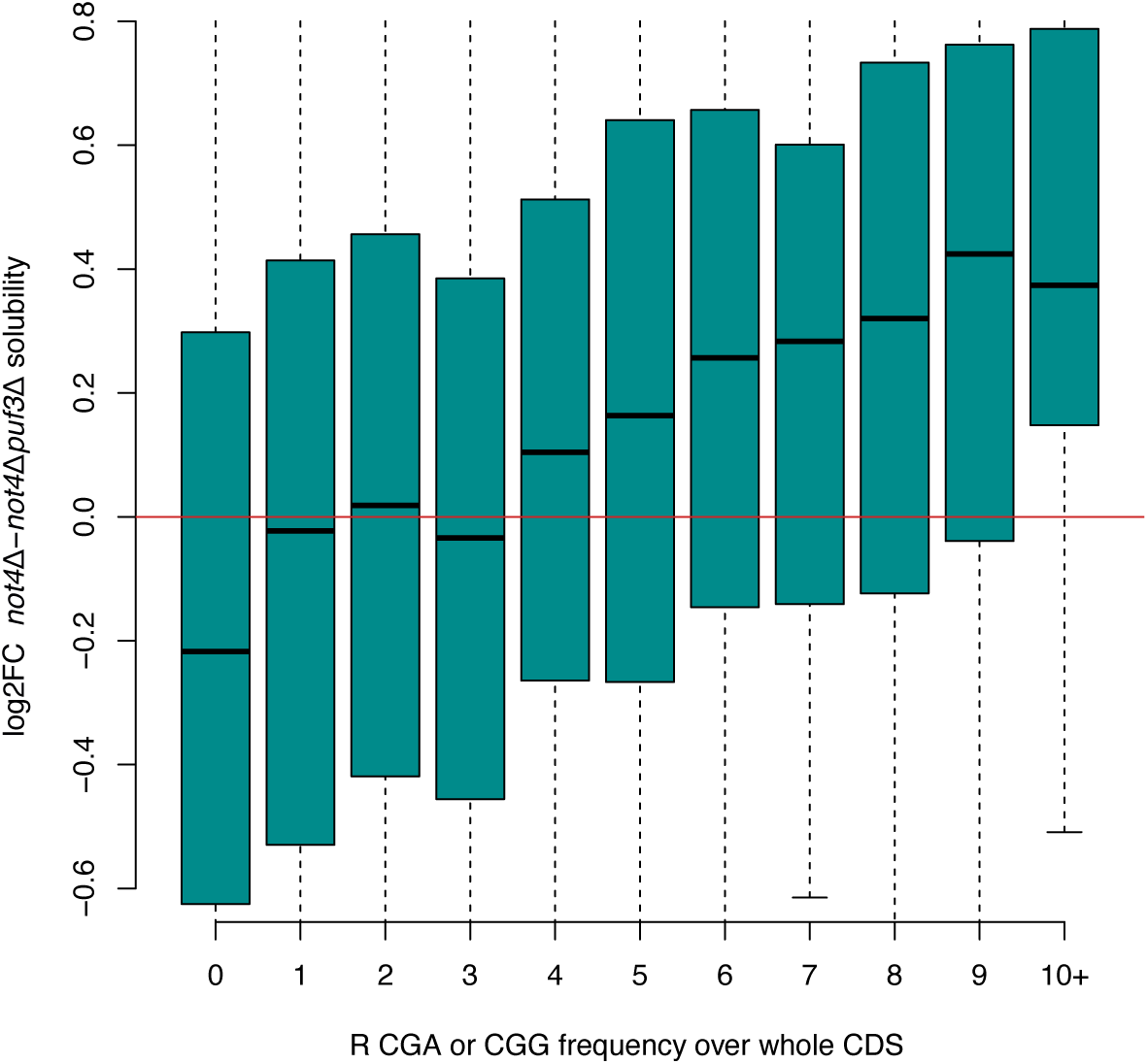
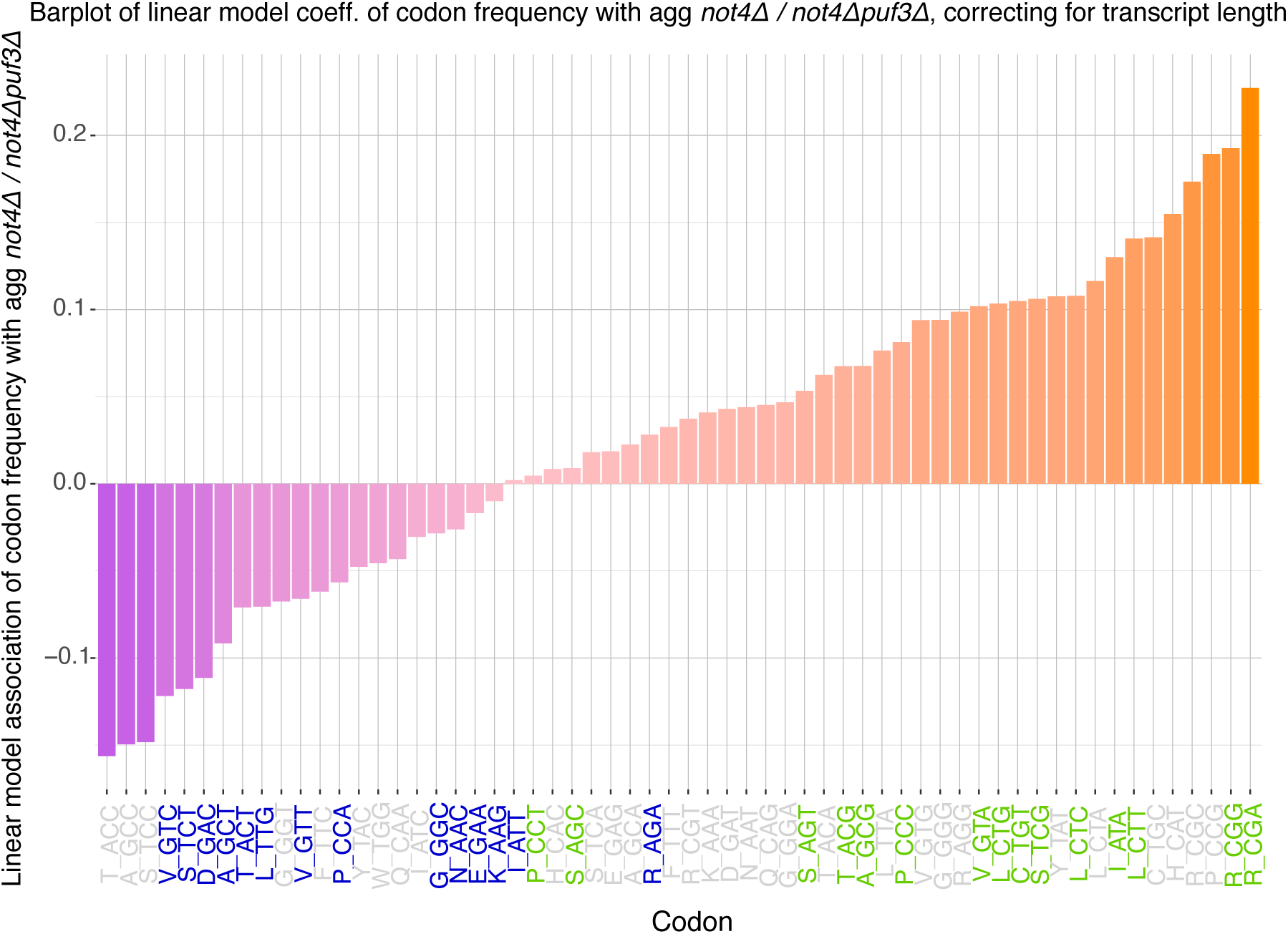
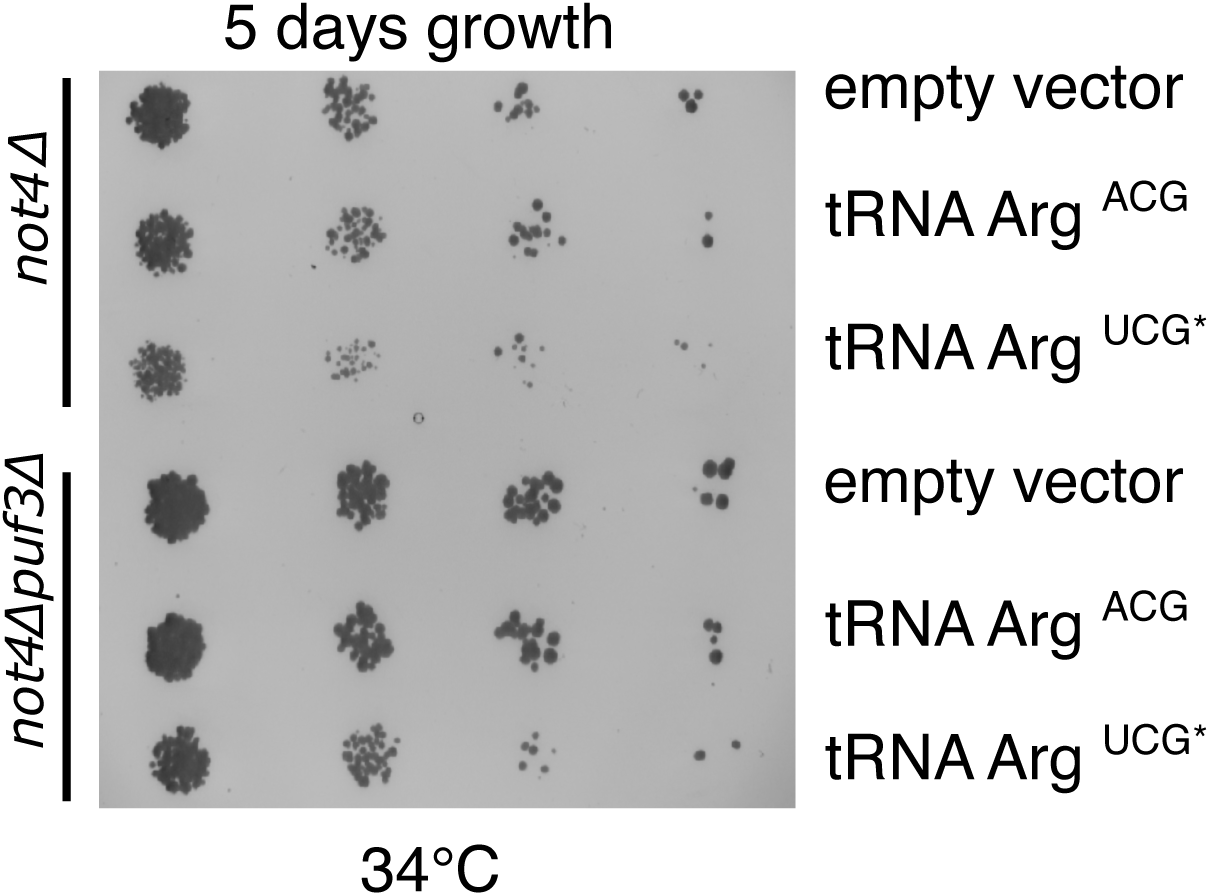
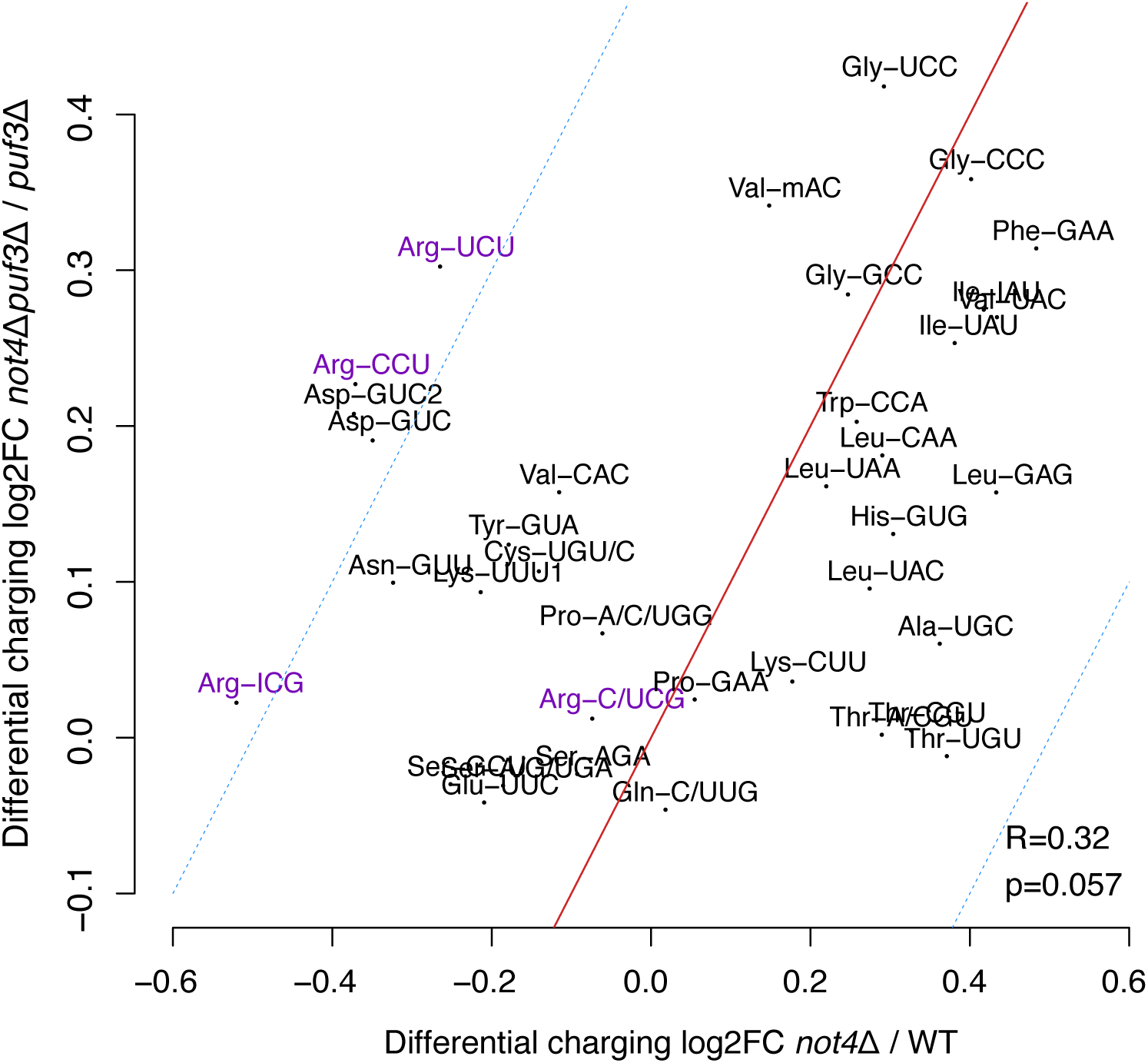
A-site RDOs at rare arginine codons upon Not4 depletion reversed by the deletion of Puf3 are associated with aggregation of encoded proteins. **A.** mRNAs whose encoded proteins were detected in *not4Δ* aggregates were classified according to the number of rare arginine codon content (CGG and CGA, x-axis) and changes in aggregation of encoded proteins in the single compared to the double mutant evaluated (y-axis). Results are shown by box plot analysis. **B**. As in **A** except that it is changes in mRNA solubility in the double compared to the single mutant that is compared (y-axis). **C**. Linear model of codon frequency in mRNAs that encode proteins that aggregate more in *not4Δ* than in *not4Δ puf3Δ,* correcting for transcript length. **D**. Spot assay (6 days growth) comparing the growth of wild type, *not4Δ*, *puf3Δ* and *not4Δpuf3Δ* cells transformed with a multicopy plasmid bearing the tRNA having the ACG anticodon, the tRNA having an anticodon that can directly decode CGA, or an empty vector (pRS416) as a control. **E**. Scatterplot comparing the change in tRNA charging in *not4Δ* compared to wild type (x-axis) or in *not4Δpuf3Δ* compared to *puf3Δ* (y-axis) (see **Supplementary Figure S4F**).

Finally, since both ribosome occupancy at the stop codon upon Not4 depletion and the rare arginine codon content of the mRNA is related to mRNA solubility and aggregation of encoded proteins in *not4Δ*, we analyzed whether mRNAs with increased ribosome occupancy at the stop codon also bear rare arginine codons. There was no correlation between the number of rare arginine codons and pausing at the stop codon. Equally, there was no significant change in periodicity comparing wild type to Not4 depletion with or without Puf3 when comparing transcripts with a 2-fold increase of ribosome occupancy at stop upon Not4 depletion relative to WT and those showing minor changes (-0.1 < log_2_FC < 0.1) (data not shown). Interestingly, the 500 top rare Arg codon-containing mRNAs encode proteins aggregating in *not4Δ*, but to a lesser extent in *not4Δpuf3Δ* (**Supplementary Table S6**). We also noticed that approximately one third of these mRNAs encode nuclear proteins. To consolidate these observations, we further investigated the role of codon content of mRNAs in the aggregation propensity of their encoded protein products in the single versus double mutant. This was done using a linear model of codon frequency in mRNAs against the relative aggregation of their encoded protein products in the single versus double mutant, correcting for transcript length. This analysis also revealed the most prominent role for rare arginine codons and additionally a bias related to codon optimality (**Figure 4C**).

Taken together, these results indicate that proteins produced from mRNAs with more rare arginine codons aggregate less in *not4Δpuf3Δ* compared to *not4Δ*, their encoding mRNAs are overall less soluble in the double mutant relative to the single mutant, and they exhibit lower A-site RDOs at the rare arginine codons upon Not4 depletion in the absence of Puf3 than in the presence of Puf3. These results indicate that Not4 and Puf3 oppositely regulate dwelling at rare arginine codons, associated with differences in mRNA solubility, and thereby prevent aggregation of encoded proteins.

### Puf3 contributes to aggregation of the arginyl-tRNA synthetase in *not4Δ*

Ribosomes with non-optimal codons in the A-site, including the non-optimal arginine codons, are enriched within the pool of ribosomes bound by Not5, in complex with Not4 (29), and they accumulate in cells lacking Not4 or Not5 (19) or when Not4 is rapidly depleted according to Ribo-Seq (see above). A possible explanation for this enrichment could be the reduced availability of the cognate aminoacyl-tRNAs in the mutant cells. We thus tested whether increasing the expression of tRNAs decoding rare arginine codons might improve growth of *not4Δ*. We overexpressed a tRNA^Arg^ACG that decodes the CGA codon. This tRNA is encoded by 6 genes in yeast and we have chosen the gene *YNCE0028C*. A34 in the ACG anticodon is modified to inosine, so that tRNA^Arg^ICG decodes CGU, CGC, and CGA, as described in (61). The yeast genome does not encode a tRNA^Arg^ with the UCG anticodon. Nevertheless, to not depend on the A34 modification, we also mutated the anticodon of tRNA^Arg^ACG and created a unique tRNA^Arg^UCG directly pairing to the CGA codon. The expression of the additional tRNA genes was toxic in *not4Δ,* particularly for the CGA modified tRNA, while the deletion of Puf3 suppressed the toxic effect at 34°C (**Figure 4D**).

In *not4Δ* we have previously observed that the amount of arginyl-tRNAs^Arg^ was generally reduced, while the levels of the uncharged tRNAs^Arg^ were unchanged (19), suggesting that charging of tRNAs^Arg^ might be impaired in *not4Δ*. We also noted that the arginine-tRNA synthetase (or ligase), encoded by *YDR341C*, catalyzing the charging of all tRNAs^Arg^, aggregated in *not4Δ*, while its aggregation in the double mutant was significantly reduced (**Supplementary Figure S4D**). This observation raised the question as to whether availability of arginyl-tRNAs^Arg^ might explain the differences in A-site RDOs at rare arginine codons. Therefore, using tRNA-tailored microarrays we measured charging levels of all tRNAs in *puf3Δ* or in *puf3Δnot4Δ* (**Supplementary Figures S4E and S4F** and **Supplementary Table S7**) and compared them to previous measurements in wild type and *not4Δ* (19). Notably, the level of arginyl-tRNA^Arg^ were improved or unchanged in the double *puf3Δnot4Δ* mutant compared to the single *puf3Δ* mutant (**Figure 4E**). Hence in the background of *puf3Δ*, we do not see that *not4Δ* results in decreased arginyl-tRNA^Arg^ as we do see in the presence of Puf3. Since the level of charged tRNAs is the most potent factor affecting A-site RDOs (62,63), it provides a plausible explanation for the lower RDOs observed by depleting Not4 in the absence of Puf3.

Charging of tRNAs depends also upon amino acid availability. Hence, we next focused on the *CPA1* gene - encoding a protein participating in arginine biosynthesis - whose expression is affected by arginine availability. This regulation occurs via production of a leader peptide from an upstream ORF (uORF) (64). Following, Not4 depletion, we observed accumulation of ribosome-protected fragments in the uORF, compatible with an excess of available arginine. This however was not detectably different in the double mutant or even exaggerated in cells lacking Puf3 (**Supplementary Figure S4G, left**). Consistent with a suppressant effect of uORF expression on the expression of the ORF, we see a commensurate significant loss in the CDS P-site PPKMs (P-sites Per Kilobase Million) upon Not4 depletion, with or without the additional deletion of Puf3 (**Supplementary Figure S4G, right**). Hence, availability of arginine is high in single and double mutants.

### Differences in the post-translational status of Puf3 and its interactome in cells with or without Not4

Our results above show that Puf3 affects translation elongation dynamics (in particular at rare arginine codons) alone or in combination with Not4. Thereby, it has a dominant negative impact in cells lacking Not4. We thus decided to purify Puf3 from both wild type and *not4Δ* cells. We used cells expressing a tagged version of Puf3 (*i.e.* Puf3-Tap-tag) and used the Tap-tag for affinity purification. Puf3-Tap-tag was slighty less expressed in *not4Δ* compared to wild type (**Supplementary Figure S5A**). It was very difficult to purify native Puf3 from cell lysates from *not4Δ* (**Supplementary Figure S5B**). However, when we mixed wild type cells expressing untagged Puf3 with *not4Δ* cells expressing Tap-tagged Puf3 (using equivalent OD_600_ cell counts of each), recovered the mixed cells by centrifugation and then broke them together with glass beads and lysis buffer, we obtained intact Tap-tagged-Puf3 protein (**Supplementary Figure S5B**). This suggests that Puf3-Tap-tag might be partially degraded during lysis of *not4Δ* cells, and this degradation *in vitro* can be prevented by extract mixing. We decided to use this mixing approach to purify Puf3-TAP-tag and compare its interactome in wild type and *not4Δ*. For purifications, all lysates originate from the same mixed pool of wild type and *not4Δ* cells, but Puf3 is Tap-tagged either only in the wild type cells (condition 1) or in the *not4Δ* cells (condition 2). We also used as a negative control mixed wild type cells and *not4Δ* cells lacking any tagged protein (condition 3). Since the lysates in all 3 conditions have the same total protein content, differences in the output of the purifications are not expected to be due to interactions in the test tube, but to represent differential purification of Puf3 interactors from the strain where Puf3 was tagged. For the analyses, we thus focused on proteins that were differentially present in the purifications of Puf3-Tap-tag from wild type and *not4Δ* and absent in the negative control.

In an initial pilot experiment, we successfully purified intact Tap-tagged Puf3 from both strains and determined that the Puf3 interactome significantly differed between the wild type and *not4Δ* (**Supplementary Figures S5C and S5D** and **Supplementary Table S8**) and that the difference in the Puf3 interactomes was not related to differences in expression or solubility of encoding mRNAs or synthesis of new proteins according to previous Ribo-Seq and RNA-Seq data comparing wild type, Not1 and Not4 depletion, and *not4Δ* (17,19) (**Supplementary Figure S5E**). This encouraged us to repeat the experiment with triplicate samples for a quantitative assessment of the Puf3 interactome in wild type and in *not4Δ* (**Supplementary Figure S5F**). Principal component analysis indicated that one of the Puf3 purifications from wild type cells was very different than the other 2 (**Supplementary Figure S5G**), hence we did not keep it for the further analyses. A volcano plot representing the comparative purifications in wild type and *not4Δ* is shown on **Figure 5A** and the comparison of the wild type and *not4Δ* Puf3 interactomes is presented in **Supplementary Table S9**. Overall, the differences in Puf3 interactomes in the wild type and *not4Δ* identified in the pilot experiment correlated with the differences in the subsequent triplicate experiment (R= 0.4, **Supplementary Figure S5H**). 355 proteins were more enriched in the Puf3 interactome from wild type cells compared to that in mutant cells (**Supplementary Table S10**). On the other hand, 42 proteins were more enriched in the Puf3 interactome from *not4Δ* compared to wild type cells (**Supplementary Table S11**). Of the 355 proteins more significantly present in the Puf3 interactome from wild type cells compared to *not4Δ* cells, 292 of these were also more significantly present in the purification from wild type cells expressing Tap-tagged Puf3 compared to the untagged control (**Supplementary Table S12**). A striking unexpected finding was the presence of Not3 in the Puf3 interactome from wild type cells but not from *not4Δ* cells. As for the 42 proteins more significantly present in the Puf3 interactome from *not4Δ* cells compared to wild type cells, all 42 were also more significantly present in the purification from *not4Δ* cells expressing Tap-tagged Puf3 compared to the untagged control (**Supplementary Table S13**).

**Figure 5.**
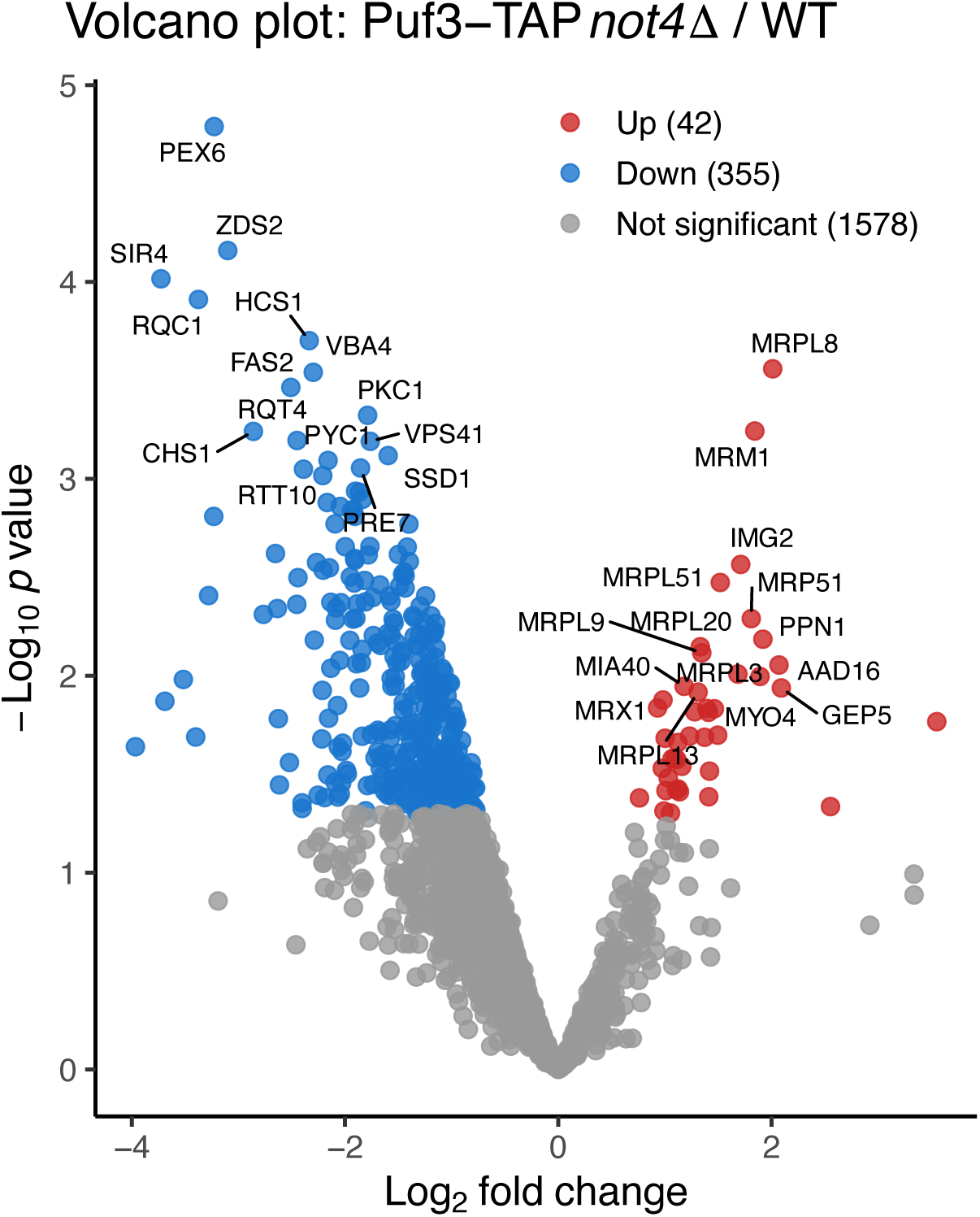
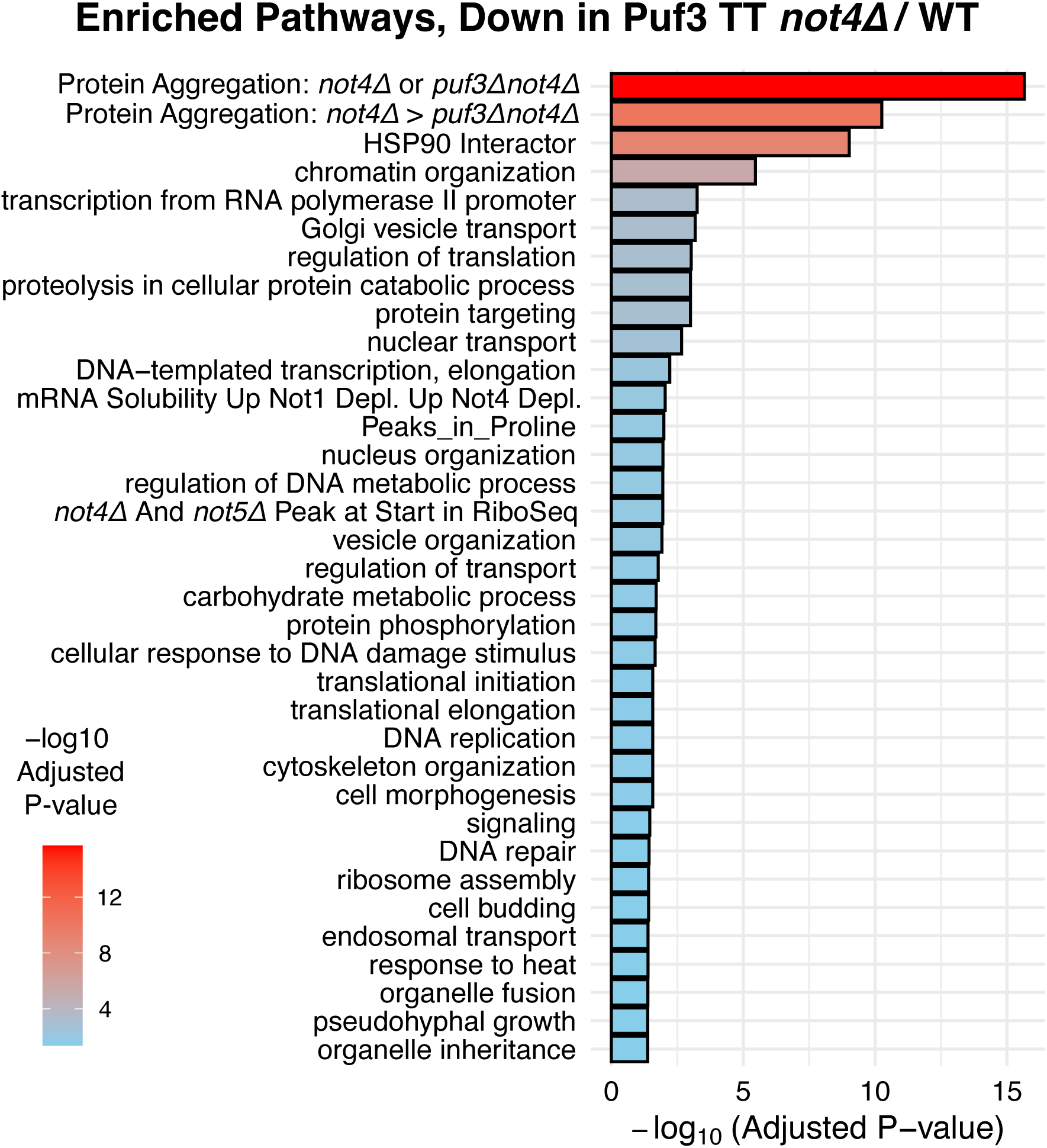
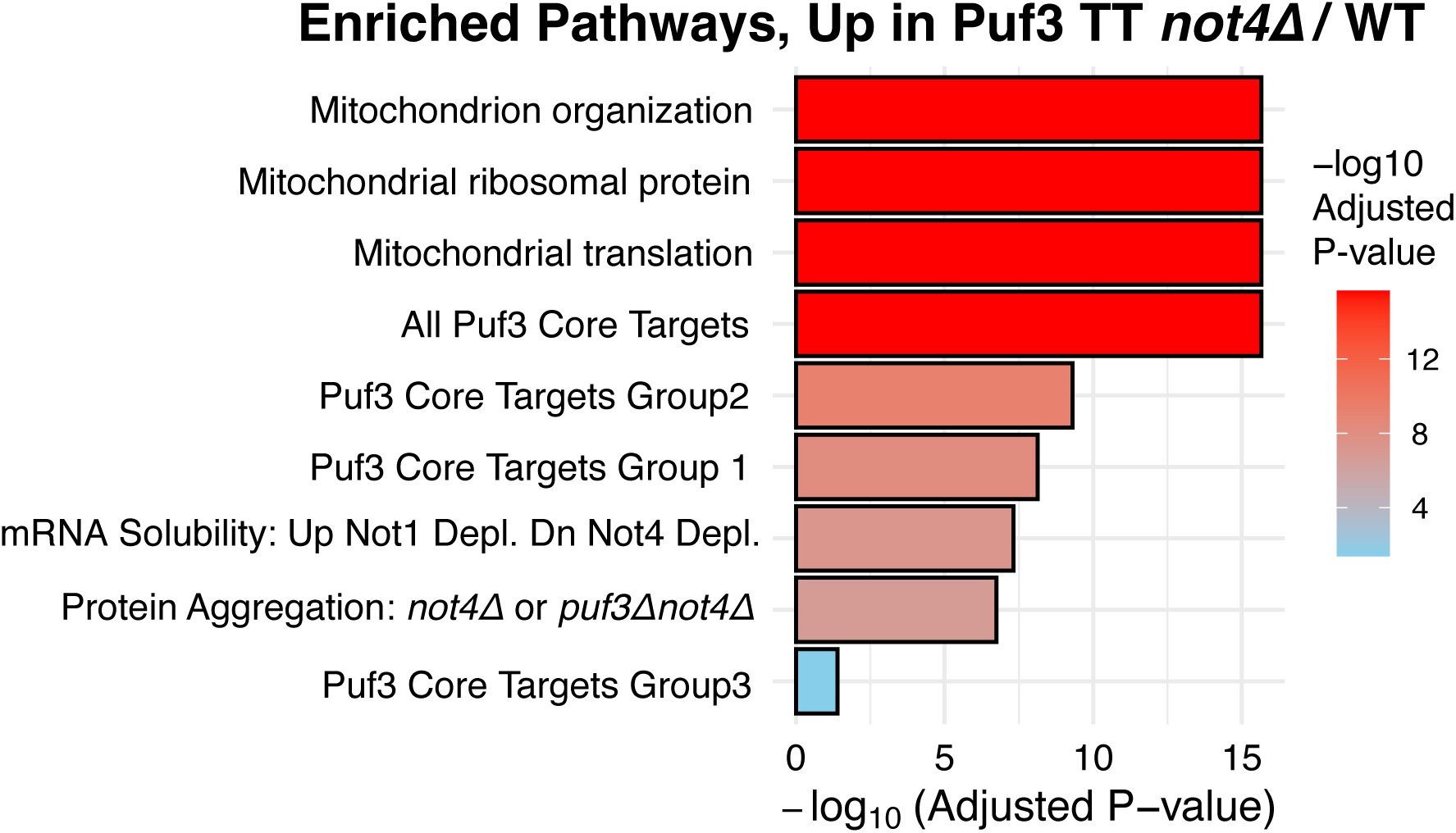
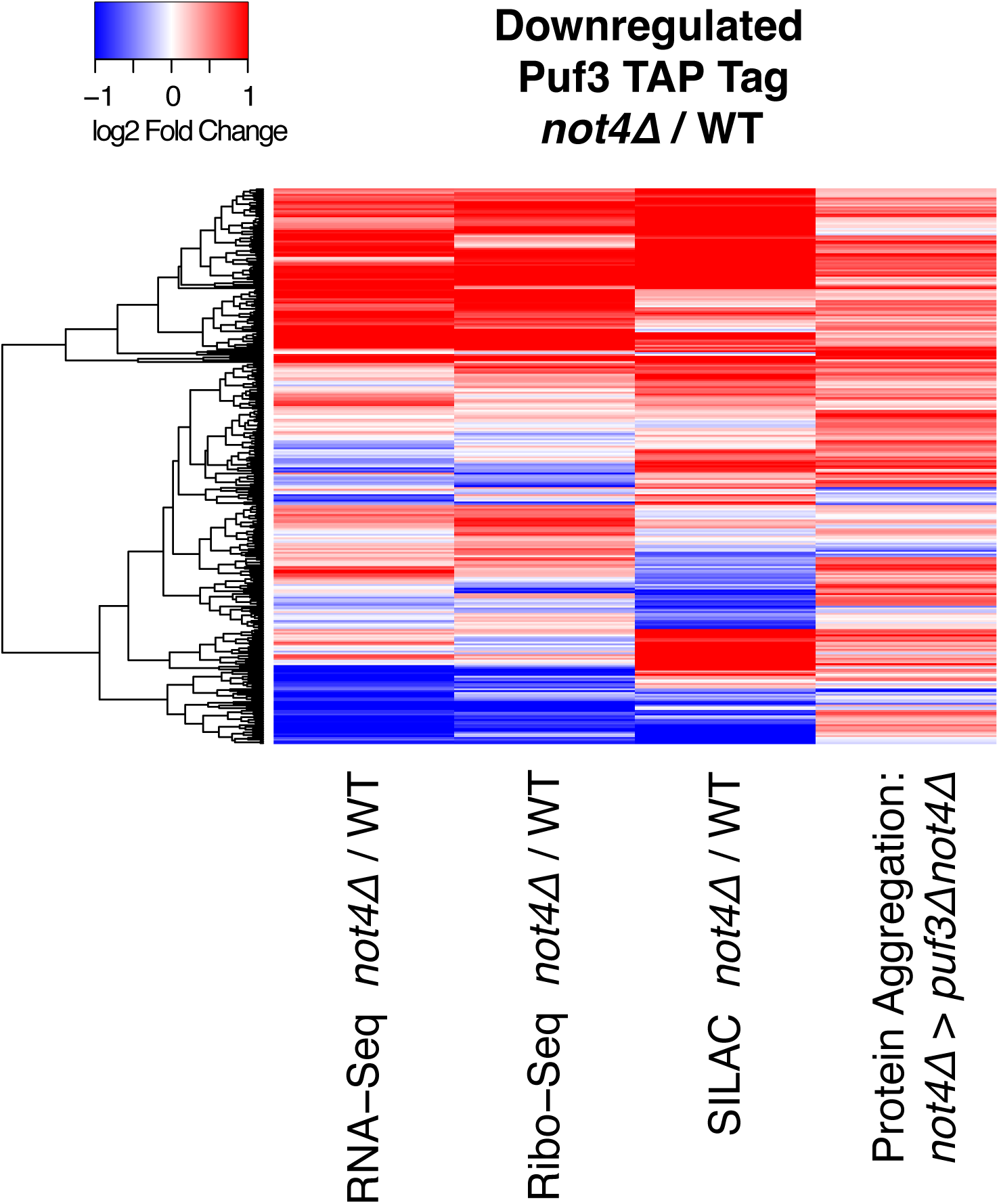
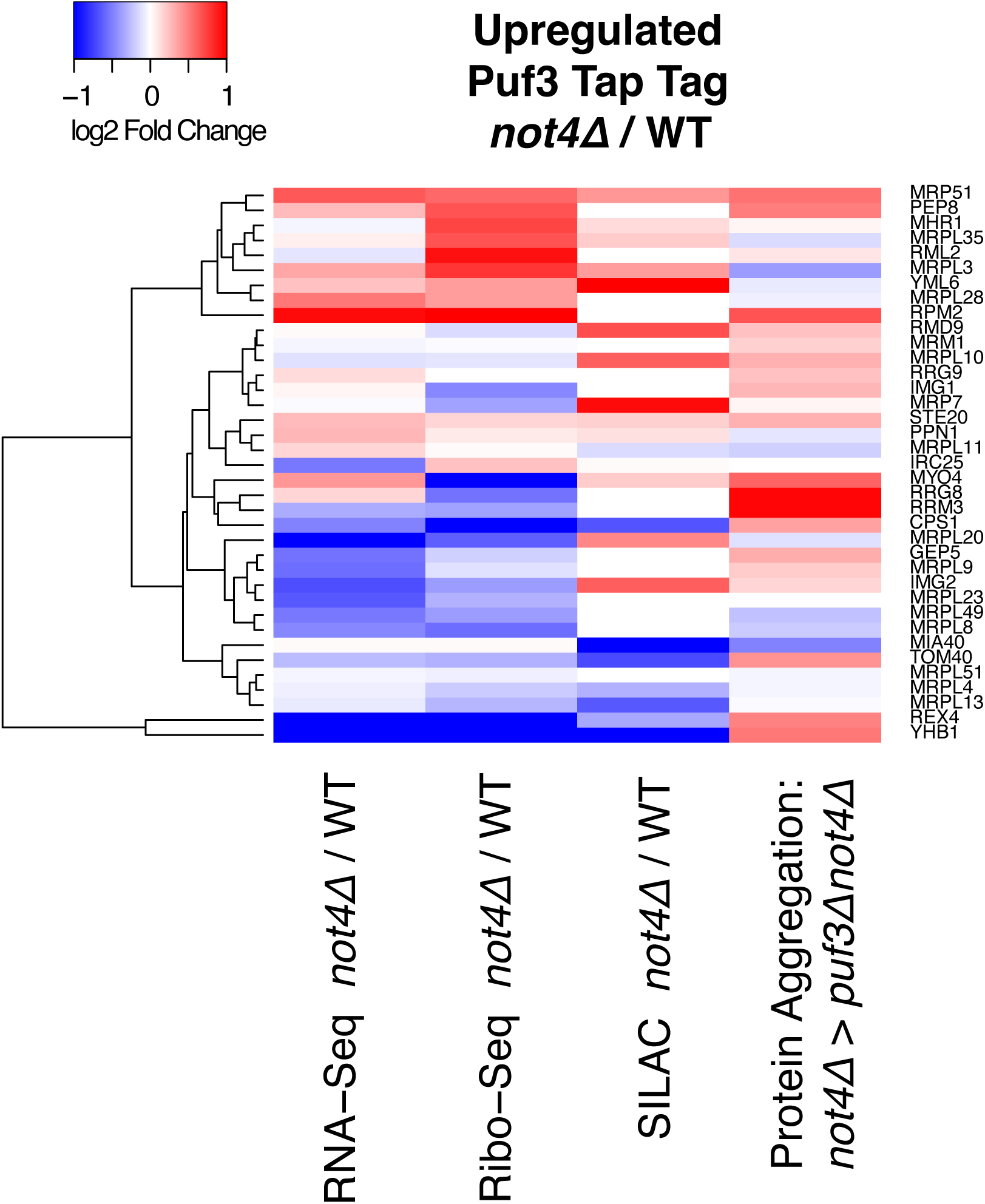
Characterization of the Puf3 interactome in *not4Δ* versus wild type. **A**. Volcano plot representing the comparative Puf3-TAP purifications in wild type and *not4Δ*. **B**. Enrichment analysis for the Puf3 interactome in wild type over *not4Δ*. **C.** Enrichment analysis for the Puf3 interactome in *not4Δ* over wild type. **D**. Changes in expression extracted from previous data sets (19) corresponding to the proteins that are present in the Puf3 interactome from wild type cells but not mutant cells. **E**. Same as in panel **D** but corresponding to the proteins present in the interactome from mutant cells but not wild type cells.

GO-term analysis of the proteins enriched in the Puf3 interactome from wild type cells compared to that of *not4Δ* revealed mainly proteins that tend to aggregate in the absence of Not4, Hsp90 interactors and proteins involved in chromatin organization (**Supplementary Table S14** and **Figure 5B**). Instead, for the proteins enriched in the interactome from *not4Δ* cells, they were mostly mitochondrial proteins and proteins encoded by mRNA targets of Puf3 (**Figure 5C** and **Supplementary Table S15**). It was particularly interesting to note the presence of Tom40 in the interactome from mutant cells only, as Tom40 is the central channel component of the TOM complex, that functions as the entry gate for most mitochondrial proteins (reviewed in (65)).

In both the wild type and *not4Δ* Puf3 interactomes (**Supplementary Table S16** for the wild type and **Supplementary Table S17** for the *not4Δ*), known associated factors of Puf3, whether direct interactors or proteins whose interaction is mediated by mRNA, such as the polyA binding protein Pab1 (66) and the Hrr25 kinase (67), as well as mitochondrial proteins, consistent with the known importance of Puf3 for mitochondrial translation (35,40,45,68), were significantly enriched. It is relevant to note that Not4 was identified as a protein significantly enriched in the Puf3 interactome from *not4Δ* cells compared to the untagged control, despite the absence of Not4 in the strain which had the tagged Puf3. Not4 was however in the mixed lysate from which the immunoprecipitation was performed. This reveals that interactions can indeed happen in the test tube, supporting our approach to focus only on proteins differentially purified from wild type or *not4Δ* cells. We compared the differential expression of proteins differentially present in the Puf3 interactomes from wild type and *not4Δ* using our previous SILAC, Ribo-Seq and RNA-Seq data in wild type and *not4Δ* (shown on **Figure 5D** and **Figure 5E** for the proteins for which we had the information from all 3 data sets (19)) and noted that there was no correlation between differential expression and differential co-purification from wild type or *not4Δ*.

Previous work has indicated that the different roles of Puf3 during fermentation and respiration, in particular Puf3’s function as a negative regulator of mRNAs encoding mitochondrial proteins, are regulated through phosphorylation of its N-terminal domain by the Hrr25 kinase (67) and a Puf3 phosphomutant derivative was reported to be dominant negative during respiration (40). We hence compared Puf3’s phosphorylation in wild type and *not4Δ*. We detected several Puf3 phosphopeptides, but importantly, the peptide (559–568) was heavily phosphorylated in the mutant and significantly less in the wild type (**Supplementary Table S18**). Hence, the phosphorylation of Puf3 is different in wild type and *not4Δ* cells growing in glucose.

Taken together these results determine that both Puf3’s interactome and post-translational status are modified in cells lacking Not4.

## DISCUSSION

In this work we determine that Puf3 contributes to protein aggregation and temperature sensitivity in cells lacking Not4. Not only does the deletion of Puf3 suppress partially temperature sensitivity and protein aggregation in yeast cells lacking Not4, but also Puf3 has a dominant negative impact on growth of *not4Δ* cells. This negative impact of Puf3 in *not4Δ* correlates with altered post-translational modifications of Puf3 and an altered Puf3 interactome in the mutant. Furthermore, we show that inappropriate translation elongation dynamics, in particular at rare arginine codons, associated with aberrant mRNA solubility, is responsible for aggregation of new proteins due to aberrant Puf3 in *not4Δ*. Besides its role in *not4Δ*, our work also uncovers a codon-specific impact of Puf3 deletion on translation elongation.

We examined the Puf3 interactome in wild type and *not4Δ* cells, focusing on the differences between wild type and mutant interactomes. It was notable that the Puf3 interactors specific to the wild type cells were enriched for proteins that tend to aggregate in *not4Δ*. These findings suggest that Not4 is a “guardian” of Puf3’s function that if left unattended can compromise protein homeostasis. Exactly how Not4 protects such a Puf3 regulatory axis is unclear. It could occur by Not4 controlling enzymes that post-translationally modify Puf3 or by regulating components of the Puf3 interactome and thereby indirectly Puf3’s phosphorylation status. In addition, Not4, like the other Not proteins, is present in condensates (19), as has also been proposed for Puf3 in glucose (40), hence availability of proteins for interactions depending upon the status of the condensates is a possibility.

We were able to determine that for some mRNAs, changes in mRNA solubility (whether increased or decreased) directly correlated with changes in aggregation of encoded proteins in *not4Δ*, because of the combined suppression by *puf3Δ* of protein aggregation and mRNA solubility. In this category are mRNAs with rare arginine codons, as well as a number of Puf3 target mRNAs, and mRNAs encoding Hsp90 interactors. For these particular mRNAs, our results indicate that protein homeostasis is improved when the mRNAs are less soluble. It seems likely that this is because the mRNAs must be targeted and/or condensed appropriately for appropriate translation elongation dynamics and correct protein targeting, folding and/or assembly. For these mRNAs, only a low number of mRNAs (90), Puf3 and Not4 seem to work in the same direction. It was not unexpected to identify mitochondrial mRNAs and Puf3 targets in this category, since the importance of both Puf3 and Not4 for targeting of mitochondrial mRNAs to mitochondria has already been described (20,69). It was, however, much more unexpected for Hsp90 interactors and new for mRNAs with rare arginine codons.

Other mRNAs instead are less soluble in *not4Δ*, corresponding to higher aggregation of encoded proteins, and this can be improved by the deletion of Puf3. We imagine that in the case of these mRNAs, some of them might become less soluble in the form of Ribosome Nascent Chains (RNCs) with misfolded nascent chains. Either Puf3 has a negative impact on the elongation dynamics in the absence of Not4, or Puf3 has an inhibitory effect on clearance of the aggregated RNCs in *not4Δ*. In this context it was interesting to find components of the Ribosome Quality control (RQC) machine as interactors of Puf3 only from wild type cells. mRNAs in this latter category include several chaperones such as Zuo1 and the Hsp90 proteins (Hsc82 and Hsp82). Obviously if chaperone production is impaired in *not4Δ* and improved in the double mutant, this will impact aggregation of many other proteins that are chaperone dependent, both co-translationally and post-translationally. The translation termination factor eRF3 (encoded by *SUP35*) could be such a protein since it’s aggregation in *not4Δ* is reduced the deletion of Puf3. In parallel to this, we observe that the absence of Puf3 reduces the increased pausing at the stop codon in *not4Δ*. For any downstream targets of chaperone function, when comparing *not4Δ* and *not4Δpuf3Δ*, one does not expect to find a correlation between mRNA solubility and protein aggregation, and this is indeed what is observed for the majority of the mRNAs encoding proteins that aggregate in *not4Δ*. An overall defect in chaperone capacity of cells lacking Not4 is very consistent with our observation that proteins tend to undergo proteolysis in extracts from *not4Δ* but that this can be compensated for by mixing *not4Δ* cells and wild type cells prior to cell lysis.

Puf3 is known to interact with the Ccr4-Not complex and accelerate deadenylation during fermentation (32), but whether this known interaction might result in modulation of other functions of the Ccr4-Not complex has not been explored. In this report we observed that indeed, Puf3 contributed to inverse changes in A-site RDOs upon Not1 and Not4 depletion, not only for its target mRNAs but generally for all mRNAs, and this in a codon-optimality related manner, a role of Puf3 that doesn’t seem to involve recognition of its target mRNAs at a consensus sequence on the mRNA, since it is observed for all mRNAs and not only its targets. An interesting observation in this regard is that the changes in A-site RDOs in *puf3Δ* correlate with those upon depletion of the eIF5A translation initiation and elongation factor. We identified eIF5A in the pilot purification of Puf3, but not consistently in the next triplicate purifications. From previous work we know that changes in A-site RDOs upon depletion of eIF5A correlate inversely with those in *not5Δ* versus wild type. Hence an alternative possibility is that in the absence of Puf3, Not5 competes more effectively compared to eIF5A for ribosomal E site binding. Potentially this could be because Puf3 is not there to interact with the Ccr4-Not complex making more Not5 available to compete with eIF5A. An alternative maybe more likely possibility is that in the absence of Puf3, Not3 in addition to Not5 is available to bind to the ribosomal E site. Indeed, we observed that Puf3 from wild type cells interacts with Not3, compatible with its previously described interaction in *S.pombe in vitro* (11), but not Puf3 from mutant cells. Notably, the N-terminal domain of Not3 is homologous to that of Not5 that is the one that binds the ribosomal E site. Maybe in *not4Δ* cells when Puf3 is present but no longer interacting with Not3, available Not3 might improve the A-site RDO changes due to the absence of Not4, if hypothetically the absence of Not4 has a negative impact on the association of Not5 with the ribosomal E site. A role of Puf3 for regulation of Not3 and A-site RDOs is certainly an exciting avenue to investigate in the future.

Regardless of the exact mechanisms linking Puf3, Not3, Not4 and Not5, this work clearly supports the idea that Puf3 has a broad role in regulation of translation elongation dynamics that impacts all mRNAs and not only its target mRNAs, and that is intrically connected to the functions of Not3, Not4 and Not5. Moreover, the findings of this work indicating that Puf3 contributes to some, but not all protein aggregation in *not4Δ*, leaves open the door to identifying other Ccr4-Not interacting RBPs that together with Puf3, might mediate the global loss of protein homeostasis in *not4Δ*.

## ACKNOWLEDGEMENTS

We thank our colleagues from the Collart laboratory, in particular Rachel Porcelli who extensively participated in this project and constructed plasmids and Luciana Romano for technical assistance. We thank the R2P platform for Ribo-seq experiments and the Proteomics Core Facility, Faculty of Medicine, University of Geneva, Switzerland for the mass spectrometry experiments. RNA-seq and Ribo-Seq sequencing were performed at the iGE3 Genomics platform of the University of Geneva (https://ige3.genomics.unige.ch).

## AUTHOR CONTRIBUTIONS

Conceptualization: LA, MAC

Data curation: MAC, GEA

Formal analysis:LA, SC, MAC, CP, ZI, GEA

Funding acquisition: MAC, SC, VP, ZI

Investigation:LA, MAC, SC, SH

Methodology: LA, MAC, SC, SH, VP, CP, ZI, OOP, GEA

Project administration: LA, MAC

Resources: MAC, GEA

Software: GEA

Supervision: MAC, VP

Validation: LA, MAC, OOP

Visualization: LA, GEA

Writing – original draft: MAC

Writing – review & editing: all authors

## SUPPLEMENTARY DATA

Supplementary Data are available at NAR online.

## DATA AVAILABILITY

The new Ribo-Seq data is accessible under GSE303644, the mass spectrometry data in the Pride partner repository under PXD069853 and the tRNA microarray data are accessible under the accession number GSE305765 in the Gene Expression Omnibus (GEO) database.

The *not4Δ* RNA-Seq and degron RNA-Seq are already online: https://www.ncbi.nlm.nih.gov/geo/query/acc.cgi?acc=GSE168290 https://www.ncbi.nlm.nih.gov/geo/query/acc.cgi?acc=GSE193912

All code used in the manuscript is available: https://github.com/georgeallenunige/NotTranslation

## SUPPLEMENTARY TABLES

**Supplementary Table S1.** Strains, plasmids and oligo list

**Supplementary Table S2**. Changes in protein aggregation in *not4Δ* versus *not4Δ puf3Δ*

Differences in identification of proteins in aggregates from *not4Δ* versus *not4Δpuf3Δ* are expressed in log_2_ fold ratios.

**Supplementary Table S3.** GO-terms enriched for proteins more aggregated in *not4Δpuf3Δ* versus *not4Δ*

**Supplementary Table S4**. GO-terms enriched within the proteins less aggregated in *not4Δpuf3Δ* than in *not4Δ*

**Supplementary Table S5. GO-terms enriched for mRNAs showing the increase of pausing at the stop codon upon depletion of Not4**

**Supplementary Table S6. GO-terms enriched within the 500 top rare Arg-codon containing mRNAs**

**Supplementary Table S7.** Expression and charging of tRNAs in wild type, *not4Δ*, *puf3Δ* and *not4Δpuf3Δ*

**Supplementary Table S8**. Identification of proteins co-purified with Puf3-Tap-tag in wild type or *not4Δ* and from a non-tagged strain as a control. Proteins identified in the different purifications from a pilot experiment are expressed with the protein probabilities.

**Supplementary Table S9.** Comparison of the wild type and Puf3 interactomes with the retained samples of the experiment in triplicate, dropping sample 6.

**Supplementary Table S10.** Proteins enriched in the Puf3 interactome from wild type cells compared to that in mutant cells.

**Supplementary Table S11.** Proteins enriched in the Puf3 interactome from *not4Δ* compared to wild type cells.

**Supplementary Table S12.** Proteins enriched in the Puf3 interactome from wild type cells compared to that in mutant cells and enriched in the specific Puf3 affinity purification compared to the no-tag control.

**Supplementary Table S13.** Proteins enriched in the Puf3 interactome from *not4Δ* cells compared to that from wild type cells and enriched in the specific Puf3 affinity purification compared to the no-tag control.

**Supplementary Table S14.** GO-term analysis of the proteins enriched in the Puf3 interactome from wild type cells compared to that of *not4Δ,* retaining only proteins enriched in the specific affinity purification compared to the no-tag control.

**Supplementary Table S15.** GO-term analysis of the proteins enriched in the Puf3 interactome from *not4Δ* cells compared to that from wild type cells, retaining only proteins enriched in the specific affinity purification compared to the no-tag control.

**Supplementary Table S16.** Proteins significantly present in the Puf3-TAP purification from wild type cells compared to the untagged control.

**Supplementary Table S17.** Proteins significantly present in the Puf3-TAP purification from *not4Δ* cells compared to the untagged control.

**Supplementary Table S18.** Puf3 phosphorylation in wild type and in *not4Δ.*

---

**Supplementary Figure S1.**
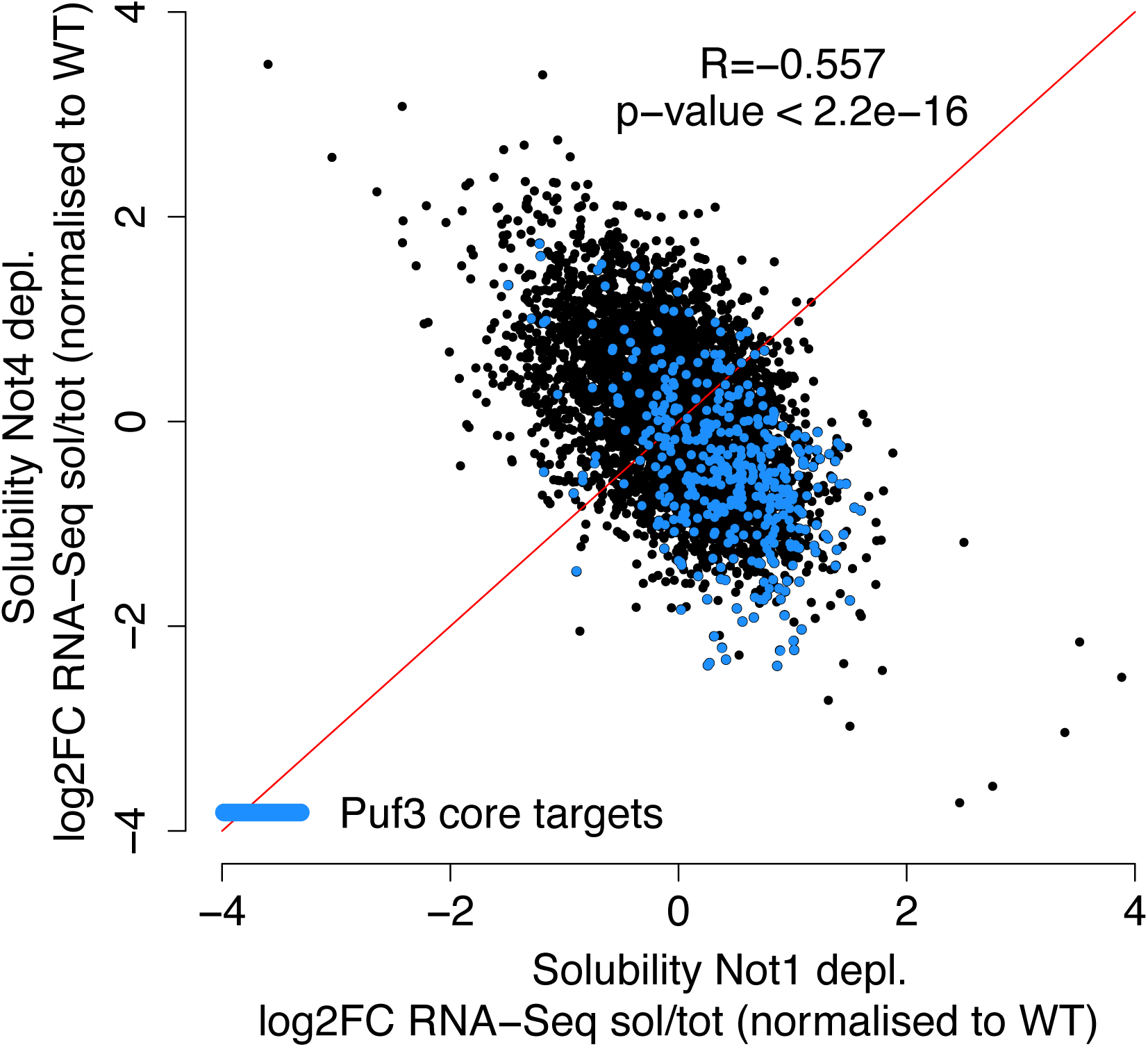
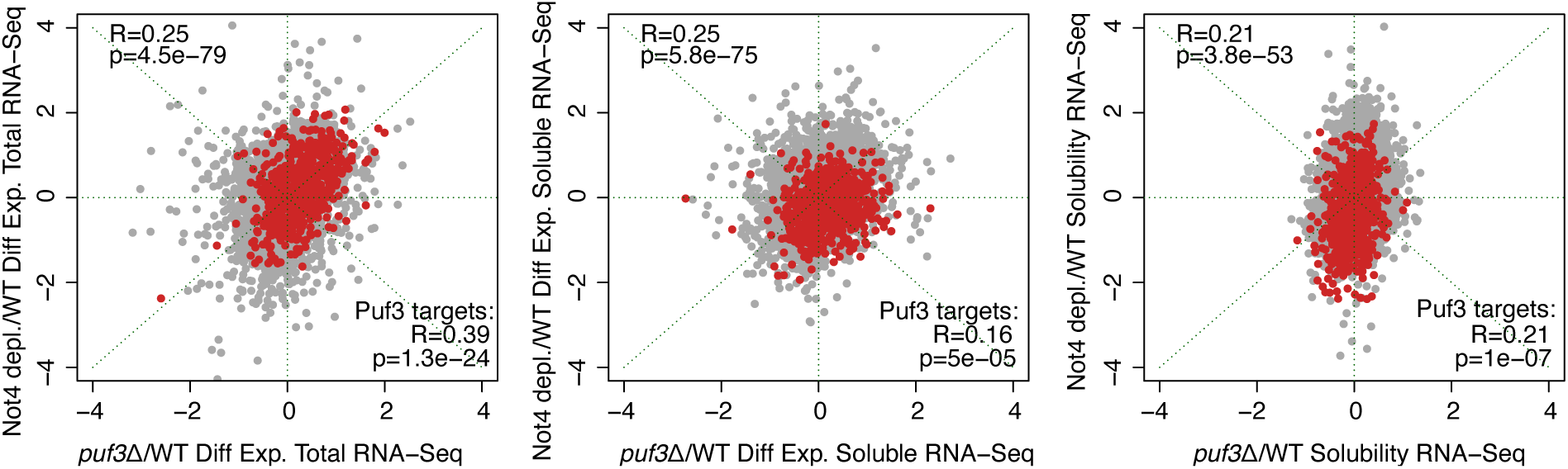
Puf3 targets are enriched within mRNAs less soluble upon Not4 depletion but more soluble upon Not1 depletion. **Related to** Figure 1. **A.** Scatterplot showing inverse solubility changes in mRNA solubilities upon Not1 (x-axis) and Not4 (y-axis) depletion (17). Puf3 target mRNAs are indicated in blue**. B.** Scatterplot showing changes in levels of total RNAs (left panel), soluble RNAs (middle panel) or solubility (right panel) of all mRNAs (in grey, R value upper left corner of each panel) or Puf3 target mRNAs (in red, R value lower right corner of each panel) comparing *puf3Δ* (x-axis) and the Not4 depletion (y-axis).

**Supplementary Figure S2.**
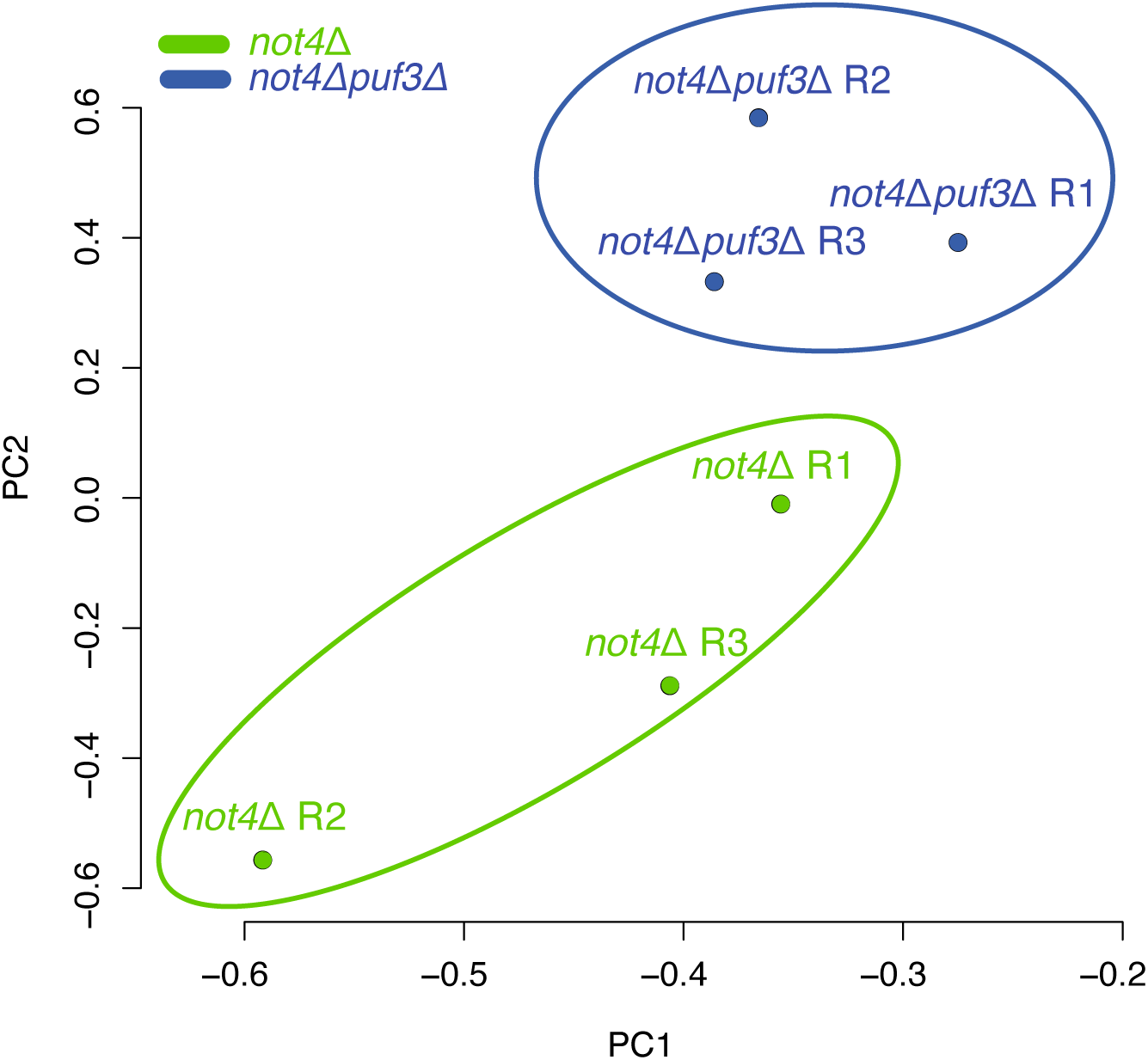
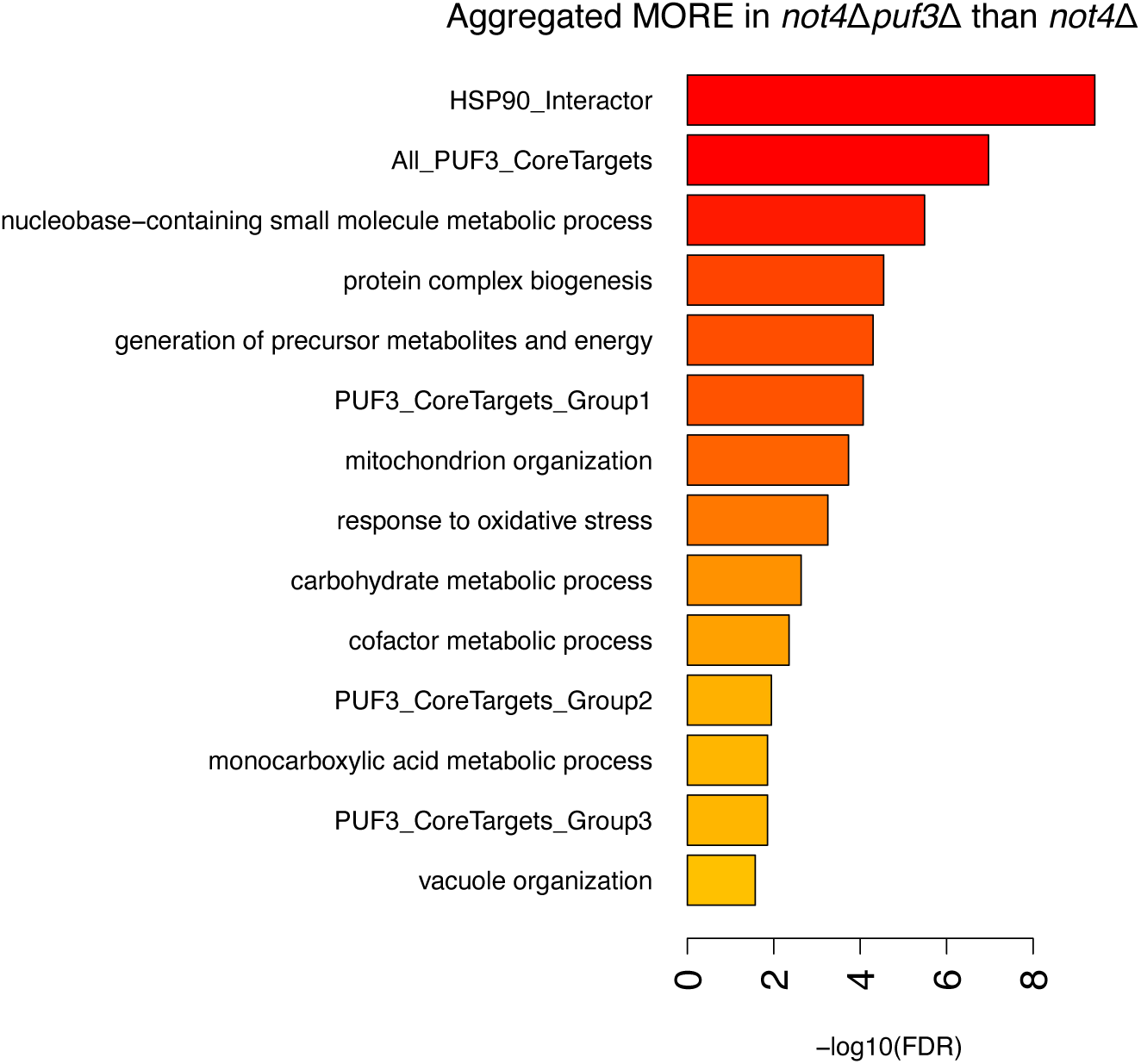
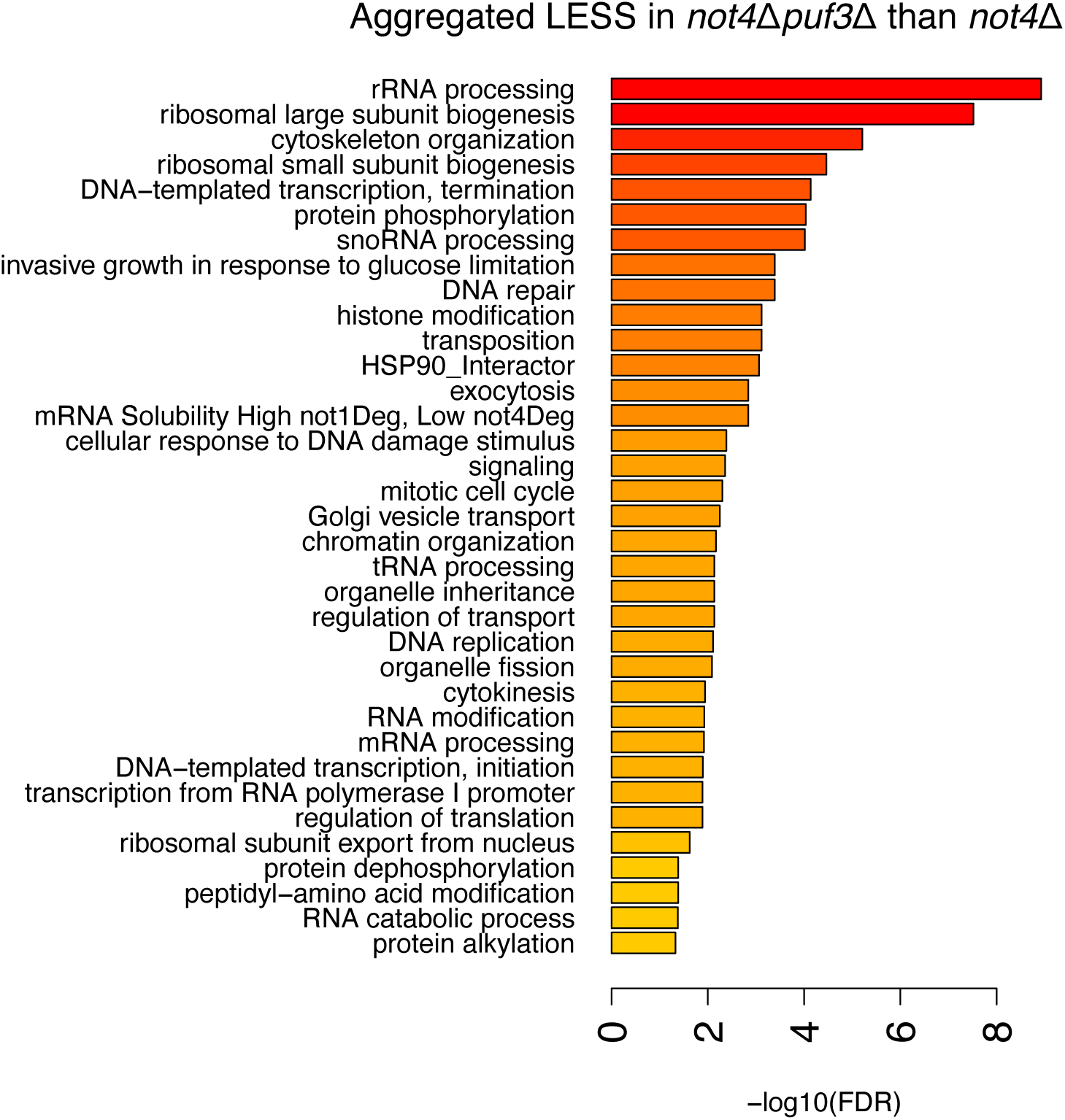
Analysis by quantitative mass spectrometry of proteins in aggregates in *not4Δ* and *not4Δpuf3Δ*. Related to Figure 2**. A.** PCA analysis of the triplicate samples of *not4Δ* and *not4Δpuf3Δ* that were analyzed by quantitative mass spectrometry. **B-C**. GO-term analysis of proteins that are either: more present in aggregates in the double mutant **(B**); or less present in aggregates in the double mutant **(C**).

**Supplementary Figure S3.**
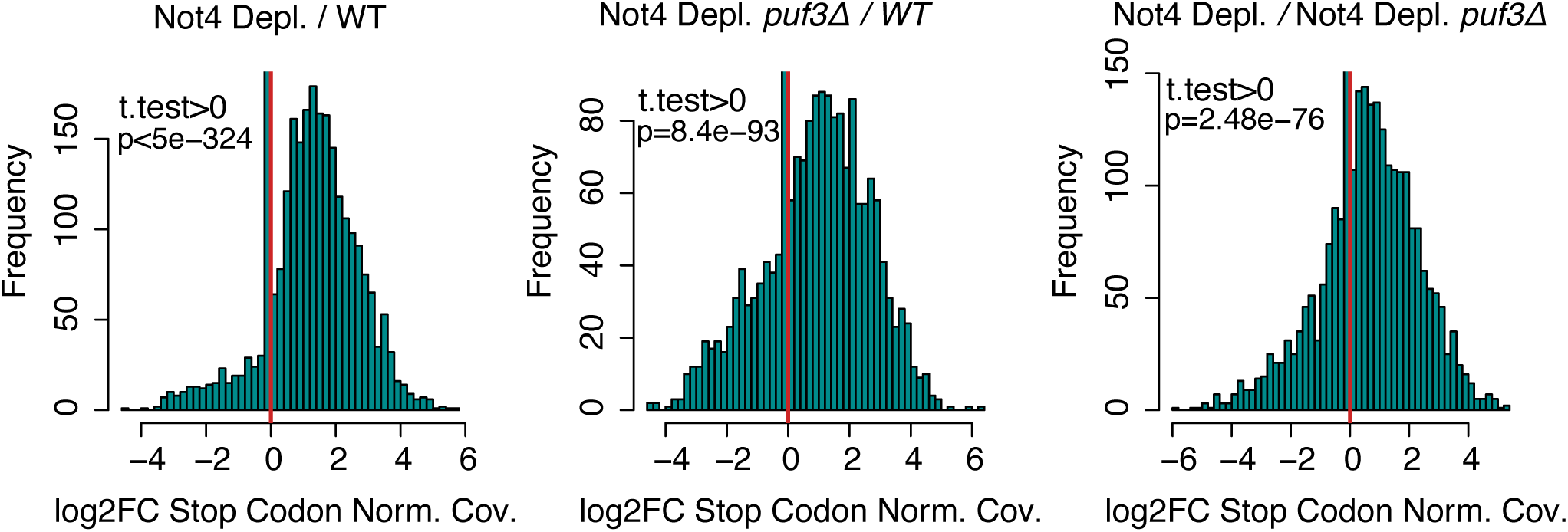
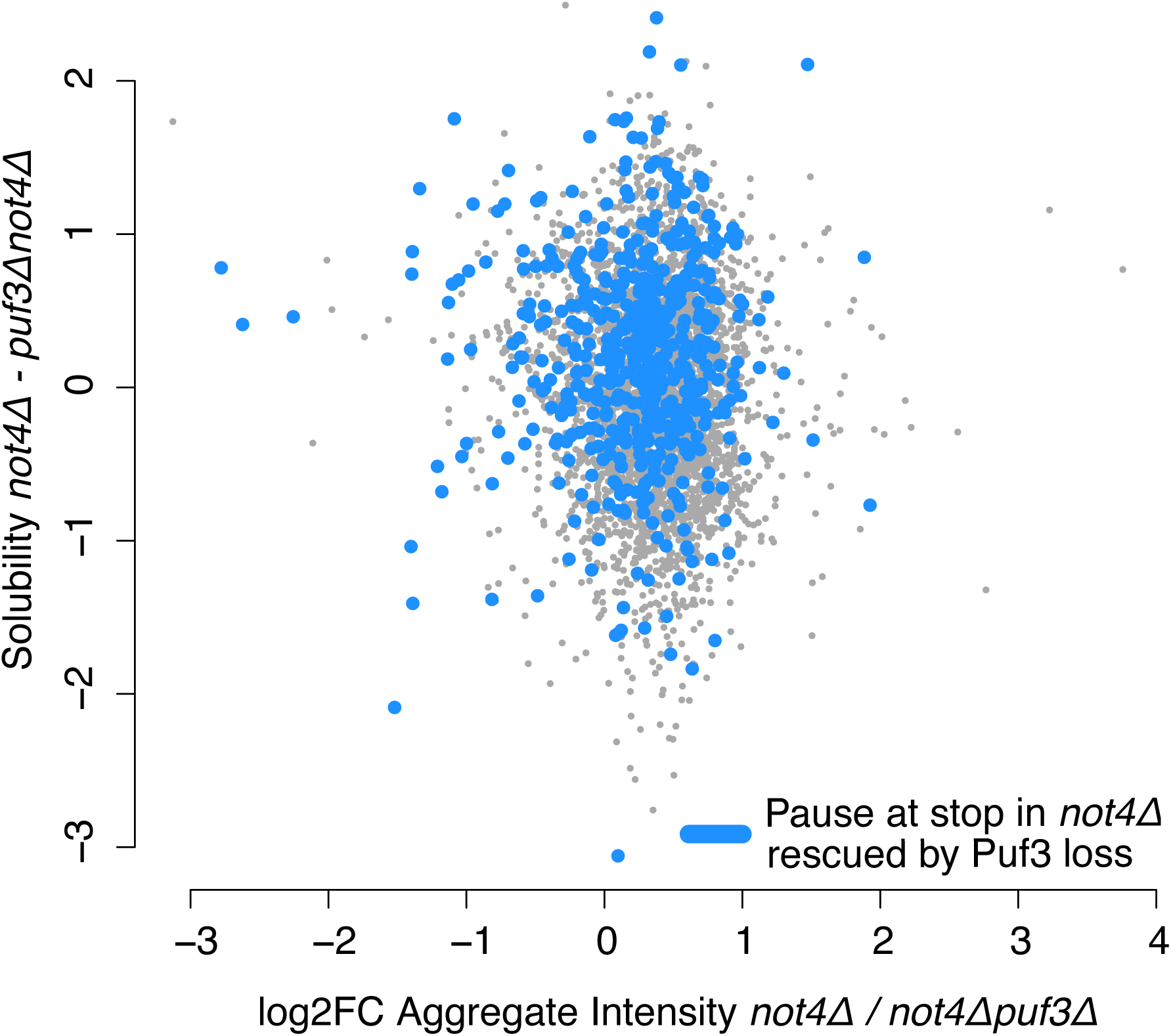
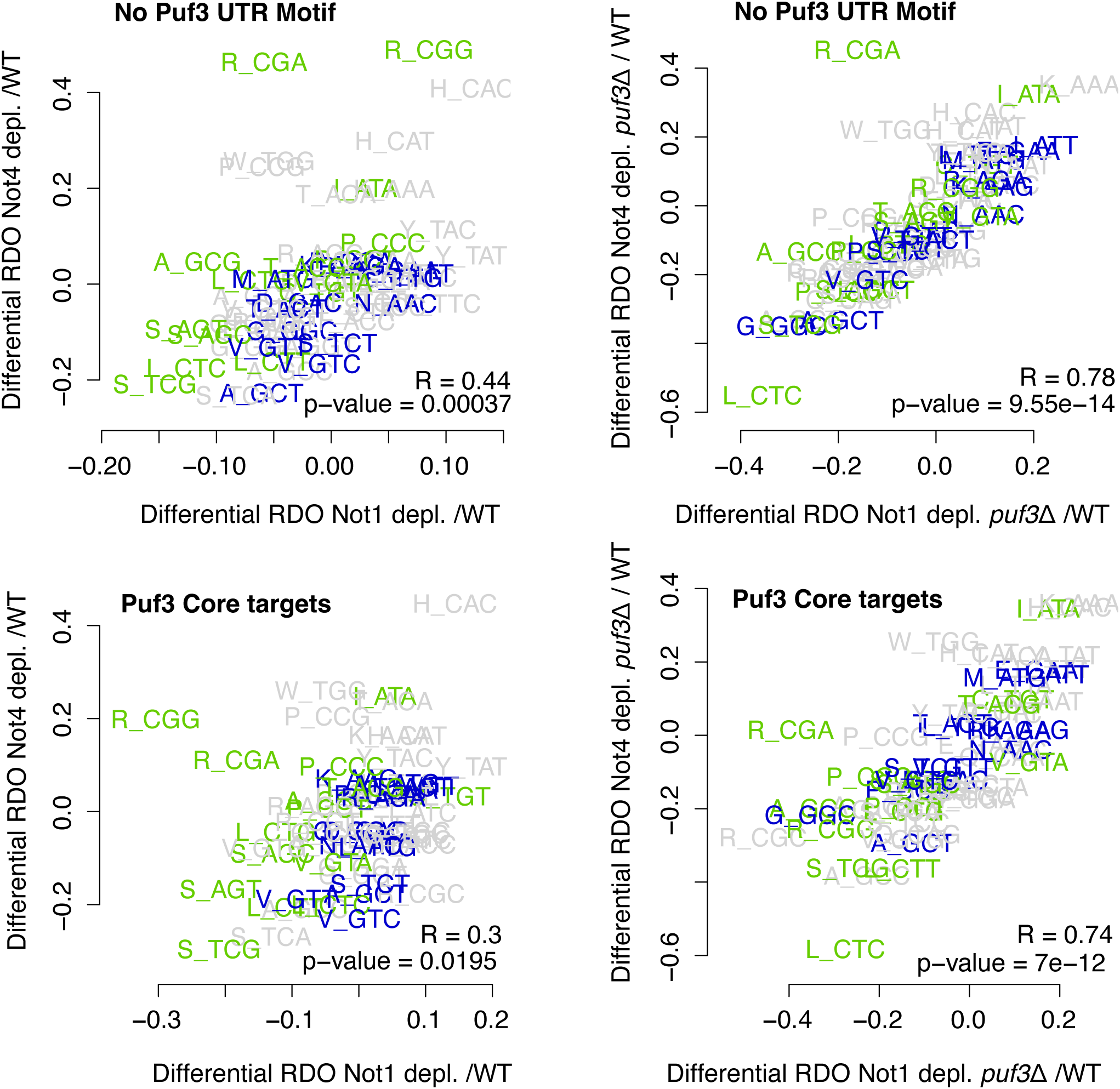
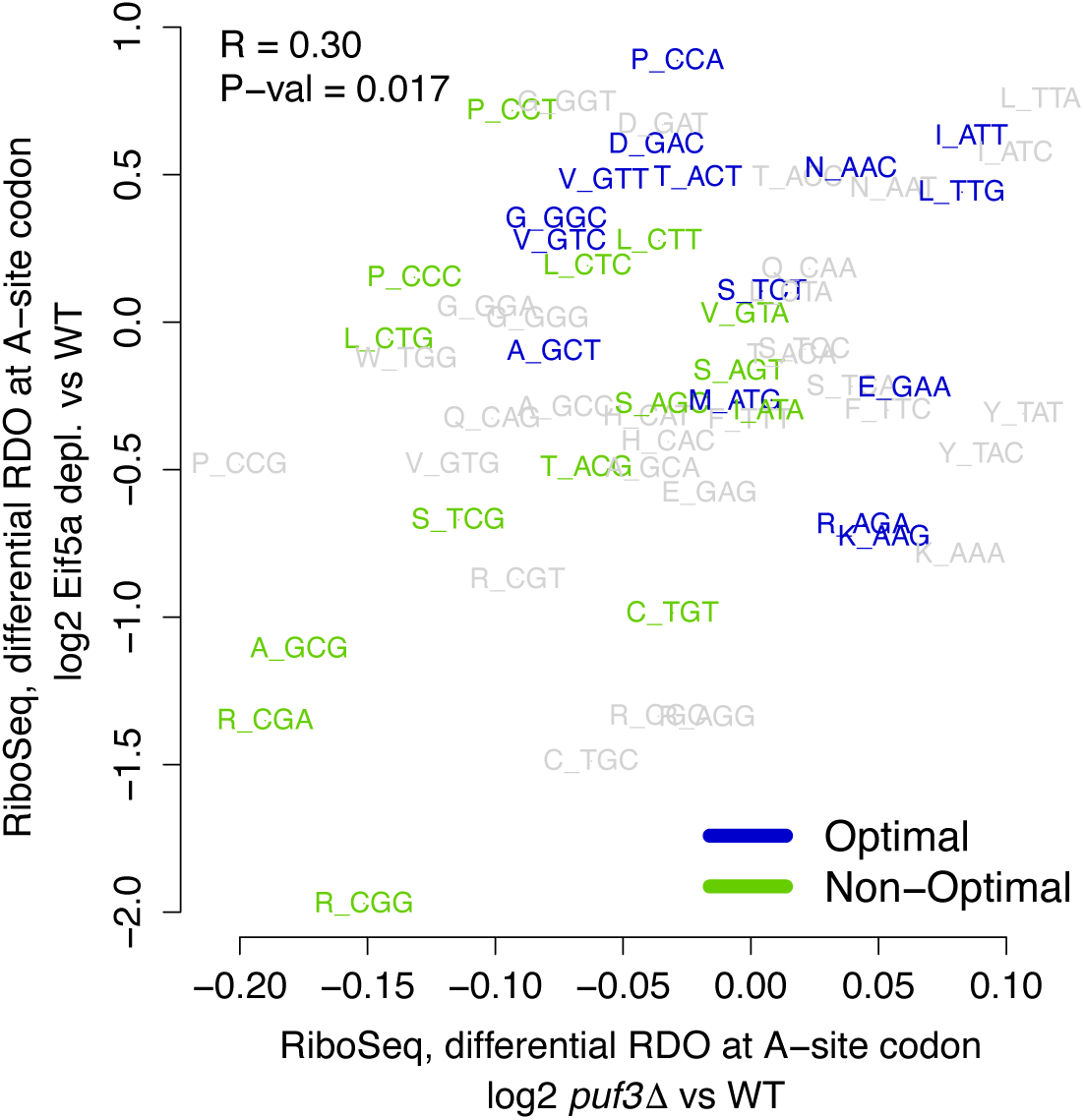
Features of Ribo-Seq upon Not1 and Not4 depletion in presence or absence of Puf3. Related to Figure 3. **A.** Histograms comparing p-site depth one codon upstream of stop for all transcripts with any coverage at this codon in two replicates of at least one sample. The depth is normalized to CDS mean and a log_2_ fold change is taken for Not4 depletion / WT (left), Not4 depletion and *puf3Δ* / WT (middle) and Not4 depletion */* Not4 depletion and *puf3Δ* (right). A histogram of these values is plotted across all included stops. **B.** Scatter plot comparing changes in mRNA solubility and protein aggregation in *not4Δ* versus *not4Δpuf3Δ*. Solubility is a log_2_ fold change of soluble /total expression for each strain, so the values are subtracted between strains. mRNAs that show increased pausing at the stop codon in *not4Δ* versus wild type are indicated in blue. **C.** Scatterplot analyses comparing codon-specific changes in A-site ribosome dwelling occupancies, upon Not1 and Not4 depletion; in cells expressing or lacking Puf3 for Puf3 target mRNAs or mRNAs that are not targets of Puf3 (no Puf3 UTR Motif) as indicated. **D.** Scatterplot comparing codon-specific changes in A-site ribosome dwelling occupancies, upon eIF5A depletion (60) (y-axis) and in *puf3Δ* (x-axis).

**Supplementary Figure S4.**
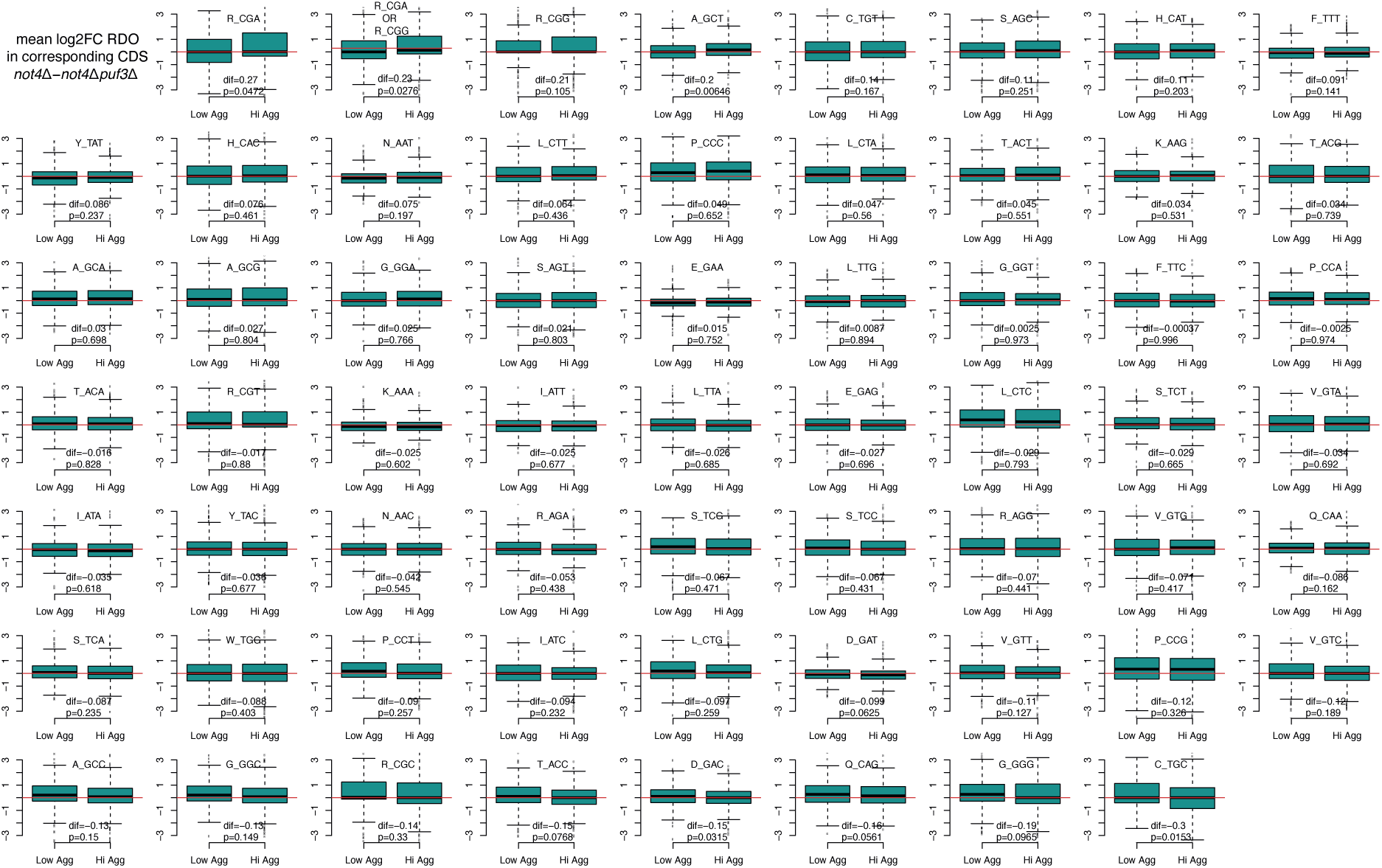
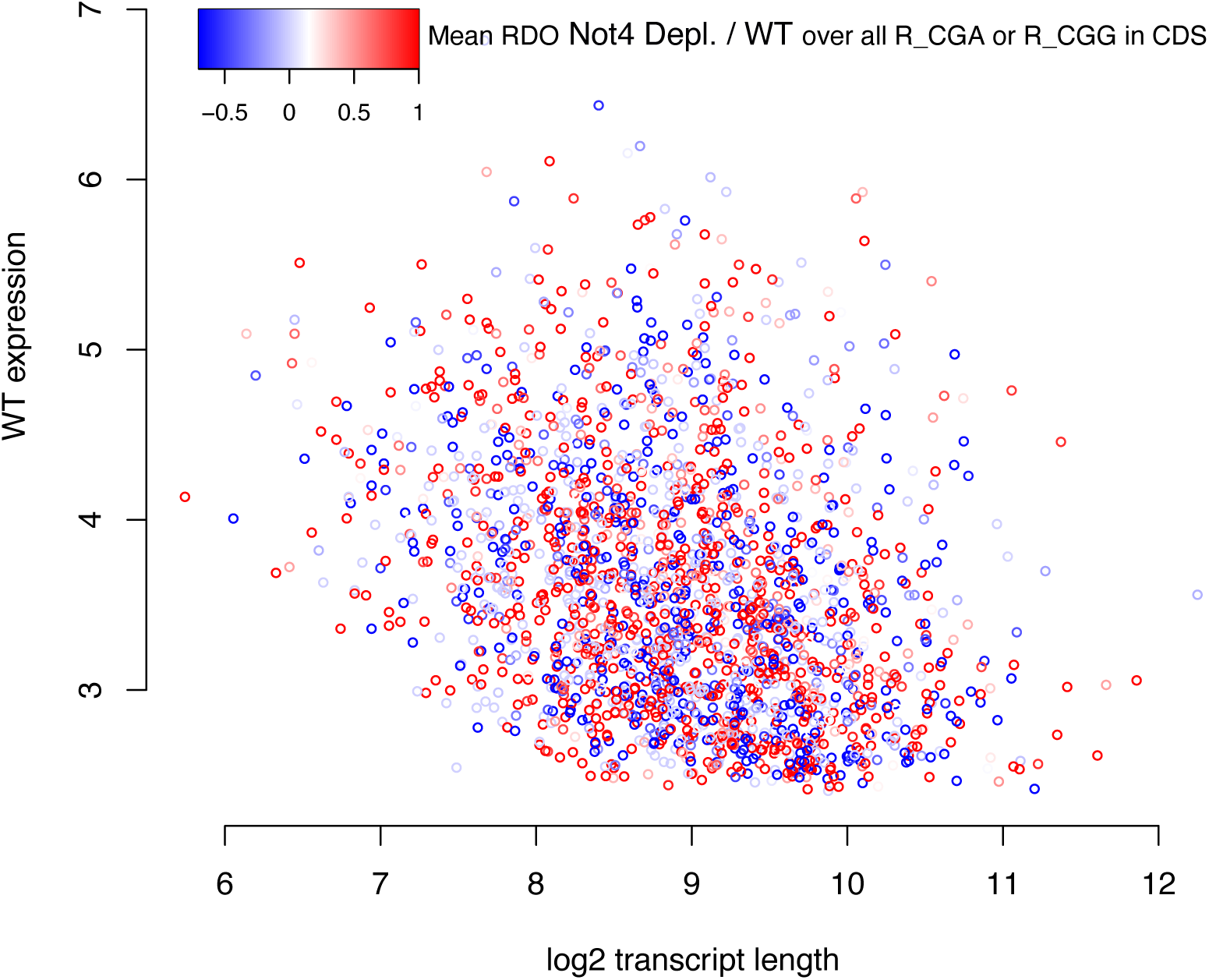
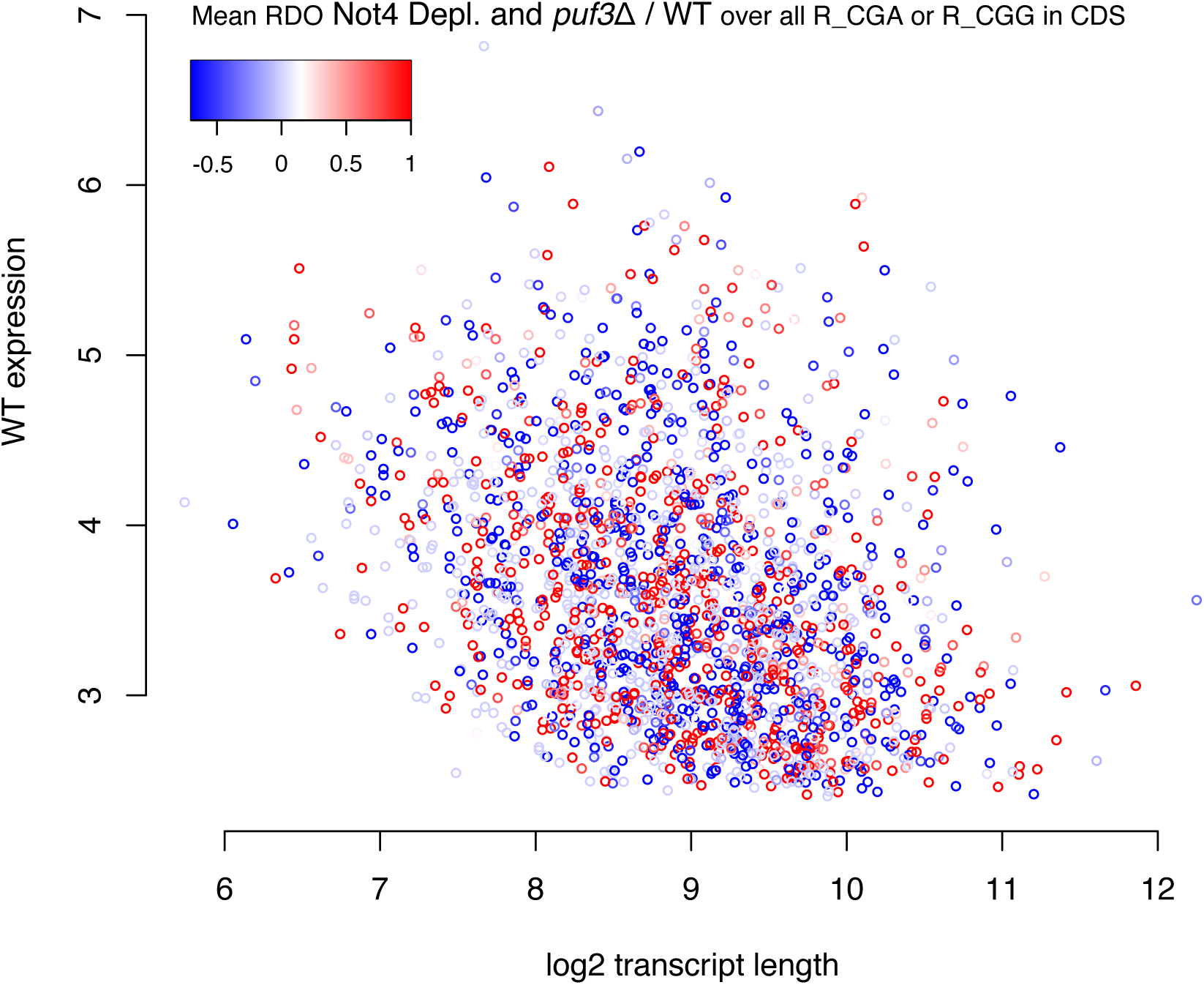
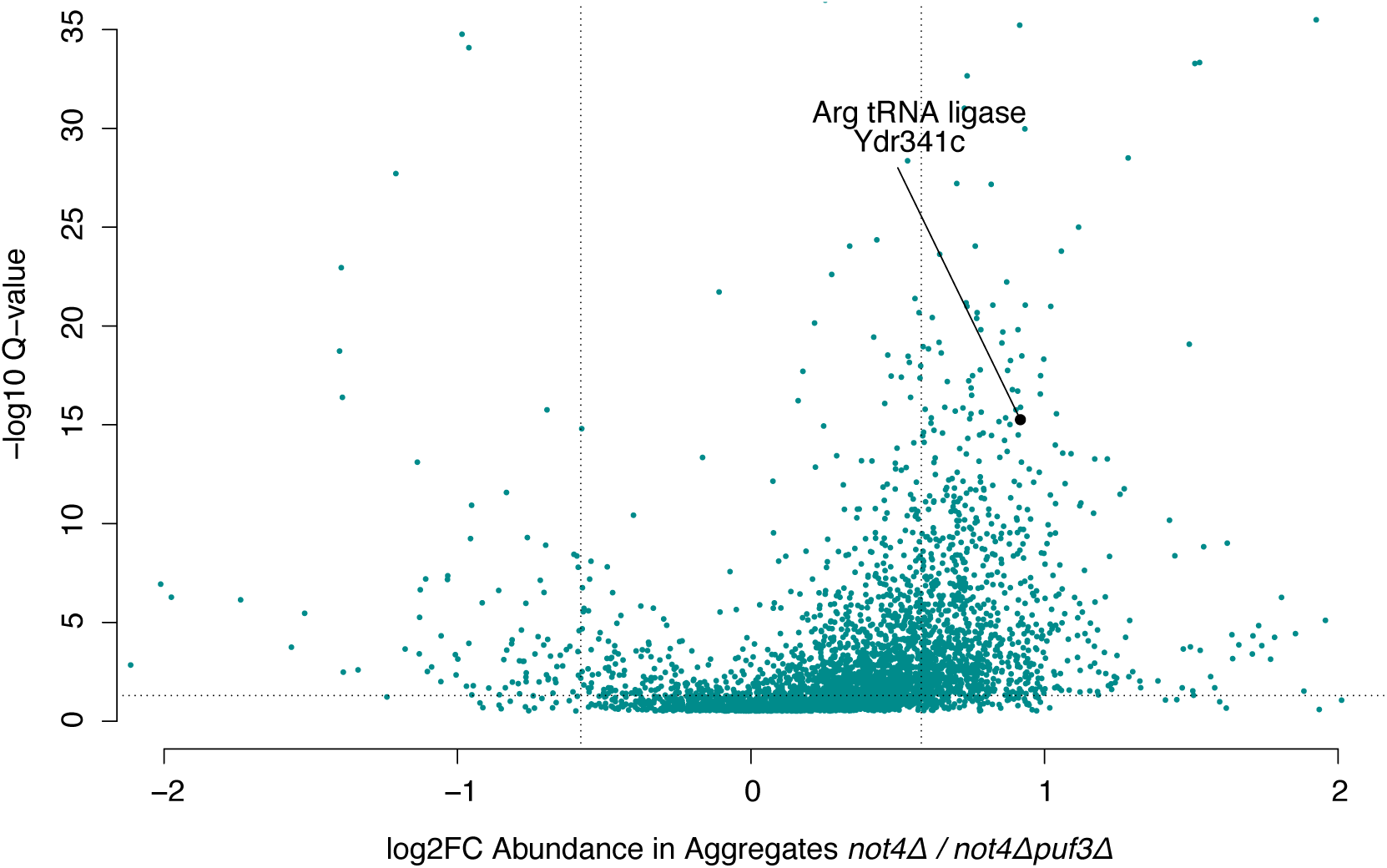
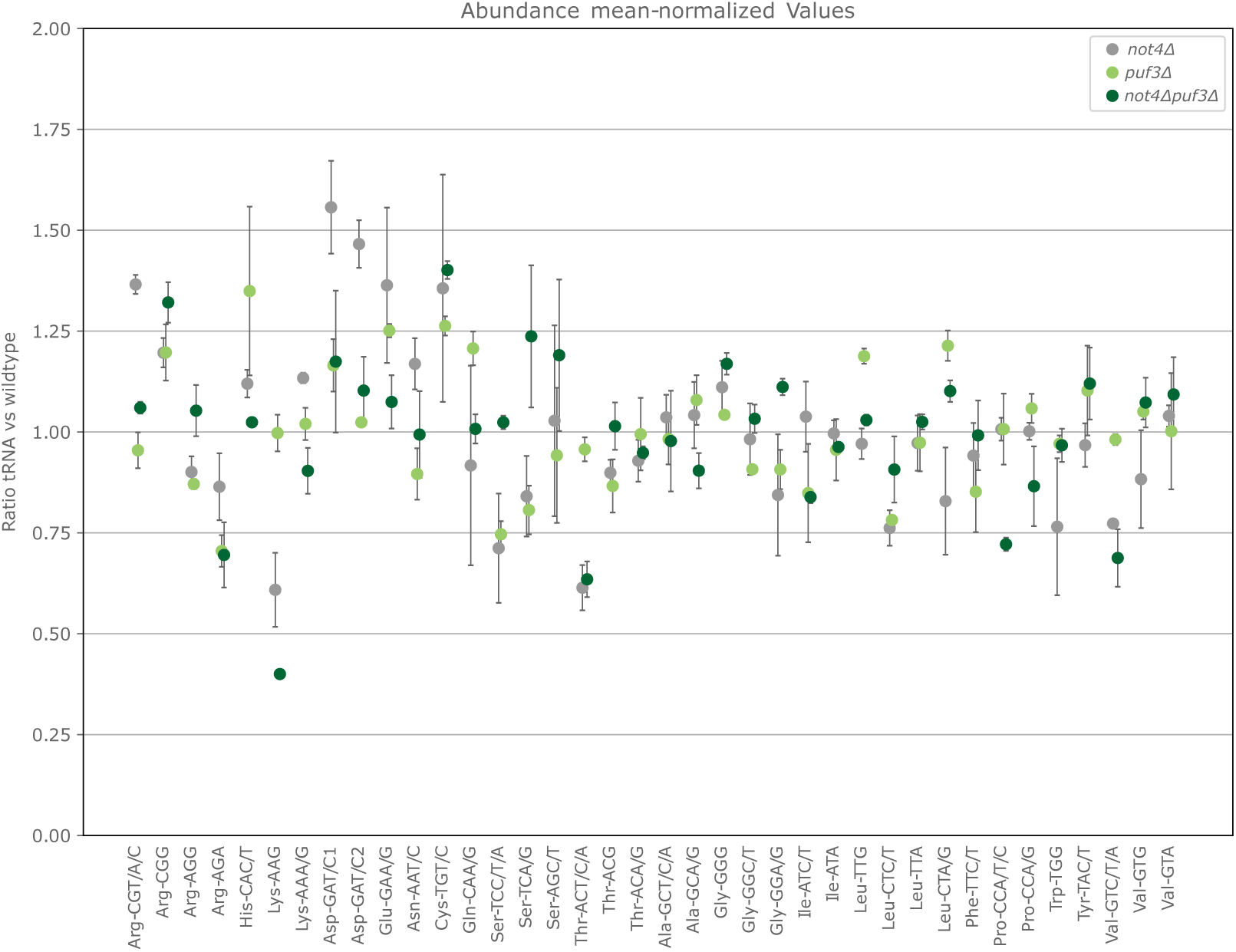
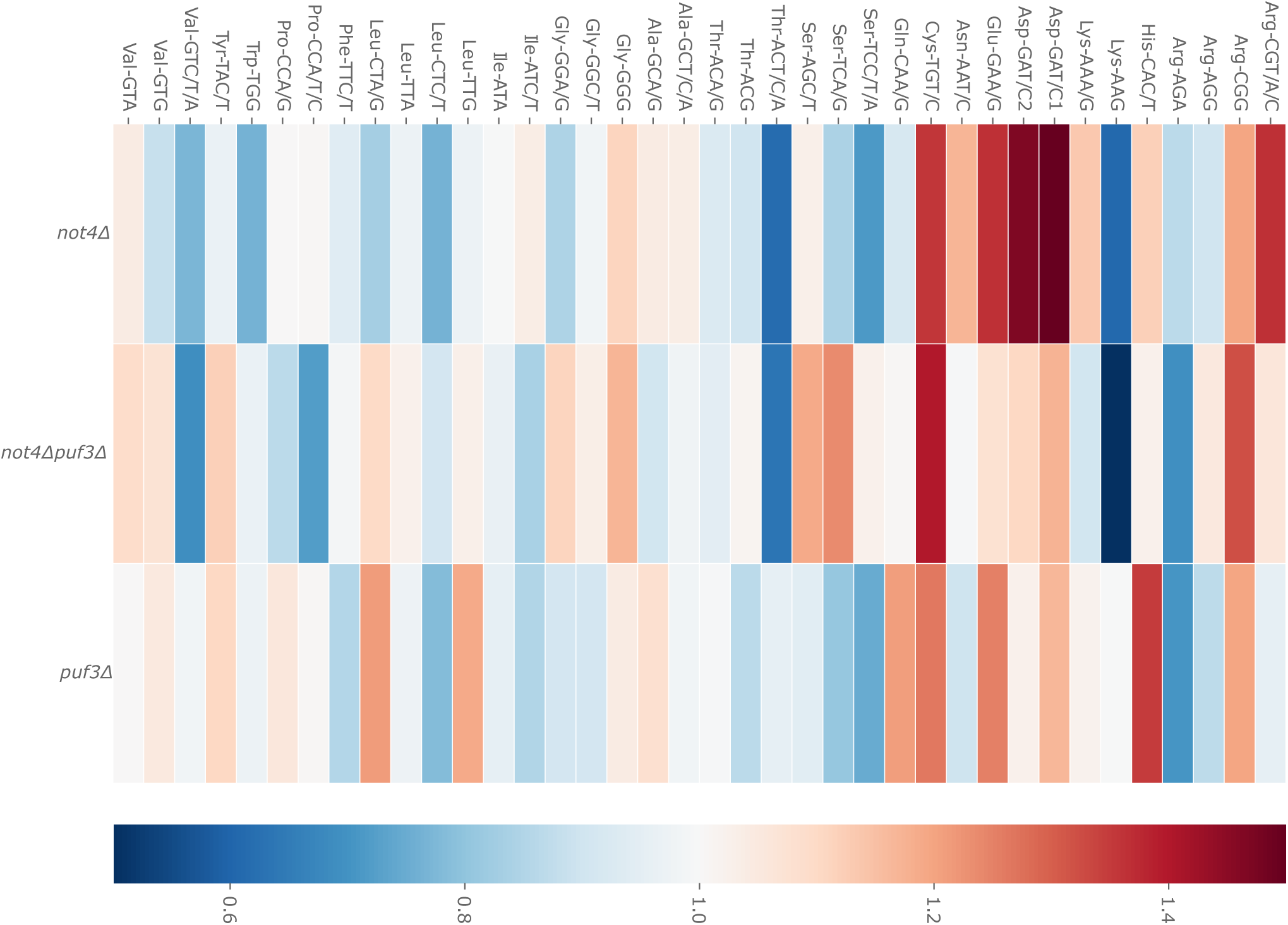
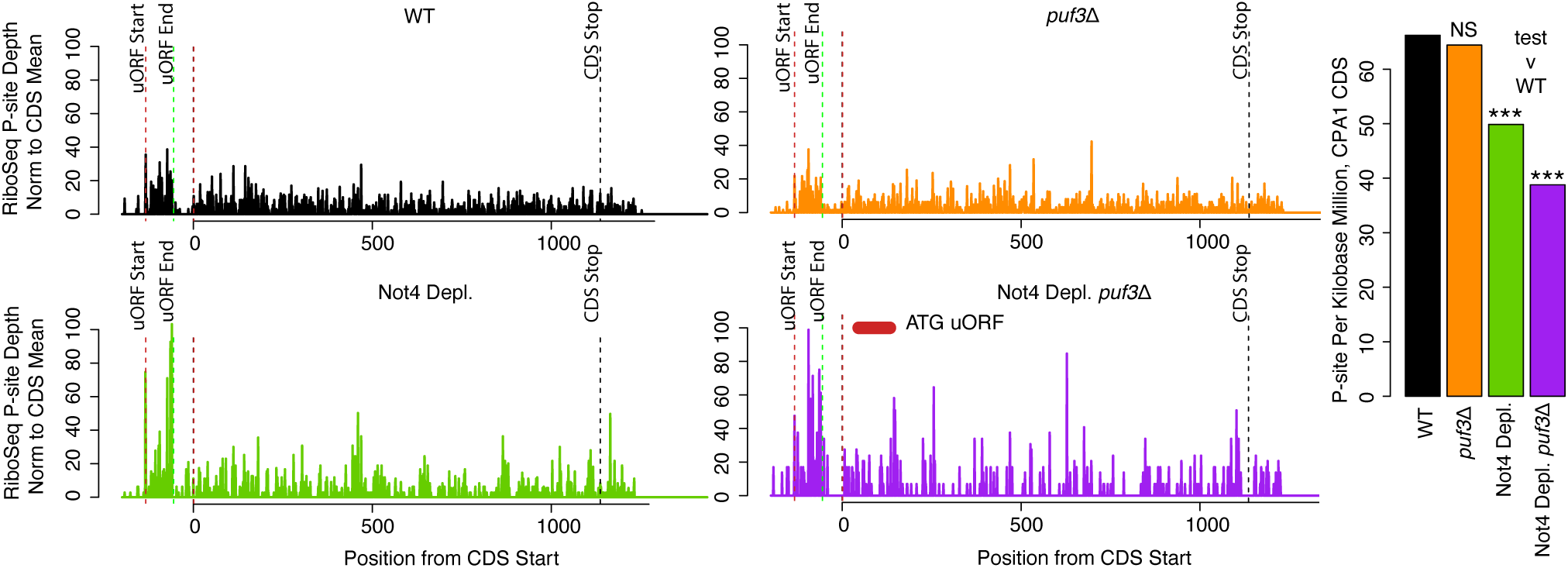
**For mRNAs showing loss of aggregation of encoded protein compared to those that do not, dwelling at rare arginine codons when comparing *not4Δ* to *not4Δpuf3Δ* is the most reduced**. Related to Figure 4. A. Changes in codon-specific A-site RDOs for the 300 mRNAs whose encoded proteins aggregate the least in the *not4Δ* single mutant compared to the double *not4Δpuf3Δ* (designated as “Low Agg”) compared to the 300 mRNAs whose encoded proteins show the highest aggregation in the single compared to the double mutant (“Hi Agg”) were evaluated. The results are shown by box plots for each codon, of the mean RDO change, *not4Δ- not4Δpuf3Δ*, in each CDS of the two groups described above – Low Agg on the left, Hi Agg on the right. These are ordered from the greatest difference (top left) to the lowest (bottom right). B and C. Scatterplots of all mRNAs according to the length (x-axis) and their mean expression level in wild-type strain assessed from the Ribo-Seq (y-axis). mRNAs were color coded if A-site RDOs at rare arginine codons were higher (red) or lower (blue) in the mutant compared to wild type. B. the mutant is Not4 depletion and C. the mutant is Not4 depletion and *puf3Δ*. D. The arginyl-tRNA synthetase is labelled in the volcano plot from Figure 2C depicting aggregated proteins differentially detected in *not4Δ* and *puf3Δnot4Δ.* E. tRNA levels measured in *puf3Δ* and *puf3Δnot4Δ*, compared to wild type, by tRNA-microarrays and presented as relative expression to wild type. For comparison, the data in *not4Δ* compared to wild type measured earlier are included (19). F. tRNA charging using tRNA microarrays measured in *puf3Δ* and *puf3Δnot4Δ* compared to wild type. The measurements in wild type and *not4Δ* as determined earlier (19) are included for comparison. The color key indicates the percentage of charged tRNAs over the total amount of each isoacceptor. The microarrays were performed in duplicates and represented as mean values. Confidence intervals between both replicates is 97% for *puf3Δ,* 99% for *puf3Δnot4Δ,* 95% for *not4Δ*, and 98% for wild-type, respectively. In E-F, tRNA probes are depicted with their cognate codon and the corresponding amino acid. G. Ribo-Seq P-site depths at the *CPA1* locus, both upstream ORF and CDS sequences, in wild type, upon Not4 depletion, in *puf3Δ* and upon Not4 depletion and *puf3Δ* (left 4 panels) and barplot of P-sites per kilobase million on the CDS for all these strains with significance test of expression change versus WT for all non-WT strains. The P-site depths at each position on *CPA1* mRNA in the single and double Not4 depletion mutant were normalized to the CDS mean and both showed high values in the uORF, relative to the main CDS.

**Supplementary Figure S5.**
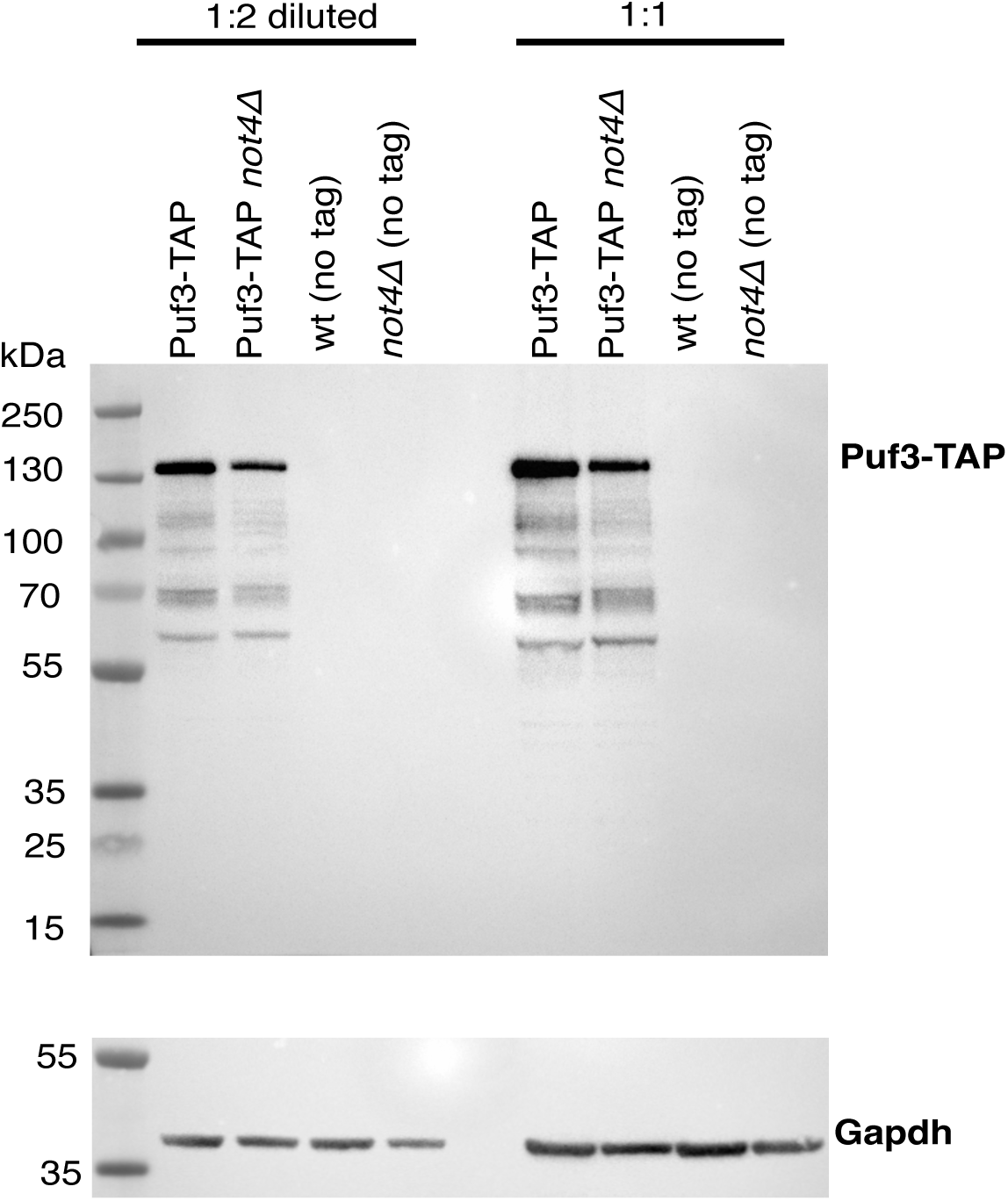
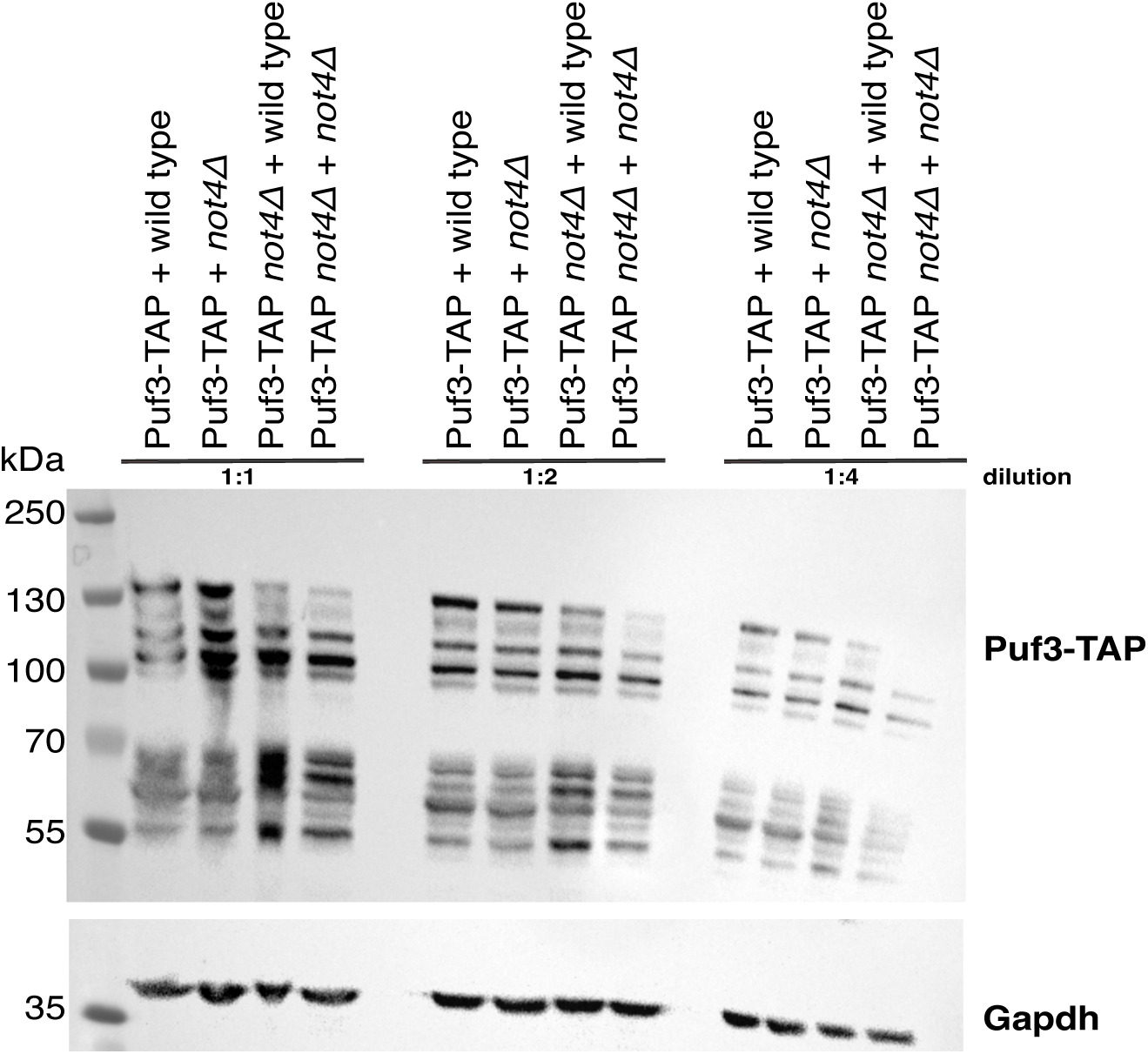
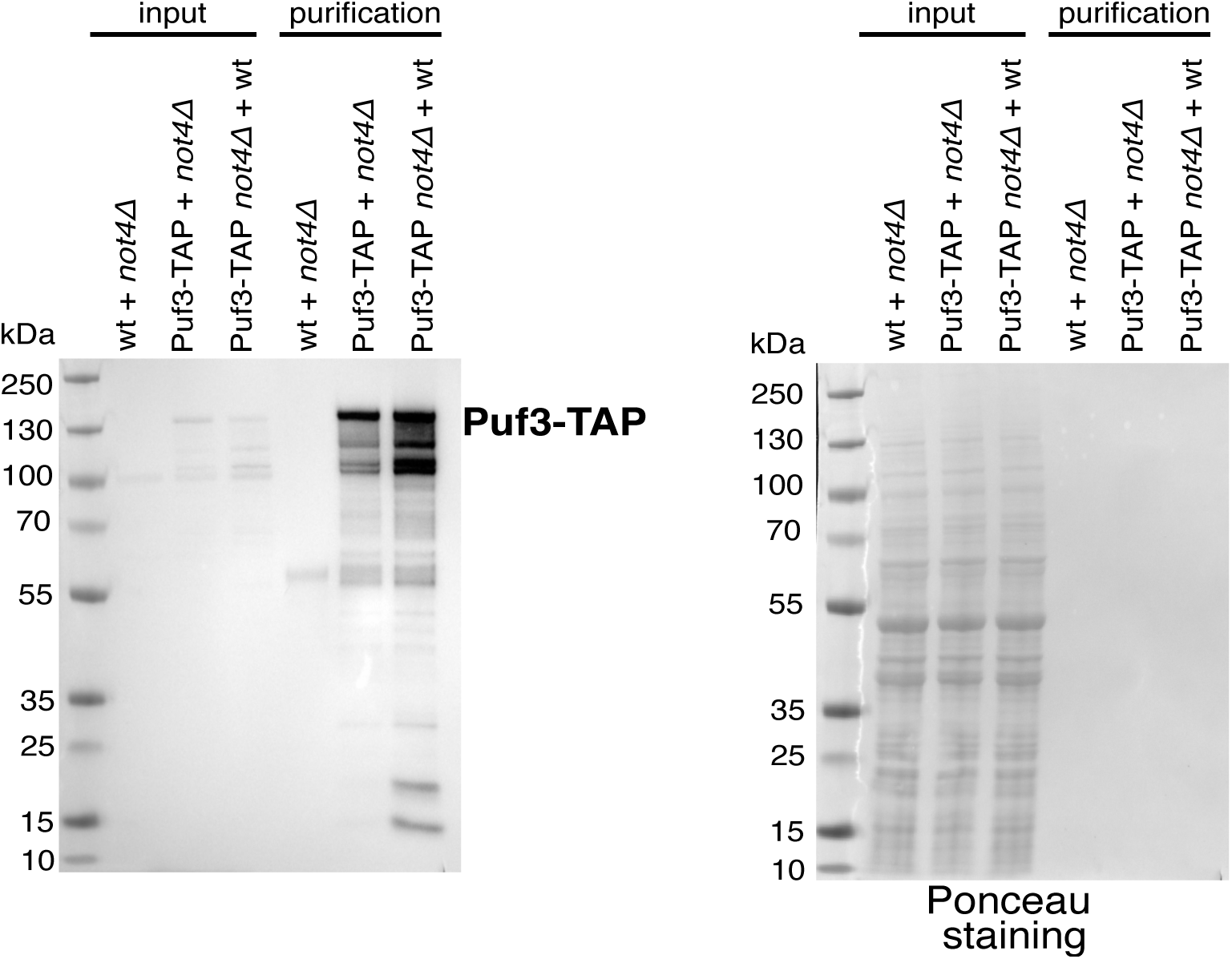
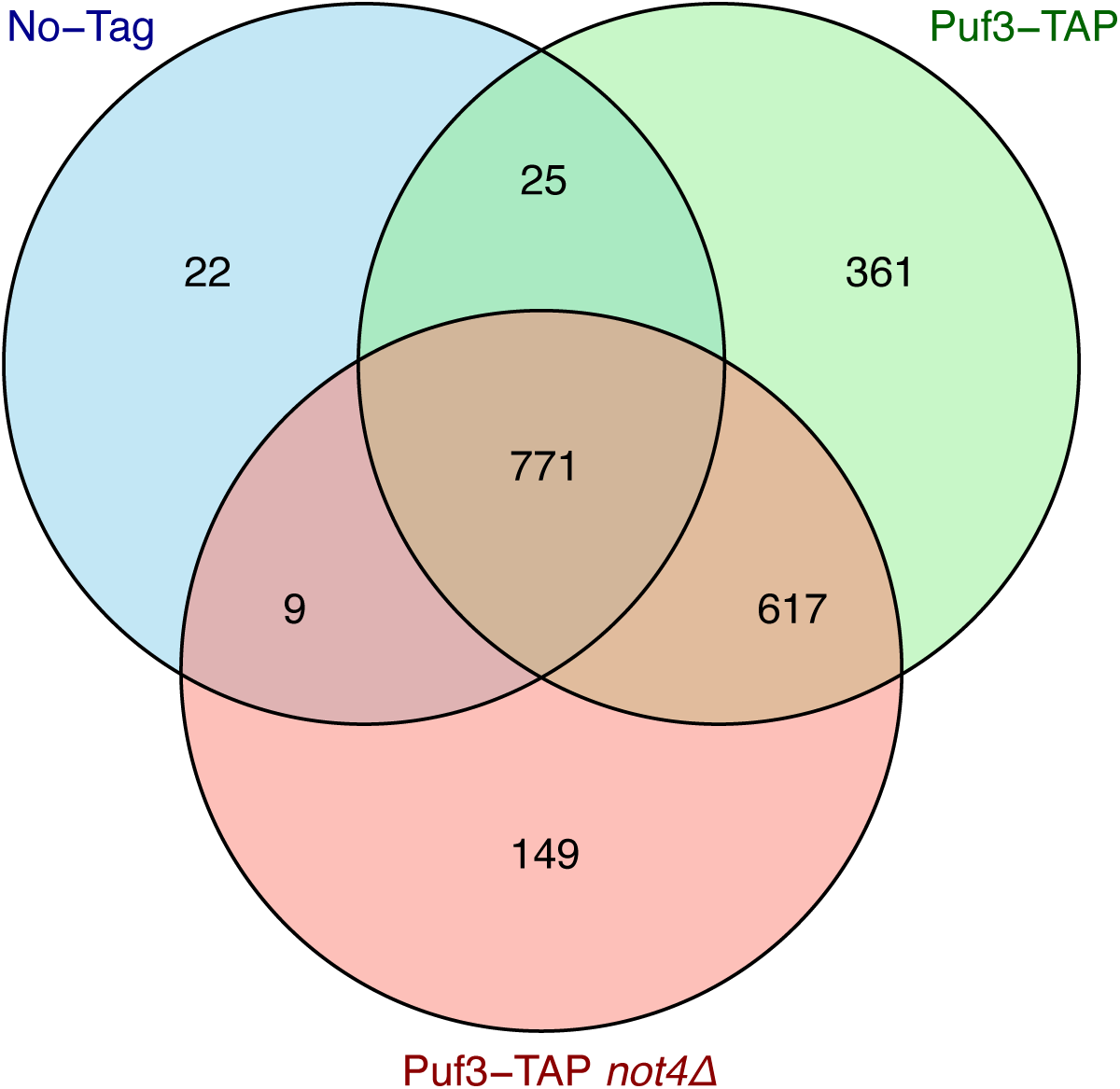
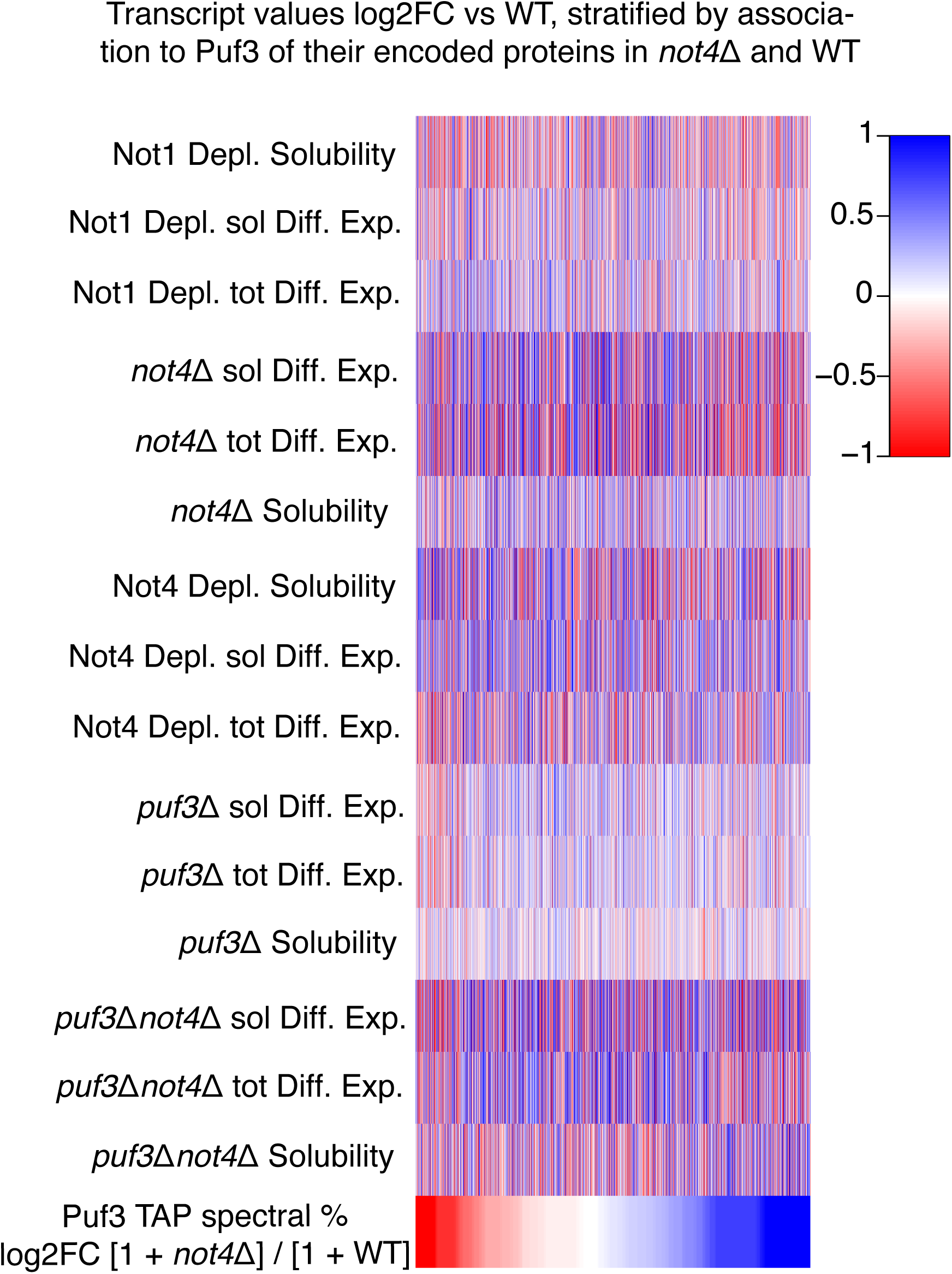
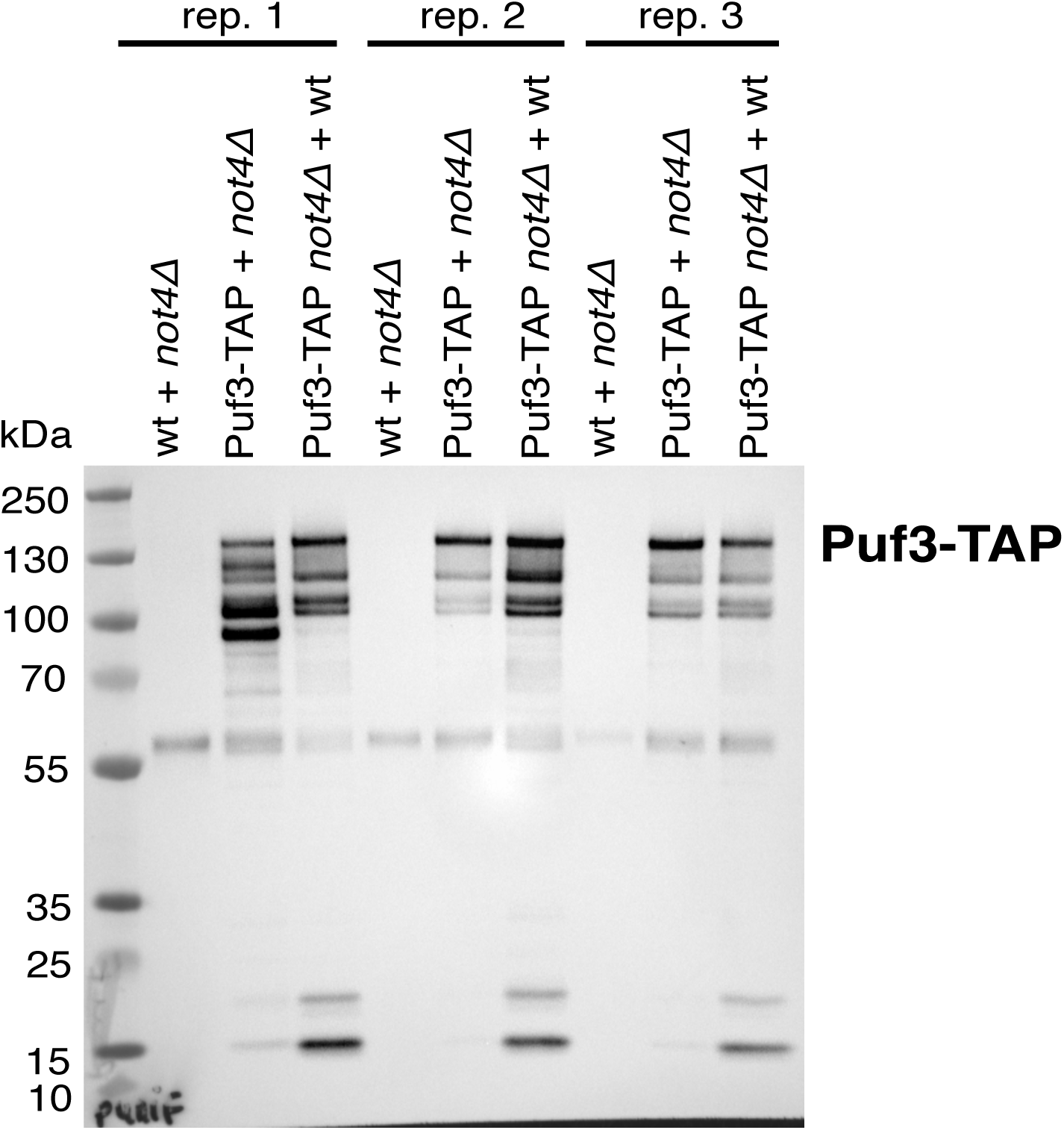
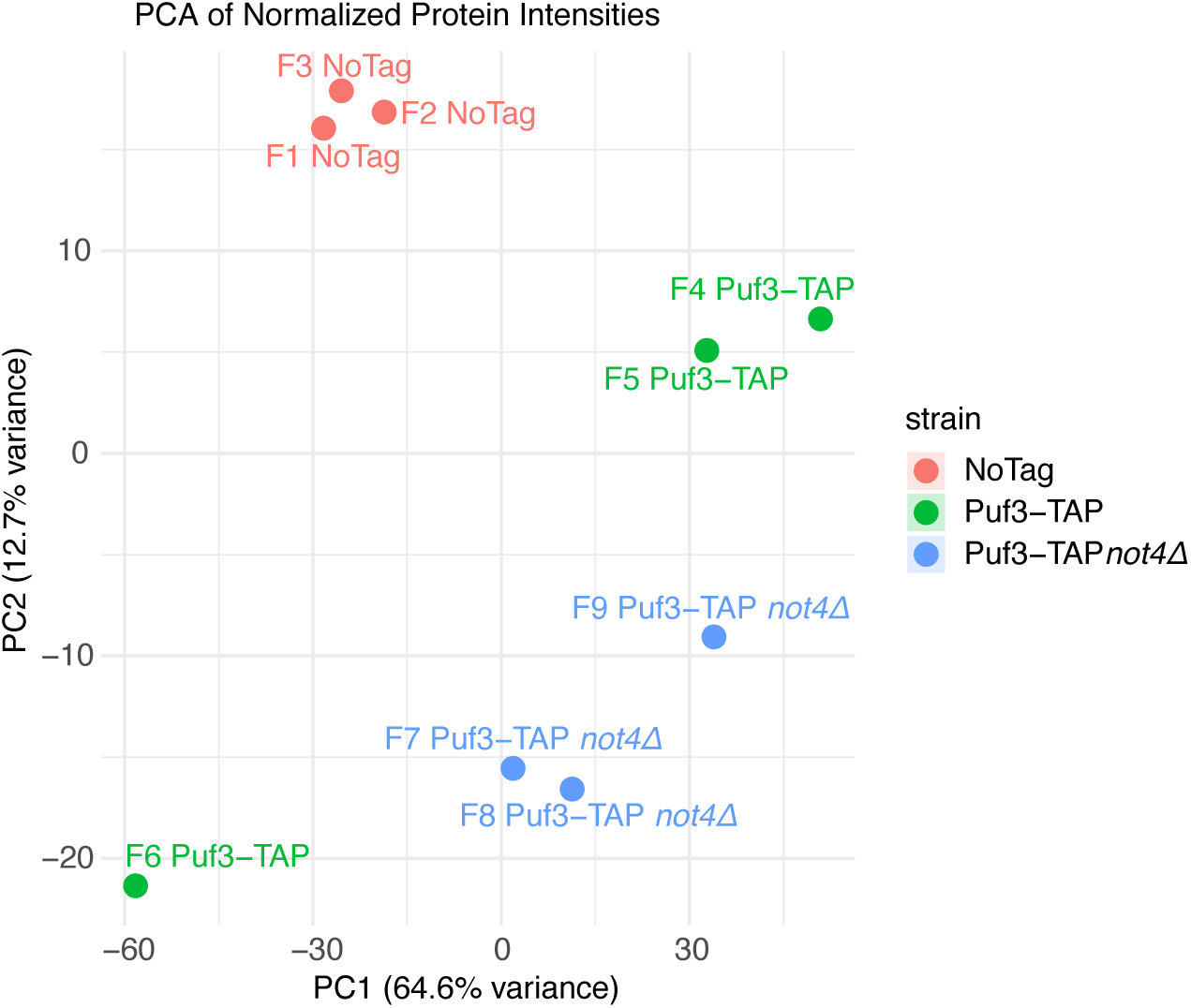
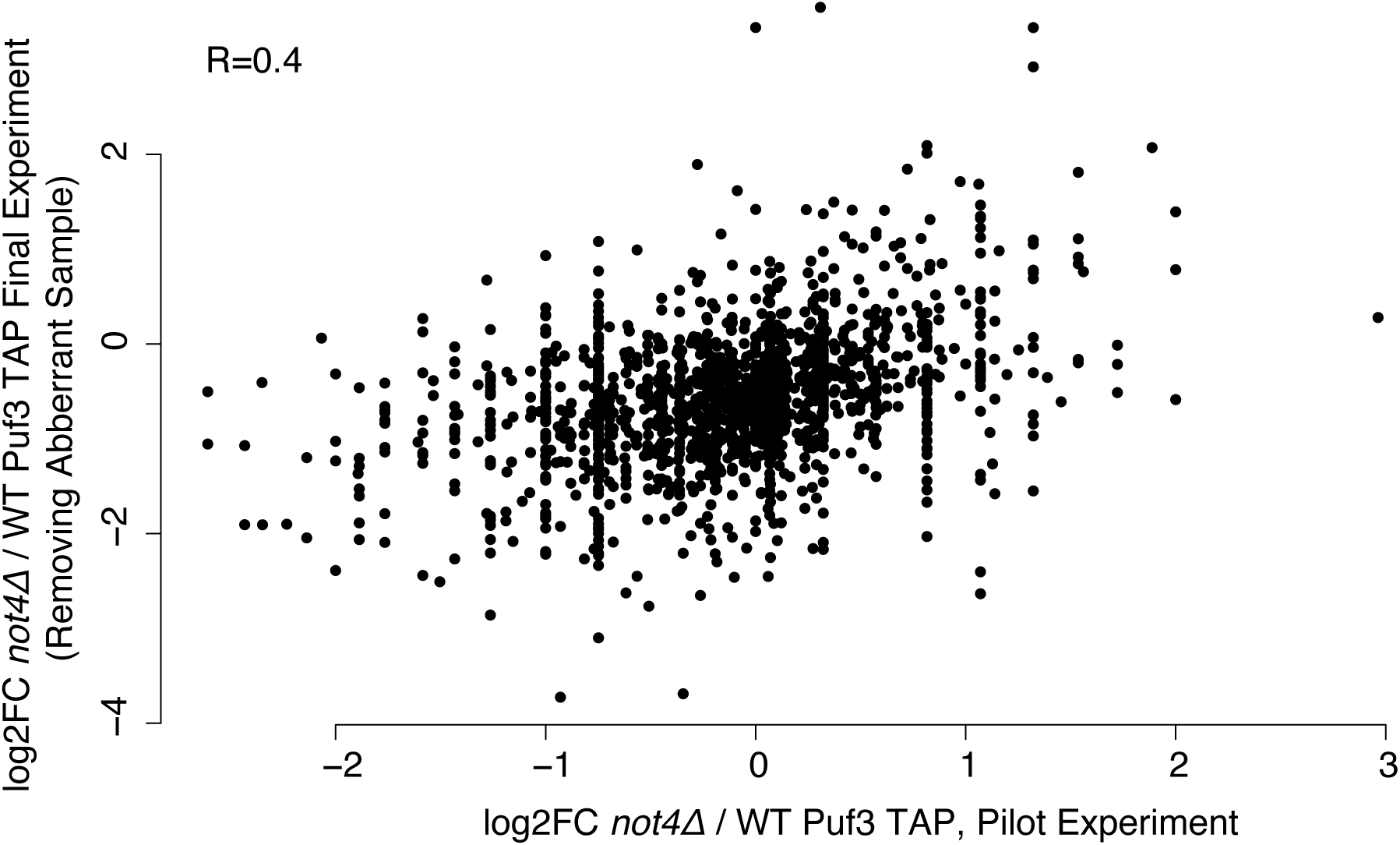
**Differential interactomes of Puf3 are identified in wild type and *not4Δ*. Related to Figure 5**. A. Total protein from post-alkaline lysis from wild type and *not4Δ* cells expressing Tap-tagged Puf3 were analyzed by western blotting with anti-PAP antibodies. Antibodies against Gapdh were used as loading control. **B**. Puf3 integrity test. Cell pellets were mixed together as described on the figure legend prior to protein extraction. The samples were analysed by western blotting using anti-PAP and Gapdh antibodies. **C.** Western blot analysis with anti-PAP antibodies to follow the pilot purification of Puf3-TAP for mass spectrometry analysis. **D.** Venn diagram for the pilot experiment indicating proteins identified in the control untagged strain purification, and purifications from the wild type and *not4Δ* strains expressing Tap-tagged Puf3. **E.** Changes in expression and solubility of mRNAs in the different indicated strains is compared to changes in differential identification of encoded proteins in the pilot affinity purification of Puf3 in wild type and *not4Δ*. **F**. Western blot analysis with anti-PAP antibodies to follow the triplicate purifications of Puf3-TAP for mass spectrometry analysis. **G.** Principal component analysis, based on normalized protein levels. **H.** Scatter plot comparing the triplicates purification of Puf3-TAP without the outlier sample to the pilot purification of Puf3-TAP.

## REFERENCES

1. Collart, M.A. (2016) The Ccr4-Not complex is a key regulator of eukaryotic gene expression. Wiley Interdiscip Rev RNA, 7, 438–454.

2. Maillet, L., Tu, C., Hong, Y.K., Shuster, E.O. and Collart, M.A. (2000) The essential function of Not1 lies within the Ccr4-Not complex. J Mol Biol, 303, 131–143.

3. Bhaskar, V., Basquin, J. and Conti, E. (2015) Architecture of the ubiquitylation module of the yeast Ccr4-Not complex. Structure, 23, 921–928.

4. Bhaskar, V., Roudko, V., Basquin, J., Sharma, K., Urlaub, H., Seraphin, B. and Conti, E. (2013) Structure and RNA-binding properties of the Not1-Not2-Not5 module of the yeast Ccr4-Not complex. Nat Struct Mol Biol, 20, 1281–1288.

5. Basquin, J., Roudko, V.V., Rode, M., Basquin, C., Seraphin, B. and Conti, E. (2012) Architecture of the nuclease module of the yeast Ccr4-not complex: the Not1-Caf1-Ccr4 interaction. Mol Cell, 48, 207–218.

6. Boland, A., Chen, Y., Raisch, T., Jonas, S., Kuzuoglu-Ozturk, D., Wohlbold, L., Weichenrieder, O. and Izaurralde, E. (2013) Structure and assembly of the NOT module of the human CCR4-NOT complex. Nat Struct Mol Biol, 20, 1289–1297.

7. Sgromo, A., Raisch, T., Bawankar, P., Bhandari, D., Chen, Y., Kuzuoglu-Ozturk, D., Weichenrieder, O. and Izaurralde, E. (2017) A CAF40-binding motif facilitates recruitment of the CCR4-NOT complex to mRNAs targeted by Drosophila Roquin. Nat Commun, 8, 14307.

8. Bartlam, M. and Yamamoto, T. (2010) The structural basis for deadenylation by the CCR4-NOT complex. Protein Cell, 1, 443–452.

9. Raisch, T. and Valkov, E. (2022) Regulation of the multisubunit CCR4-NOT deadenylase in the initiation of mRNA degradation. Curr Opin Struct Biol, 77, 102460.

10. Pavanello, L., Hall, M. and Winkler, G.S. (2023) Regulation of eukaryotic mRNA deadenylation and degradation by the Ccr4-Not complex. Front Cell Dev Biol, 11, 1153624.

11. Stowell, J.A.W., Webster, M.W., Kogel, A., Wolf, J., Shelley, K.L. and Passmore, L.A. (2016) Reconstitution of Targeted Deadenylation by the Ccr4-Not Complex and the YTH Domain Protein Mmi1. Cell Rep, 17, 1978–1989.

12. Guenole, A., Velilla, F., Chartier, A., Rich, A., Carvunis, A.R., Sardet, C., Simonelig, M. and Sobhian, B. (2022) RNF219 regulates CCR4-NOT function in mRNA translation and deadenylation. Sci Rep, 12, 9288.

13. Wilczynska, A., Gillen, S.L., Schmidt, T., Meijer, H.A., Jukes-Jones, R., Langlais, C., Kopra, K., Lu, W.T., Godfrey, J.D., Hawley, B.R. et al. (2019) eIF4A2 drives repression of translation at initiation by Ccr4-Not through purine-rich motifs in the 5’UTR. Genome Biol, 20, 262.

14. Kuzuoglu-Ozturk, D., Bhandari, D., Huntzinger, E., Fauser, M., Helms, S. and Izaurralde, E. (2016) miRISC and the CCR4-NOT complex silence mRNA targets independently of 43S ribosomal scanning. EMBO J, 35, 1186–1203.

15. Okada, H., Schittenhelm, R.B., Straessle, A. and Hafen, E. (2015) Multi-functional regulation of 4E-BP gene expression by the Ccr4-Not complex. PLoS One, 10, e0113902.

16. Rasch, F., Weber, R., Izaurralde, E. and Igreja, C. (2020) 4E-T-bound mRNAs are stored in a silenced and deadenylated form. Genes Dev, 34, 847–860.

17. Allen, G., Weiss, B., Panasenko, O.O., Huch, S., Villanyi, Z., Albert, B., Dilg, D., Zagatti, M., Schaughency, P., Liao, S.E. et al. (2023) Not1 and Not4 inversely determine mRNA solubility that sets the dynamics of co-translational events. Genome Biol, 24, 30.

18. Gupta, I., Villanyi, Z., Kassem, S., Hughes, C., Panasenko, O.O., Steinmetz, L.M. and Collart, M.A. (2016) Translational Capacity of a Cell Is Determined during Transcription Elongation via the Ccr4-Not Complex. Cell Rep, 15, 1782–1794.

19. Allen, G.E., Panasenko, O.O., Villanyi, Z., Zagatti, M., Weiss, B., Pagliazzo, L., Huch, S., Polte, C., Zahoran, S., Hughes, C.S. et al. (2021) Not4 and Not5 modulate translation elongation by Rps7A ubiquitination, Rli1 moonlighting, and condensates that exclude eIF5A. Cell Rep, 36, 109633.

20. Chen, S., Allen, G., Panasenko, O.O. and Collart, M.A. (2023) Not4-dependent targeting of MMF1 mRNA to mitochondria limits its expression via ribosome pausing, Egd1 ubiquitination, Caf130, no-go-decay and autophagy. Nucleic Acids Res, 51, 5022–5039.

21. Halter, D., Collart, M.A. and Panasenko, O.O. (2014) The Not4 E3 ligase and CCR4 deadenylase play distinct roles in protein quality control. PLoS One, 9, e86218.

22. Panasenko, O.O. and Collart, M.A. (2012) Presence of Not5 and ubiquitinated Rps7A in polysome fractions depends upon the Not4 E3 ligase. Mol Microbiol, 83, 640–653.

23. Panasenko, O.O. and Collart, M.A. (2011) Not4 E3 ligase contributes to proteasome assembly and functional integrity in part through Ecm29. Mol Cell Biol, 31, 1610–1623.

24. Kassem, S., Villanyi, Z. and Collart, M.A. (2017) Not5-dependent co-translational assembly of Ada2 and Spt20 is essential for functional integrity of SAGA. Nucleic Acids Res, 45, 1186–1199.

25. Pekovic, F., Rammelt, C., Kubikova, J., Metz, J., Jeske, M. and Wahle, E. (2023) RNA binding proteins Smaug and Cup induce CCR4-NOT-dependent deadenylation of the nanos mRNA in a reconstituted system. Nucleic Acids Res, 51, 3950–3970.

26. Vindry, C., Lauwers, A., Hutin, D., Soin, R., Wauquier, C., Kruys, V. and Gueydan, C. (2012) dTIS11 Protein-dependent polysomal deadenylation is the key step in AU-rich element-mediated mRNA decay in Drosophila cells. J Biol Chem, 287, 35527–35538.

27. Fabian, M.R., Frank, F., Rouya, C., Siddiqui, N., Lai, W.S., Karetnikov, A., Blackshear, P.J., Nagar, B. and Sonenberg, N. (2013) Structural basis for the recruitment of the human CCR4-NOT deadenylase complex by tristetraprolin. Nat Struct Mol Biol, 20, 735–739.

28. Chekulaeva, M., Mathys, H., Zipprich, J.T., Attig, J., Colic, M., Parker, R. and Filipowicz, W. (2011) miRNA repression involves GW182-mediated recruitment of CCR4-NOT through conserved W-containing motifs. Nat Struct Mol Biol, 18, 1218–1226.

29. Buschauer, R., Matsuo, Y., Sugiyama, T., Chen, Y.H., Alhusaini, N., Sweet, T., Ikeuchi, K., Cheng, J., Matsuki, Y., Nobuta, R. et al. (2020) The Ccr4-Not complex monitors the translating ribosome for codon optimality. Science, 368.

30. Panasenko, O.O., Somasekharan, S.P., Villanyi, Z., Zagatti, M., Bezrukov, F., Rashpa, R., Cornut, J., Iqbal, J., Longis, M., Carl, S.H. et al. (2019) Co-translational assembly of proteasome subunits in NOT1-containing assemblysomes. Nat Struct Mol Biol, 26, 110–120.

31. Olivas, W. and Parker, R. (2000) The Puf3 protein is a transcript-specific regulator of mRNA degradation in yeast. EMBO J, 19, 6602–6611.

32. Webster, M.W., Stowell, J.A. and Passmore, L.A. (2019) RNA-binding proteins distinguish between similar sequence motifs to promote targeted deadenylation by Ccr4-Not. Elife, 8.

33. Kershaw, C.J., Costello, J.L., Talavera, D., Rowe, W., Castelli, L.M., Sims, P.F., Grant, C.M., Ashe, M.P., Hubbard, S.J. and Pavitt, G.D. (2015) Integrated multi-omics analyses reveal the pleiotropic nature of the control of gene expression by Puf3p. Sci Rep, 5, 15518.

34. Gerber, A.P., Herschlag, D. and Brown, P.O. (2004) Extensive association of functionally and cytotopically related mRNAs with Puf family RNA-binding proteins in yeast. PLoS Biol, 2, E79.

35. Lapointe, C.P., Stefely, J.A., Jochem, A., Hutchins, P.D., Wilson, G.M., Kwiecien, N.W., Coon, J.J., Wickens, M. and Pagliarini, D.J. (2018) Multi-omics Reveal Specific Targets of the RNA-Binding Protein Puf3p and Its Orchestration of Mitochondrial Biogenesis. Cell Syst, 6, 125–135 e126.

36. Rowe, W., Kershaw, C.J., Castelli, L.M., Costello, J.L., Ashe, M.P., Grant, C.M., Sims, P.F., Pavitt, G.D. and Hubbard, S.J. (2014) Puf3p induces translational repression of genes linked to oxidative stress. Nucleic Acids Res, 42, 1026–1041.

37. Liu, S., Liu, S., He, B., Li, L., Li, L., Wang, J., Cai, T., Chen, S. and Jiang, H. (2021) OXPHOS deficiency activates global adaptation pathways to maintain mitochondrial membrane potential. EMBO Rep, 22, e51606.

38. Lee, D., Ohn, T., Chiang, Y.C., Quigley, G., Yao, G., Liu, Y. and Denis, C.L. (2010) PUF3 acceleration of deadenylation in vivo can operate independently of CCR4 activity, possibly involving effects on the PAB1-mRNP structure. J Mol Biol, 399, 562–575.

39. Bulmash, A.S., Fischer, A.D., Russo, J., Mueller, S.M. and Olivas, W.M. (2023) Yeast Puf3p-mediated mRNA decay is regulated by carbon source-specific differential interaction of Puf3p with Pop2p and Yak1p. FEBS Lett, 597, 1606–1622.

40. Lee, C.D. and Tu, B.P. (2015) Glucose-Regulated Phosphorylation of the PUF Protein Puf3 Regulates the Translational Fate of Its Bound mRNAs and Association with RNA Granules. Cell Rep, 11, 1638–1650.

41. Miller, M.A., Russo, J., Fischer, A.D., Lopez Leban, F.A. and Olivas, W.M. (2014) Carbon source-dependent alteration of Puf3p activity mediates rapid changes in the stabilities of mRNAs involved in mitochondrial function. Nucleic Acids Res, 42, 3954–3970.

42. Garrido-Godino, A.I., Gupta, I., Gutierrez-Santiago, F., Martinez-Padilla, A.B., Alekseenko, A., Steinmetz, L.M., Perez-Ortin, J.E., Pelechano, V. and Navarro, F. (2021) Rpb4 and Puf3 imprint and post-transcriptionally control the stability of a common set of mRNAs in yeast. RNA Biol, 18, 1206–1220.

43. Kruk, J.A., Dutta, A., Fu, J., Gilmour, D.S. and Reese, J.C. (2011) The multifunctional Ccr4-Not complex directly promotes transcription elongation. Genes Dev, 25, 581–593.

44. Collart, M.A. and Oliviero, S. (2001) Preparation of yeast RNA. Curr Protoc Mol Biol, Chapter 13, Unit13 12.

45. Wang, Z., Sun, X., Wee, J., Guo, X. and Gu, Z. (2019) Novel insights into global translational regulation through Pumilio family RNA-binding protein Puf3p revealed by ribosomal profiling. Curr Genet, 65, 201–212.

46. Martin, M. (2011) Cutadapt Removes Adapter Sequences From High-Throughput Sequencing Reads. Embnet, 17.

47. Kim, D., Langmead, B. and Salzberg, S.L. (2015) HISAT: a fast spliced aligner with low memory requirements. Nat Methods, 12, 357–360.

48. Langmead, B. and Salzberg, S.L. (2012) Fast gapped-read alignment with Bowtie 2. Nat Methods, 9, 357–359.

49. Lauria, F., Tebaldi, T., Bernabo, P., Groen, E.J.N., Gillingwater, T.H. and Viero, G. (2018) riboWaltz: Optimization of ribosome P-site positioning in ribosome profiling data. PLoS Comput Biol, 14, e1006169.

50. Love, M.I., Huber, W. and Anders, S. (2014) Moderated estimation of fold change and dispersion for RNA-seq data with DESeq2. Genome Biol, 15, 550.

51. Pechmann, S. and Frydman, J. (2013) Evolutionary conservation of codon optimality reveals hidden signatures of cotranslational folding. Nat Struct Mol Biol, 20, 237–243.

52. Polte, C., Wedemeyer, D., Oliver, K.E., Wagner, J., Bijvelds, M.J.C., Mahoney, J., de Jonge, H.R., Sorscher, E.J. and Ignatova, Z. (2019) Assessing cell-specific effects of genetic variations using tRNA microarrays. BMC Genomics, 20, 549.

53. Koplin, A., Preissler, S., Ilina, Y., Koch, M., Scior, A., Erhardt, M. and Deuerling, E. (2010) A dual function for chaperones SSB-RAC and the NAC nascent polypeptide-associated complex on ribosomes. J Cell Biol, 189, 57–68.

54. Namane, A. and Saveanu, C. (2022) Composition and Dynamics of Protein Complexes Measured by Quantitative Mass Spectrometry of Affinity-Purified Samples. Methods Mol Biol, 2477, 225–236.

55. Kall, L., Storey, J.D. and Noble, W.S. (2008) Non-parametric estimation of posterior error probabilities associated with peptides identified by tandem mass spectrometry. Bioinformatics, 24, i42–48.

56. Nesvizhskii, A.I., Keller, A., Kolker, E. and Aebersold, R. (2003) A statistical model for identifying proteins by tandem mass spectrometry. Anal Chem, 75, 4646–4658.

57. Ritchie, M.E., Phipson, B., Wu, D., Hu, Y., Law, C.W., Shi, W. and Smyth, G.K. (2015) limma powers differential expression analyses for RNA-sequencing and microarray studies. Nucleic Acids Res, 43, e47.

58. Preissler, S., Reuther, J., Koch, M., Scior, A., Bruderek, M., Frickey, T. and Deuerling, E. (2015) Not4-dependent translational repression is important for cellular protein homeostasis in yeast. EMBO J, 34, 1905–1924.

59. Truman, A.W., Kristjansdottir, K., Wolfgeher, D., Ricco, N., Mayampurath, A., Volchenboum, S.L., Clotet, J. and Kron, S.J. (2015) Quantitative proteomics of the yeast Hsp70/Hsp90 interactomes during DNA damage reveal chaperone-dependent regulation of ribonucleotide reductase. J Proteomics, 112, 285–300.

60. Schuller, A.P., Wu, C.C., Dever, T.E., Buskirk, A.R. and Green, R. (2017) eIF5A Functions Globally in Translation Elongation and Termination. Mol Cell, 66, 194–205 e195.

61. Wada, M. and Ito, K. (2023) The CGA codon decoding through tRNA(Arg) (ICG) supply governed by Tad2/Tad3 in Saccharomyces cerevisiae. FEBS J, 290, 3480–3489.

62. Davyt, M., Bharti, N. and Ignatova, Z. (2023) Effect of mRNA/tRNA mutations on translation speed: Implications for human diseases. J Biol Chem, 299, 105089.

63. Pavlov, M.Y., Ullman, G., Ignatova, Z. and Ehrenberg, M. (2021) Estimation of peptide elongation times from ribosome profiling spectra. Nucleic Acids Res, 49, 5124–5142.

64. Delbecq, P., Werner, M., Feller, A., Filipkowski, R.K., Messenguy, F. and Pierard, A. (1994) A segment of mRNA encoding the leader peptide of the CPA1 gene confers repression by arginine on a heterologous yeast gene transcript. Mol Cell Biol, 14, 2378–2390.

65. Araiso, Y., Imai, K. and Endo, T. (2022) Role of the TOM Complex in Protein Import into Mitochondria: Structural Views. Annu Rev Biochem, 91, 679–703.

66. Klass, D.M., Scheibe, M., Butter, F., Hogan, G.J., Mann, M. and Brown, P.O. (2013) Quantitative proteomic analysis reveals concurrent RNA-protein interactions and identifies new RNA-binding proteins in Saccharomyces cerevisiae. Genome Res, 23, 1028–1038.

67. Bhondeley, M. and Liu, Z. (2020) Mitochondrial Biogenesis Is Positively Regulated by Casein Kinase I Hrr25 Through Phosphorylation of Puf3 in Saccharomyces cerevisiae. Genetics, 215, 463–482.

68. Hayashi, S., Iwamoto, K. and Yoshihisa, T. (2023) A non-canonical Puf3p-binding sequence regulates CAT5/COQ7 mRNA under both fermentable and respiratory conditions in budding yeast. PLoS One, 18, e0295659.

69. van de Poll, F., Sutter, B.M., Acoba, M.G., Caballero, D., Jahangiri, S., Yang, Y.S., Lee, C.D. and Tu, B.P. (2023) Pbp1 associates with Puf3 and promotes translation of its target mRNAs involved in mitochondrial biogenesis. PLoS Genet, 19, e1010774.

